# Optimizing network neuroscience computation of individual differences in human spontaneous brain activity for test-retest reliability

**DOI:** 10.1101/2021.05.06.442886

**Authors:** Chao Jiang, Ye He, Richard F. Betzel, Yin-Shan Wang, Xiu-Xia Xing, Xi-Nian Zuo

**Affiliations:** School of Psychology, Capital Normal University, Beijing, China; School of Artificial Intelligence, Beijing University of Posts and Telecommunications, Beijing, China; Department of Psychological and Brain Sciences, Indiana University, Bloomington, Indiana, United States; State Key Laboratory of Cognitive Neuroscience and Learning, Beijing Normal University, Beijing, China; Developmental Population Neuroscience Research Center, International Data Group/McGovern Institute for Brain Research, Beijing Normal University, Beijing, China; Department of Applied Mathematics, College of Mathematics, Faculty of Science, Beijing University of Technology, Beijing, China; National Basic Science Data Center, Beijing, China; Institute of Psychology, Chinese Academy of Sciences, Beijing, China

**Keywords:** individual difference, reliability, open science, spontaneous brain activity, connectome

## Abstract

A rapidly emerging application of network neuroscience in neuroimaging studies has provided useful tools to understand individual differences in intrinsic brain function by mapping spontaneous brain activity, namely intrinsic functional network neuroscience (ifNN). However, the variability of methodologies applied across the ifNN studies - with respect to node definition, edge construction, and graph measurements-makes it difficult to directly compare findings and also challenging for end users to select the optimal strategies for mapping individual differences in brain networks. Here, we aim to provide a benchmark for best ifNN practices by systematically comparing the measurement reliability of individual differences under different ifNN analytical strategies using the test-retest design of the Human Connectome Project. The results uncovered four essential principles to guide ifNN studies: 1) use a whole brain parcellation to define network nodes, including subcortical and cerebellar regions, 2) construct functional networks using spontaneous brain activity in multiple slow bands, 3) optimize topological economy of networks at individual level, 4) characterise information flow with specific metrics of integration and segregation. We built an interactive online resource of reliability assessments for future ifNN (ibraindata.com/research/ifNN).

**AUTHOR SUMMARY:** It is an essential mission for neuroscience to understand the individual differences in brain function. Graph or network theory offer novel methods of network neuroscience to address such a challenge. This article documents optimal strategies on the test-retest reliability of measuring individual differences in intrinsic brain networks of spontaneous activity. The analytical pipelines are identified to optimize for highly reliable, individualized network measurements. These pipelines optimize network metrics for high inter-individual variances and low inner-individual variances by defining network nodes with whole-brain parcellations, deriving the connectivity with spontaneous high-frequency slow-band oscillations, constructing brain graphs with topology-based methods for edge filtering, and favoring multi-level or multi-modal metrics. These psychometric findings are critical for translating the functional network neuroscience into clinical or other personalized practices requiring neuroimaging markers.

## INTRODUCTION

Over the past two decades, network neuroscience has helped transform the field of neuroscience (D. Bassett et al., 2020), providing a quantitative methodology framework for modeling brains as graphs (or networks) composed of nodes (brain regions) and edges (their connections), namely connectomics (Sporns, 2013a). The organization and topology of macro-scale brain networks can be characterized by a growing suite of connectomic measurements including efficiency, centrality, clustering, small-word topology, rich-club, etc (Craddock et al., 2013). In parallel, resting-state fMRI (rfMRI) has opened up new avenues towards understanding the intrinsic human brain function (Biswal et al., 2010). In conjunction with network neuroscience, rfMRI has led to the emergence of a multidisciplinary field, intrinsic functional connectomics or network neuroscience (ifNN), in which the brain’s intrinsic, interregional connectivity is estimated from rfMRI recordings. It has been widely used to investigate the system-level organization of the human brain function and its relationship with individual differences (Dubois & Adolphs, 2016) in developmental (Zuo et al., 2017), socio-cultural (Pessoa, 2018) and clinical conditions (Fornito, Zalesky, & Breakspear, 2015).

Highly reliable measurements are essential for studying individual differences. In general, reliability characterises a proportion of measurement variability between different subjects relative to the overall variability including both between-subject and within-subject (i.e., random) components (Xing & Zuo, 2018). It is commonly used to assess the consistency or agreement between measurements, or the ability to obtain consistent measures over time. Beyond that, it can also serve as a measure of discriminablity (Xing & Zuo, 2018; Zuo, Biswal, & Poldrack, 2019; Zuo, Xu, & Milham, 2019). For example, if a measurement can more sufficiently capture individual characteristics (i.e., better differentiate a group of individuals), it will produce higher between-subject variability and thus higher reliability than a measurement underestimating the between-subject variability. Such reliability concept has well-established statistical theory and applications in fields such as psychology (Elliott, Knodt, Caspi, Moffitt, & Hariri, 2021) and medicine (Kraemer, 2014) where it is used in psychometric theory and diagnosis theory, respectively. Specifically, in psychology, reliability is important for assessing the validity of psychological tests, and in medicine, it is important for accurately diagnosing and treating patients. In the field of human brain mapping, more recent studies have demonstrated that the measurement reliability is equivalent to the “fingerprint” or discriminability of the measurement under the Gaussian distribution (Bridgeford et al., 2021; Milham, Vogelstein, & Xu, 2021). Therefore, the optimization of measurement reliability of the individual differences can help guide ifNN processing and analysis pipelines for individualized or personalized (e.g., neurodevelopmental (Herting, Gautam, Chen, Mezher, & Vetter, 2018) or clinical (Matthews & Hampshire, 2016)) research.

Previous studies have demonstrated that many functional network measurements with rfMRI have limited reliability (Noble, Scheinost, & Constable, 2019; Zuo & Xing, 2014). These low levels of reliability could be an indication of failure in handling individual variability at different levels (Elliott, Knodt, & Hariri, 2021; Hallquist & Hillary, 2019). In particular, experimental design and processing decisions related to scan duration, determining frequency range, and regressing global signal have impacts on rfMRI measurements and thus their reliability (Noble et al., 2019; Zuo et al., 2013). Although less focused on reliability, existing network neuroscience studies revealed that their findings are influenced by choices of parcellation templates (Bryce et al., 2021; Wang et al., 2009), edge construction and definition, and choice of graph metrics (Liang et al., 2012). How these decisions affect the measurement reliability in ifNN deserves further investigation. These analytical choices have been implemented in different software packages but can vary from one package to another, and thus introduce more analytic variability (Botvinik-Nezer et al., 2020). Beyond limited examinations on reliability (Aurich, Filho, da Silva, & Franco, 2015; Braun et al., 2012; Termenon, Jaillard, Delon-Martin, & Achard, 2016), a systematic investigation into the measurement reliability is warranted to guide ifNN software use and analyses.

We conducted a systematic ifNN reliability analysis using the test-retest rfMRI data from the Human Connectome Project (HCP). The HCP has developed its imaging acquisition and data pre-processing (Glasser et al., 2013) by integrating various strategies optimized for reliability in previous studies (Noble et al., 2019; Noble, Scheinost, & Constable, 2021; Zuo & Xing, 2014; Zuo et al., 2013). We thus analyzed the minimally pre-processed HCP rfMRI data and focused our work on four key post-analytic stages: node definition, edge construction, network measurement, and reliability assessments. In the end, we propose a set of principles to guide researchers in performing reliable ifNN, advancing the field-standard call for the best practices in network neuroscience. We released all the codes and reliability data by building an online platform for sharing the data and computational resources to foster future ifNN.

## MATERIALS AND METHODS

A typical analysis pipeline in ifNN includes steps for node definition (parcellations) and edge construction (frequency bands, connectivity estimation and filtering schemes) (Fig. 1a). To determine an optimal pipeline, we combine the most reliable strategies across different parts of the analysis by comparing the reliability of derived global network metrics. The HCP test-retest data were employed for reliability evaluation (Fig. 1b) using the intraclass correlation (ICC) statistics on the measurement reliability. Overall reliability assessments associated with the various analytic strategies as well as their impact on between- and within-subject variability (Fig. 1c) are investigated. We calculated the between-subject variability (*V_b_*) and within-subject variability (*V_w_*) and normalized them to values between 0 and 1 by the total sample variances. The changes in these variability measures, Δ*V_b_* and Δ*V_w_*, were used to create a reliability gradient represented by a vector. The length of the arrow reflects the amplitude of the change in reliability when comparing one choice (pink circle, *J*) to another choice (red circle, *K*). The direction of the arrow, **JK**, indicates the sources of the change in reliability. In this case, the reliability increases from a moderate to a substantial level with an increase in between-subject variability (Δ*V_b_* > 0) and a decrease in within-subject variability (Δ*V_w_* < 0). We then determine the optimized pipelines based on the highest reliability measurements, while documenting the derived both global and local network metrics and both their reliability and variability at an individual level.

**Figure 1.**
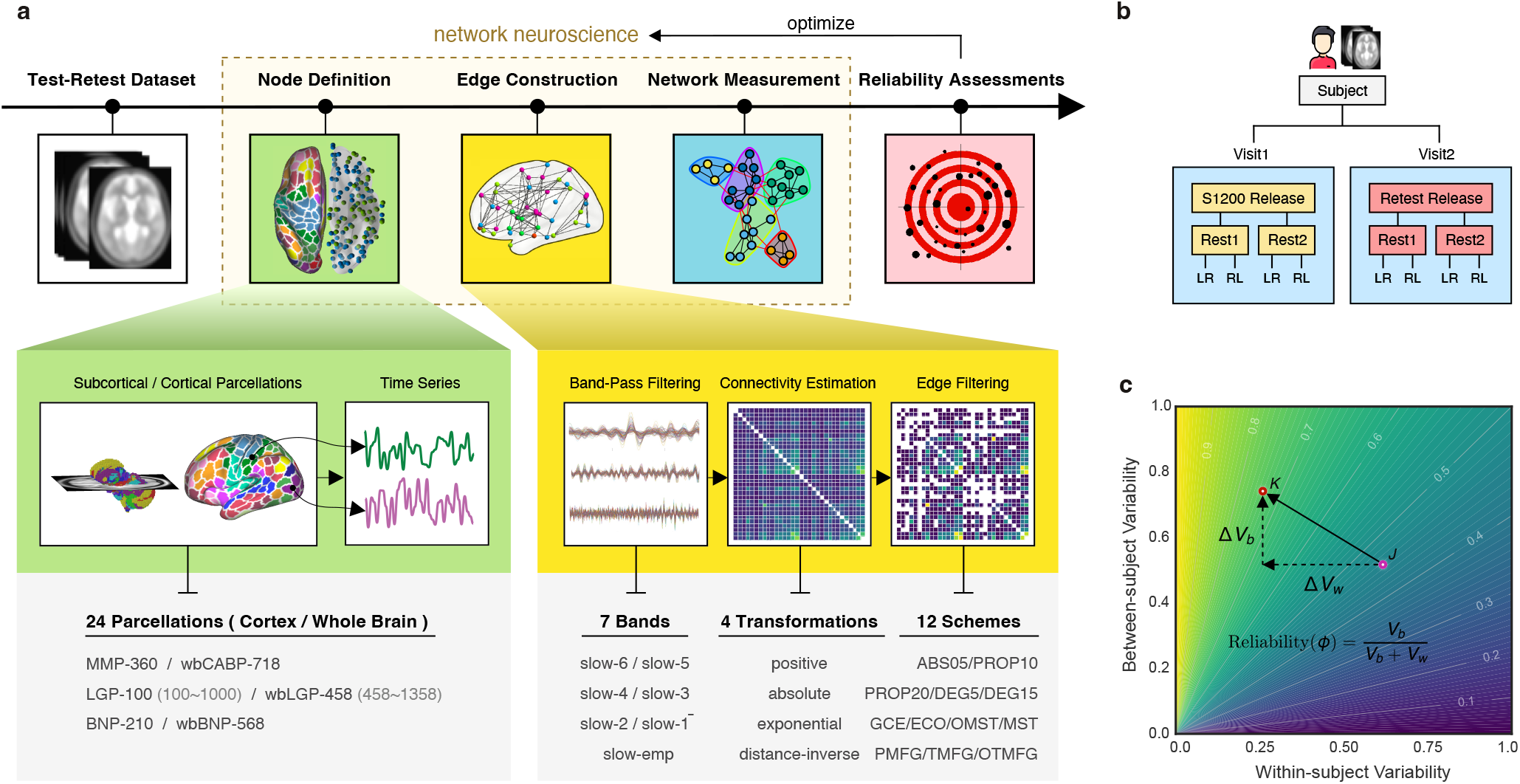
Analytical pipelines for reliable ifNN. a) There are five stages during our analyses: (1) test-retest dataset (white box) downloaded from HCP website, (2) node definition (green box) defining nodes using a set of brain areas of 24 different partitions of the human brain, (3) edge construction (yellow box) estimating individual correlation matrices using the six frequency bands (slow 1-6) from Buzsaki’s theoretical framework on the brain oscillations as well as the widely used empirical frequency band (Slow-emp) and transferring these matrices into adjacency matrices using 7 × 4 × 12 different strategies on edge construction including band-pass filtering, connectivity estimation and edge filtering, (4) network analysis (blue box) systematically calculating various brain graph metrics on measurements of information flow, and (5) reliability assessment (red box) evaluating test-retest reliability with massive linear mixed models. b) The test-retest data shared multimodal MRI datasets of 46 subjects in the HCP S1200 release and the HCP Retest release. Each subject underwent the first four test scans on two days (two scans per day: Rest1 and Rest2) and return several months later to finish the four retest scans on another two days. c) Measurement reliability refers to the inter-individual or between-subject variability *V_b_* relative to the intra-individual or within-subject variability *V_w_*. Variability of both between-subject (*V_b_*) and within-subject (*V_w_*) are normalized into between 0 and 1 by the total sample variances. Their changes (Δ*V_b_* and Δ*V_w_*) introduce a reliability gradient as represented by the vector (the black arrow). The length of the arrow reflects the amplitude of reliability changes when the reliability assessment from one choice (pink circle, *J*) to another choice (red circle, *K*). Further, the arrow’s direction (**JK**) indicates the sources of this reliability change. Here the reliability becomes from moderate to substantial level with increases of between-subject variability (Δ*V_b_* > 0) and decreases of within-subject variability (Δ*V_w_* < 0).

Specifically, using the HCP test-retest dataset, our analytic procedure implemented the four post-analytic stages: node definition, edge construction, network measurement and reliability assessments. The test-retest rfMRI dataset underwent the standardized preprocessing pipeline developed by the HCP team (Glasser et al., 2013). The second step defines nodes (green box) using sets of brain areas based on 24 partitions, and then extracts the nodal time series. During the third step (yellow box), individual correlation matrices are first estimated based upon the six frequency bands derived from Buzsaki’s theoretical framework on brain oscillations (Buzsaki & Draguhn, 2004) along with the classical band widely used (0.01 - 0.08 Hz). These matrices are then converted into adjacency matrices using 4 × 12 = 48 strategies on edge filtering. In the fourth step, we performed graph analyses (blue box) by systematically calculating the brain graph metrics at global, modular and nodal scales. Finally, test-retest reliability was evaluated (red box) as ICCs with the linear mixed models. We present details of these analyses in the following sections.

### Test-Retest Dataset

The WU-Minn Consortium in HCP shared a set of test-retest multimodal MRI datasets of 46 subjects from both the S1200 release and the Retest release. These subjects were retested using the full HCP 3T multimodal imaging and behavioral protocol. Each subject underwent the four scans on two days (two scans per day: Rest1 versus Rest2) during the first visit and returned several months later to finish the four scans on another two days during the second visit (Fig. 1b). The test-retest interval ranged from 18 to 328 days (mean: 4.74 months, standard deviation: 2.12 months). Only 41 subjects (28 females, age range: 26-35 years; 13 males, age range: 22-33 years) had full length rfMRI data across all the eight scans, 2 visits × 2 days × 2 (LR and RL ecoding directions), and were included in the subsequent analyses. Then we averaged across the RL and LR encodings for each day, so each subject had 4 repeated measurements in the ICC estimation. This sample size is larger that the minimal sample size (*N =* 35) for fair reliability with 80% power and significance level of 0.05 based on the above mentioned test-retest design (4 observations per subject) (Bujang & Baharum, 2017). The HCP rfMRI protocols for scanning and preprocessing images have been optimized for reliability.

During the scanning, participants were instructed to keep their eyes open and to let their mind wander while fixating on a cross-hair projected on a dark background. Data were collected at the 3T Siemens Connectome Skyra MRI scanner with a 32-channel head coil. All functional images were acquired using a multiband gradient-echo EPI imaging sequence (2mm isotropic voxel, 72 axial slices, TR = 720ms, TE = 33.1ms, flip angle = 52°, field of view = 208 × 180 mm^2^, matrix size = 104 × 90 and a multiband factor of 8). A total of 1200 images was acquired for a duration of 14 min and 24 s. Details on the imaging protocols can be found in (Smith et al., 2013).

The protocols of rfMRI image preprocessing and artifact-removal procedures are documented in detail elsewhere and generated the minimally preprocessed HCP rfMRI images. Artifacts were removed using the ICA-based X-noiseifier (ICA + FIX) procedure, followed by MS-MAll for inter-subject registration. The preprocessed rfMRI data were represented as a time series of grayordinates (4D), combining both cortical surface vertices and subcortical voxels (Glasser et al., 2013).

### Node Definition

A brain graph defines a node as a brain area, which is generally derived by an element of brain parcellation (parcel) according to borders or landmarks of brain anatomy, structure or function as well as an element of volume (voxel) in imaging signal acquisition or a cluster of voxels (Sporns, 2013b). Due to the high computational demand of voxel-based brain graph, in this study we defined nodes as parcels according to the following brain parcellation strategies (Fig. 2a). A surface-based approach has been demonstrated to outperform other approaches for fMRI analysis (Coalson, Van Essen, & Glasser, 2018; Zuo et al., 2013) and thus the nodes are defined in the surface space (total 24 surface parcellation choices). Of note, we adopted a naming convention for brain parcellations as follows: *‘ParcAbbr-NumberOfParcels’* (e.g., LGP-100 or its whole-brain version wbLGP-458).

**Figure 2.**
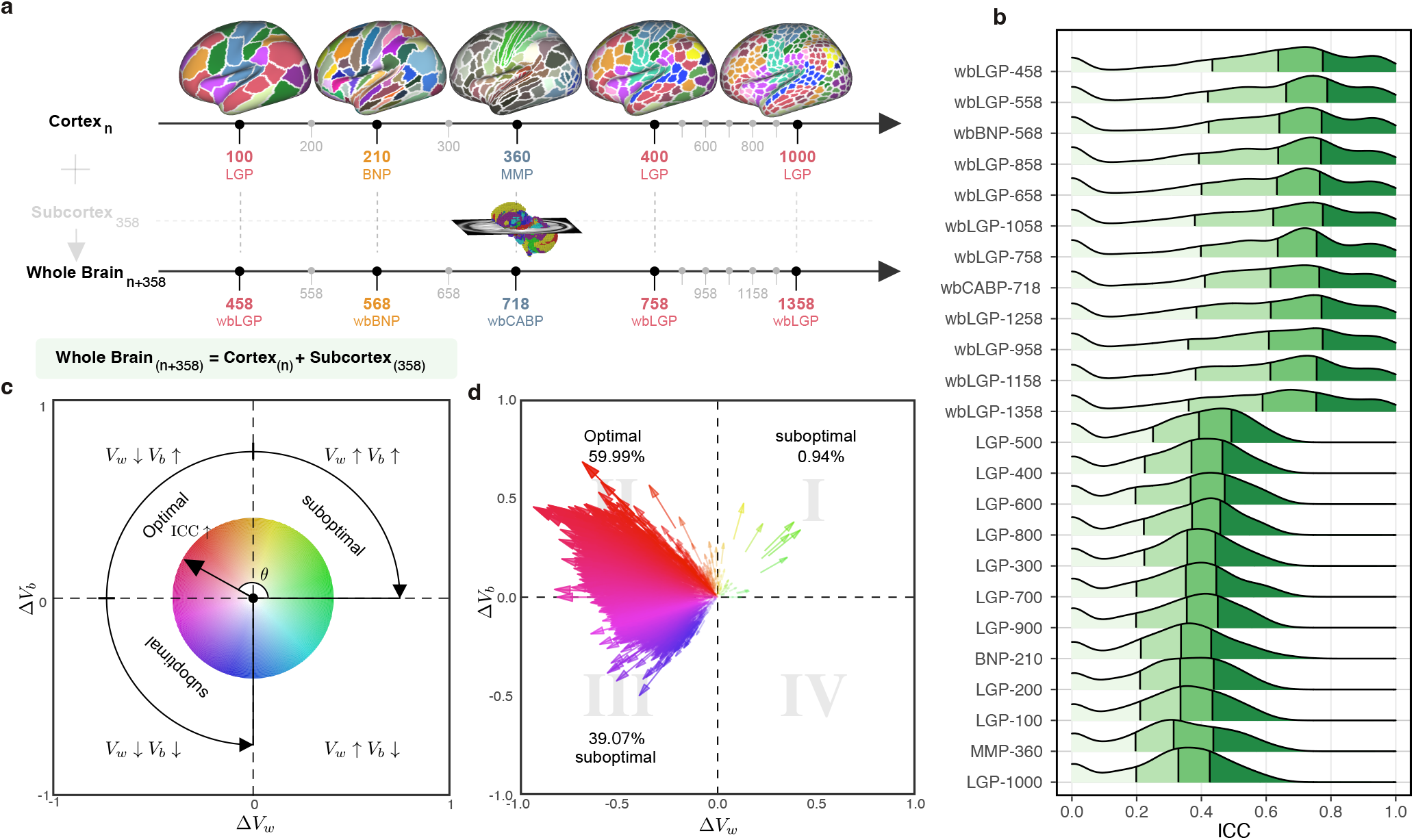
Parcellation choices impact measurement reliability and individual variability. a) Node definitions are derived from the process of spatially partitioning the human cortex and whole brain (including both cortical and subcortical nodes) at various resolutions, see more details of these name abbreviations in **Methods**. b) Density plots are visualized for distributions of the ICCs under the various parcellation choices on node definition. These density distributions are ranked from top to bottom according to decreases of the mean ICCs while the four colors depict the four quantiles. c) Reliability gradient between any one whole-brain parcellation choice and its corresponding cortical parcellation choice is decomposed into the axis of changes of the between-subject variability (Δ*V_b_*) and the axis of changes of the within-subject variability (Δ*V_w_*). This gradient can be represented as an vector, which is the black arrow from the origin with an angle *θ* with the *x*-axis while the color encodes this angle and the transparency or the length reflects the magnitude of the degree of ICC improvement. According to the anatomy of reliability, the optimal space is in the second quadrant (quadII) while the first and third quadrant (quadI and quadIII) are suboptimal for reliability. d) The improvement in the reliability of the pipeline, which is defined from the cortical parcellations to the corresponding whole-brain parcellations (including the subcortex), is illustrated by gradient arrows in the plane of individual variability, while controlling for all other processing steps. Each arrow represents a specific global metric, while controlling for all other processing steps. The position of the arrows reflects the magnitude of between- and within-subject variability changes (Δ*V_b_*, Δ*V_w_*), and the size of the arrows indicates the magnitude of ICC changes.

#### HCP Multi-Modal Parcellation (MMP)

A cortical parcellation generated from multi-modal images of 210 adults from the HCP database, using a semi-automated approach (Glasser et al., 2016). Cortical regions are delineated with respect to their function, connectivity, cortical architecture, and topography, as well as, expert knowledge and meta-analysis results from the literature (Glasser et al., 2016). The atlas contains 180 parcels for each hemisphere.

#### Local-Global Parcellation (LGP)

A gradient-weighted Markov Random Field model integrating local gradient and global similarity approaches produces the novel parcellations (Schaefer et al., 2018). The final version of LGP comes with a multi-scale cortical atlas including 100, 200, 300, 400, 500, 600, 700, 800, 900, and 1000 parcels (equal numbers across the two hemispheres). One benefit of using LGP is to have nodes with almost the same size, and these nodes are also assigned to the common large-scale functional networks (Thomas Yeo et al., 2011).

#### Brainnetome Parcellation (BNP)

Both anatomical landmarks and connectivity-driven information are employed to develop this volumetric brain parcellation (Fan et al., 2016). Specifically, anatomical regions defined as in (Desikan et al., 2006) are parcellated into subregions using functional and structural connectivity fingerprints from HCP datasets. Cortical parcels are obtained by projecting their volume space to surface space. It is noticed that the original BNP contains both cortical (105 areas per hemisphere) and subcortical (36 areas) regions but only the 210 cortical parcels are included for the subsequent analyses.

#### Whole-Brain Parcellation (wb)

Inclusion of subcortical areas has been shown unignorable influences on brain graph analyses (D. Greene et al., 2020; Noble et al., 2019), and we thus also constructed brain graphs with subcortical structures in volume space as nodes by adding these nodes to the cortical brain graphs. To get a high-resolution subcortical parcellation, we adopted the 358 subcortical parcels in (Ji et al., 2019). The authors employed data of 337 unrelated HCP healthy volunteers and extended the MMP cortical network partition into subcortex. This results a set of whole-brain parcellations by combining these subcortical parcels with the aforementioned cortical parcellations, namely **wbMMP**,**wbLGP** and **wbBNP**. We noticed that the wbMMP-718 has been named by the authors of (Ji et al., 2019) as the Cole-Anticevic Brain-wide Network Partition, and we thus renamed the wbMMP-718 as wbCABP-718 for consistency.

### Edge Construction

After defining the node with each parcellation, in each parcel, regional mean time series were estimated by averaging the vertex time series at each time point. To construct an edge between a pair of nodes, their representative time series entered into the following steps in order: *band-pass filtering, inter-node connectivity transformation*, and *edge filtering*.

#### Band-Pass Filtering

Resting-state functional connectivity studies have typically focused on fluctuations below 0.08 Hz or 0.1 Hz (Biswal, Zerrin Yetkin, Haughton, & Hyde, 1995; Fox & Raichle, 2007), and assumed that only these frequencies contribute significantly to inter-regional functional connectivity (FC) while other frequencies are artifacts (Cordes et al., 2001). In contrast, however, other studies have found that specific frequency bands of the rfMRI oscillations make unique and neurobiologically meaningful contributions to resting-state functional connectivity (Salvador et al., 2005; Zuo & Xing, 2014). More recently, with fast fMRI methods, some meaningful FC patterns were reported across much higher frequency bands (Boubela et al., 2013). These observations motivate exploring a range of frequency bands beyond those typically studied in resting-state functional connectivity studies.

Buzsaki and Draguhn (Buzsaki & Draguhn, 2004) proposed a hierarchical organization of frequency bands driven by the natural logarithm linear law. This offers a theoretical template for partitioning rfMRI frequency content into multiple bands (Fig. 3a). The frequencies occupied by these bands have a relatively constant relationship to each other on a natural logarithmic scale and have a constant ratio between any given pair of neighboring frequencies (Buzsáki, 2009). These different oscillations are linked to different neural activities, including cognition, emotion regulation, and memory (Achard, Salvador, Whitcher, Suckling, & Bullmore, 2006; Buzsáki, 2009; Fox & Raichle, 2007). Advanced by the fast imaging protocols offered by the HCP scanner, the short scan interval (TR = 720ms) allows us to obtain more oscillation classes that the traditional rfMRI method. We incorporate the Buzsaki’s framework (Buzsaki & Draguhn, 2004; Penttonen & Buzsáki, 2003) with the HCP fast-TR datasets by using the DREAM toolbox (Gong et al., 2021) in the Connectome Computation System (Xing, Xu, Jiang, Wang, & Zuo, 2022; Xu, Yang, Jiang, Xing, & Zuo, 2015). It decomposed the time series into the six slow bands as illustrated in Fig. 3a.

**Figure 3.**
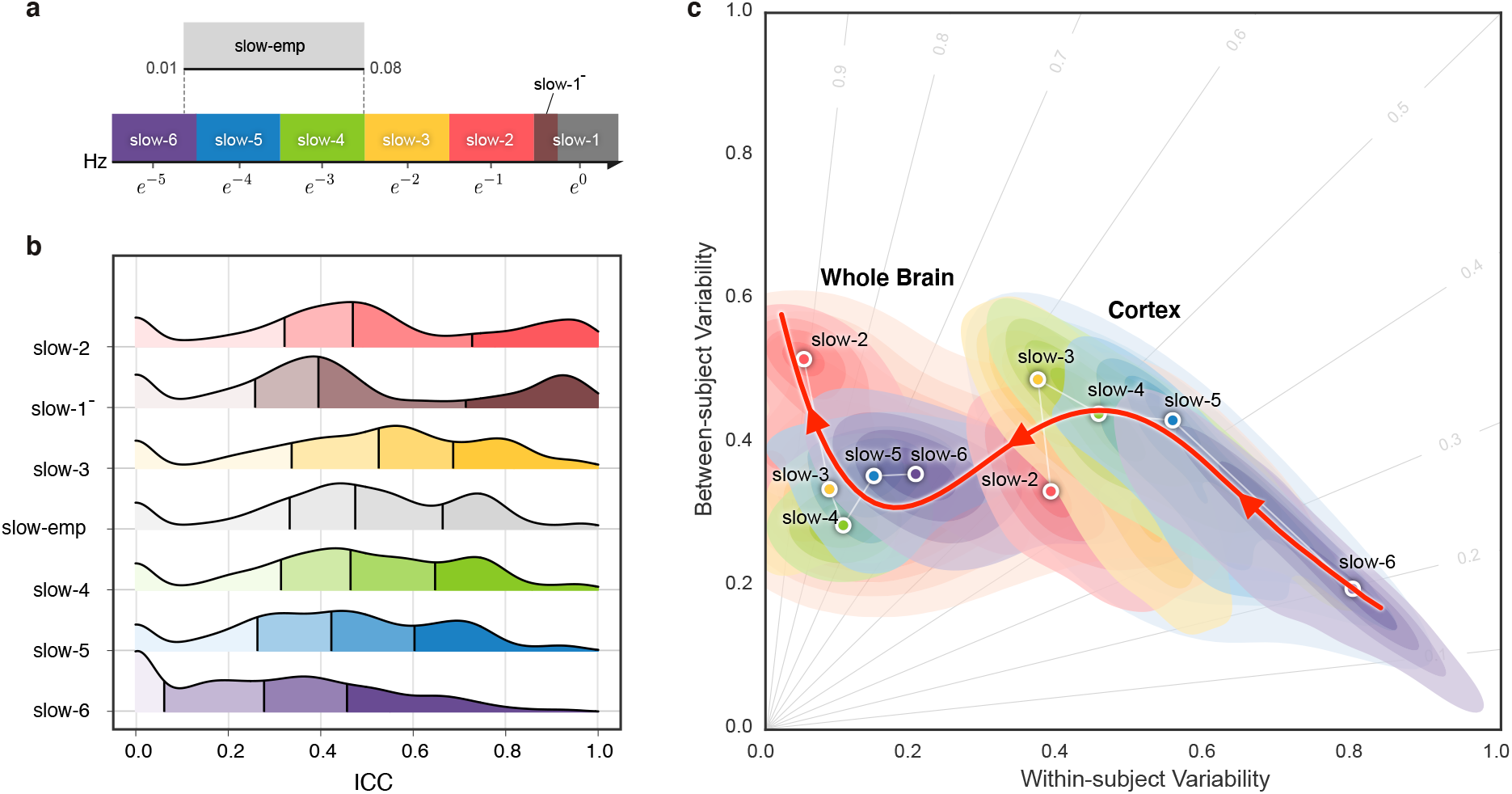
Reliability gradient across the slow bands and changes of related individual variability. a) Classes of frequency bands for slow oscillations derived from the natural logarithm linear law. b) Density plots are visualized for the ICC distributions under the various frequency bands. These density distributions are ranked from top to bottom according to decreases of the mean ICCs while the vertical lines depict the four quartiles. c) Network measurements are projected onto the reliability anatomy plane coordinated by both between- and within-subject variability. These dot plots are fitted into the topographic (contour) maps where the local maxima for each band is labeled as a circle. To highlight the trend of increasing reliability as the frequency band increases, a fourth-order polynomial curve (represented by a red line) is fitted to the frequency contour plot peak points, tracing the reliability flow along slow-to-fast oscillations in the cortex and whole brain.

#### Connectivity Transformation

For each scan, individual nodal representative time series were band-pass filtered with each of the six frequency bands, and another empirical frequency band, slow-emp (0.01-0.08Hz). The Pearson’s correlation *r_ij_* ∈ [-1, 1] between the filtered time series of each pair of nodes *i* = 1,…, *N, j* = 1,…, *N* was calculated (*N* is the number of nodes). These correlation values provided an estimation on the edge strengths between the two nodes, and formed a N × N symmetric correlation matrix *R* = (*r_ij_*) for each given subject, scan, parcellation, and frequency band.

Many network metrics are not well defined for negatively weighted connections. In order to ensure that the connection weights are positive only, we applied four types of transformations to the symmetric correlation matrix: the **positive** (Eq.pos), **absolute** (Eq.abs), **exponential** (Eq.exp) and **distance-inverse** (Eq.div) functions, respectively. This avoids the negative values in the inter-node connectivity matrix *W* = (*w_ij_*) where *z_ij_* = tanh^-1^ (*r_ij_*) is Fisher’s *z*–transformation.

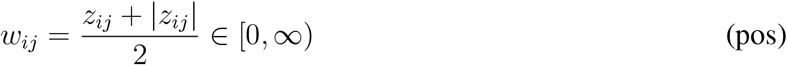

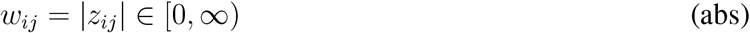

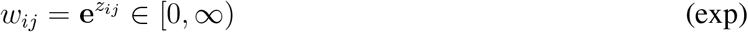

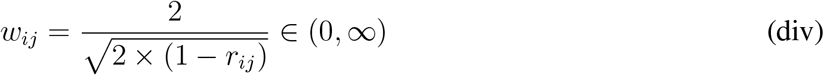

The connectivity matrix represents a set of the node parcels and relational quantities between each pair of the nodes, and will serve as the basis of following edge filtering procedure for generation of the final brain graphs.

#### Edge Filtering

In a graph, edges represent a set of relevant interactions of crucial importance to obtain parsimonious descriptions of complex networks. Filtering valid edges can be highly challenging due to the lack of ‘ground truth’ of the human brain connectome. To provide a reliable way of building candidate edges, we sampled the following 12 schemes on edge filtering and applied them to the connectivity matrices.

##### Absolute Weight Thresholding (ABS)

This approach selects those edges that exceed a manually defined absolute threshold (e.g., correlations higher than 0.5), setting all correlations smaller than 0.5 to 0 (ABS_05_). This is a simple approach to reconstruct networks Hagmann et al. (2007).

##### Proportional Thresholding (PROP)

It is a common step in the reconstruction of functional brain networks to ensure equal edge density across subjects (D. Bassett et al., 2009; Rubinov, Sporns, van Leeuwen, & Breakspear, 2009; van den Heuvel et al., 2017). It keeps the number of connections fixed across all individuals to rule out the influence of network density on the computation and comparison of graph metrics across groups. This approach includes the selection of a fixed percentage of the strongest conncections as edges in each individual network or brain graph. Compared to ABS, PROP has been argued to reliably separate density from topological effects (Braun et al., 2012; Ginestet, Nichols, Bullmore, & Simmons, 2011) and to result in more stable network metrics (Garrison, Scheinost, Finn, Shen, & Constable, 2015). This makes it a commonly used approach for network construction and analysis in disease-related studies. Here, we focused on two threshholds that are commonly reported in the literature: 10% (PROP_10_) and 20% (PROP_20_).

##### Degree Thresholding (DEG)

The structure of a graph can be biased by the number of existing edges. Accordingly, statistical measures derived from the graph should be compared against graphs that preserve the same average degree, K. A threshold of the degree can be chosen to produce graphs with a fixed mean degree (e.g., *K* = 5, DEG_5_), which is the average nodal degrees of an individual graph from a single subject’s scan. Many network neuroscience studies have taken this choice for *K* = 5 (S. I. Dimitriadis, Laskaris, Del Rio-Portilla, & Koudounis, 2009; Micheloyannis et al., 2006; Milo et al., 2002; Stam, Jones, Nolte, Breakspear, & Scheltens, 2006). We also include the DEG_15_ for denser graphs of the brain networks.

##### Global Cost Efficiency Optimization (GCE)

Given a network with a cost *ρ*, its global efficiency is a function of the cost *E_g_*(*ρ*), and its GCE is *J*(*ρ*) = *E_g_*(*ρ*) – *ρ*. Several studies suggested that brain networks, in particular those with small-world topology, maximize their global-cost efficiency (D. S. Bassett et al., 2008), i.e., *J^max^* = max*_ρ_ J*(*ρ*). Computationally, this scheme is implemented by looping all network costs (e.g., adding edges with weights in order) to find the *J^max^* (see Fig. 2b) where the corresponding edge weight was determined as the threshold for edge filtering. In this sense, GCE is an individualised and optimised version of ABS, PROP and DEG while the latter three are commonly employed with a fixed threshold for all individuals.

##### Overall Efficiency Cost Optimization (ECO)

Both global and local efficiency are important graph features to characterize the structure of complex systems in terms of integration and segregation of information (Latora & Marchiori, 2001). ECO was proposed to determine a network density threshold for filtering out the weakest links (De Vico Fallani, Latora, & Chavez, 2017). It maximizes an extension of *J^max^*, the ratio between the overall (both global and local) efficiency and its wiring cost max*_ρ_ J^ext^*(*ρ*) = (*E_g_*(*ρ*) + *E_loc_*(*ρ*))/*ρ* where *E_loc_* denotes the network local efficiency. The study (Latora & Marchiori, 2001) also demonstrated that, to maximize *J*, these networks have to be sparse with an average node degree *K* ≃ 3.

##### Minimum Spanning Tree (MST)

This is an increasingly popular method for identifying the smallest and most essential set of connections while ensuring that the network forms a fully connected graph (Guo, Qin, Chen, Xu, & Xiang, 2017; Meier, Tewarie, & Van Mieghem, 2015; Otte et al., 2015; van Nieuwenhuizen et al., 2018). The tenet of using MST is to summarize information and index structure of the graph, and thus remove edges with redundant information (Mantegna, 1999). Specifically, an MST filtered graph will contain N nodes connected *via N* – 1 connections with minimal cost and no loops. This addresses key issues in existing topology filtering schemes that rely on arbitrary and user-specified absolute thresholds or densities.

##### Orthogonal Minimum Spanning Tree (OMST)

This topological filtering scheme was proposed recently (S. Dimitriadis, Antonakakis, Simos, Fletcher, & Papanicolaou, 2017) to maximize the information flow over the network *versus* the cost by selecting the connections via the OMSTs. It samples the full-weighted brain network over consecutive rounds of MST that are orthogonal to each other (see Fig. 2b). Practically, we extracted the 1st MST, and then we cleared their connections and we tracked the 2nd MST from the rest of the network connections, etc. Such an iterative procedure (stopped by the Mth MST) can get orthogonal MSTs and topologically filter brain network by optimizing the GCE under the constrains by the MST, leading to an integration of both GCE and MST

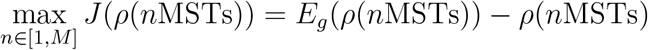

##### Planar Maximally Filtered Graph (PMFG)

The idea underneath PMFG (Tumminello, Aste, Di Matteo, & Mantegna, 2005) is to filter a dense matrix of weights by retaining the largest possible subgraph while imposing global constraints on the derived network topology. Edges with the strong connection weights are retained while constraining the subgraph to be a (spanning) tree globally. Similarly, during the PMFG construction, the largest weights are retained while constraining the subgraph to be a planar graph globally. The PMFG algorithm searches for the maximum weighted planar subgraph by adding edges one by one. The resulting matrix is sparse with 3(*N* – 2) edges. It starts by sorting all the edges of a dense matrix of weights in non-increasing order and tries to insert every edge in the PMFG. Edges that violate the planarity constraint are discarded.

##### Triangulated Maximally Filtered Graph (TMFG)

The algorithm for implementing PMFG is computationally expensive, and is therefore impractical when applied to large brain networks (Massara, Di Matteo, & Aste, 2016). A more efficient algorithms, TMFG, was developed that exhibited greatly reduced computational complexity compared to PMFG. This method captures the most relevant information between nodes by approximating the network connectivity matrix with the endorsement association matrix and minimizing spurious associations. The TMFG derived network contains 3-node (triangle) and 4-node (tetrahedron) cliques, imposing a nested hierarchy and automatically generates a chordal network (Massara et al., 2016; Song, Di Matteo, & Aste, 2012). Although TMFG is not widely applied in network neuroscience studies, it as been applied elsewhere and proven to be a suitable choice for modeling interrelationships between psychological constructs like personality traits (Christensen, Kenett, Aste, Silvia, & Kwapil, 2018).

##### Orthogonal TMF Graph (OTMFG)

To combine both the TMFG’s efficiency and OMST’s accuracy, we propose OTMFG to maximize the information flow over the network *versus* the cost by selecting the connections of the orthogonal TMFG. It samples the full-weighted brain network over consecutive rounds of TMFG that are orthogonal to each other.

In summary, as illustrated in Fig. 4a, the 12 edge filtering schemes transform a fully weighted matrix into a sparse matrix to represent the corresponding brain network. They can be categorized into two classes: threshold-based *versus* topology-based schemes. ABS_05_, PROP_10_, PROP_20_, DEG_5_, DEG_15_, ECO and GCE rely on a threshold for filtering and retaining edges with higher weights than the threshold. These schemes normally ignore the topological structure of the entire network and can result in isolated nodes. In contrast, the topology-based methods including MST, OMST, PMFG, TMFG and OTMFG, all consider the global network topology in determining which edges to retain. As illustrated in Fig. 4b, all the schemes are plotted in the *ρ* – *J^max^* plane for their network economics.

**Figure 4.**
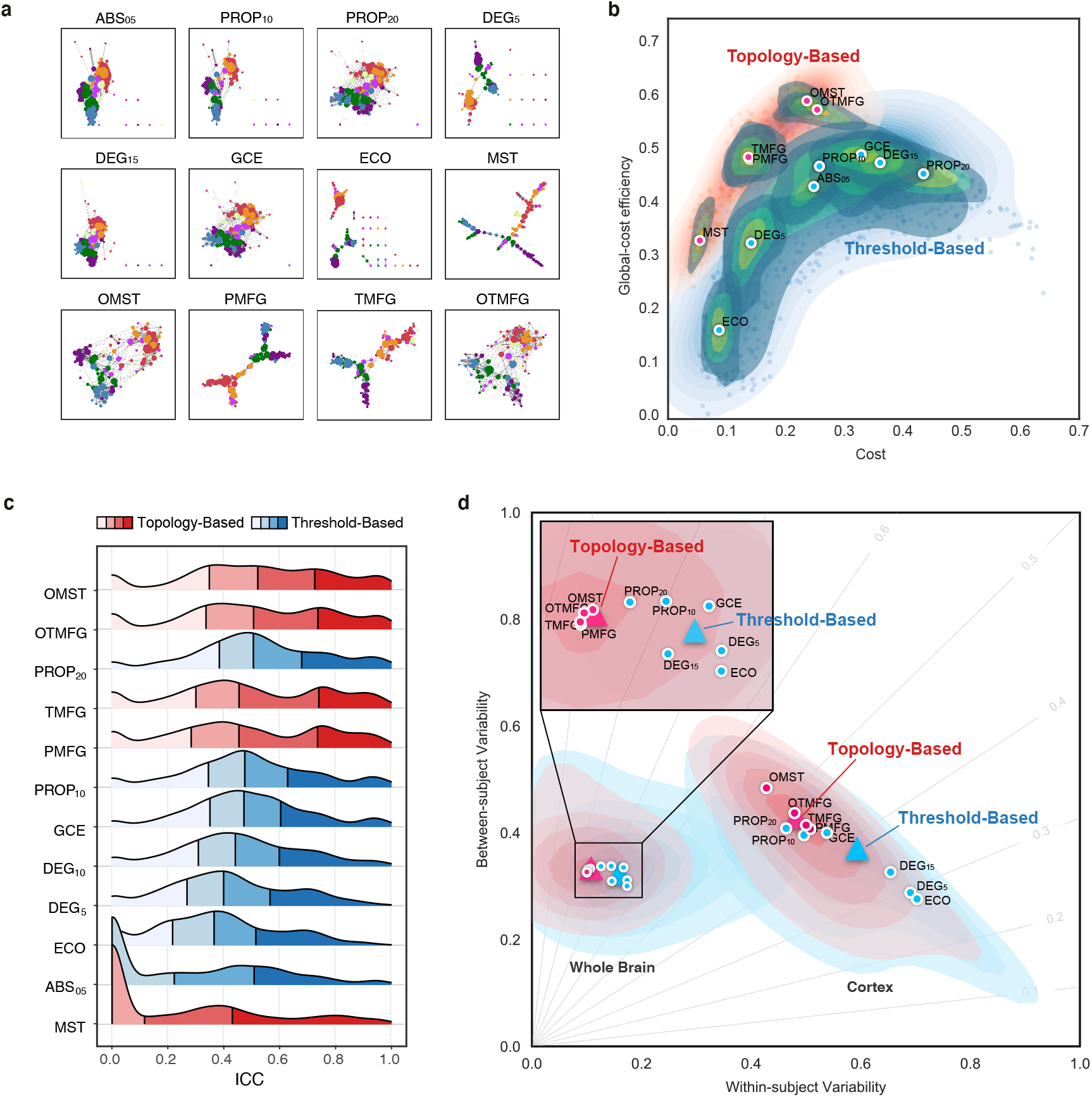
Edge filtering schemes and their networking performance. (a) Twelve schemes of filtering edge are applied to an individual connectivity matrix, resulting in the 12 brain networks with their nodes colored as the Yeo2011-7Networks (Thomas Yeo et al., 2011). (b) Global cost efficiency are plotted against network wiring costs of all the brain networks derived with the 12 edge filtering schemes from all the individual rfMRI scans. Red dots represent the topology-based while blue dots are for threshold-based networks. These dot plots are fitted into the topographic (contour) maps where the local maxima for each filtering choice is labeled as a circle. (c) Density plots are for ICC distributions under various the 12 edge filtering schemes. These density distributions are ranked from top to bottom according to decreases of the mean ICCs while the two colors depict the topology-based and threshold-based schemes. Four quartiles were indicated by vertical lines. (d) Network measurements are projected onto the reliability anatomy plane coordinated by both between- and within-subject variability. Red dots represent the topology-based while blue dots are for threshold-based networks. The topographic (contour) maps fit the dots and label the local maxima as a circle for each scheme and the global maxima as a triangle for the topology and threshold groups, respectively.

### Network Analysis

We performed graph-theory-driven network analysis by calculating several common graph-based metrics for the resulting graphs. These measures, broadly, can be interpreted based on whether the characterize the extent to which network structure allows for integrated or segregation information flow. Examples of integrative measures include average shortest path length (*L_p_*), global efficiency (*E_g_*), and pseudo diameter (D). Segregation measures include clustering coefficient (*C_p_*), local efficiency (*E_local_*), transitivity (*T_r_*), modularity (Q), and a suite of nodal centrality measures (Table 1). All the metrics are calculated using the Brain Connectivity Toolbox (Rubinov & Sporns, 2010). We employed **graph-tool** (https://graph-tool.skewed.de) and **NetworKit** (https://networkit.github.io) to achieve high performance comparable (both in memory usage and computation time) to that of a pure C/C++ library. We treated these metrics as the network measurements for subsequent reliability analysis.

**Table 1.**
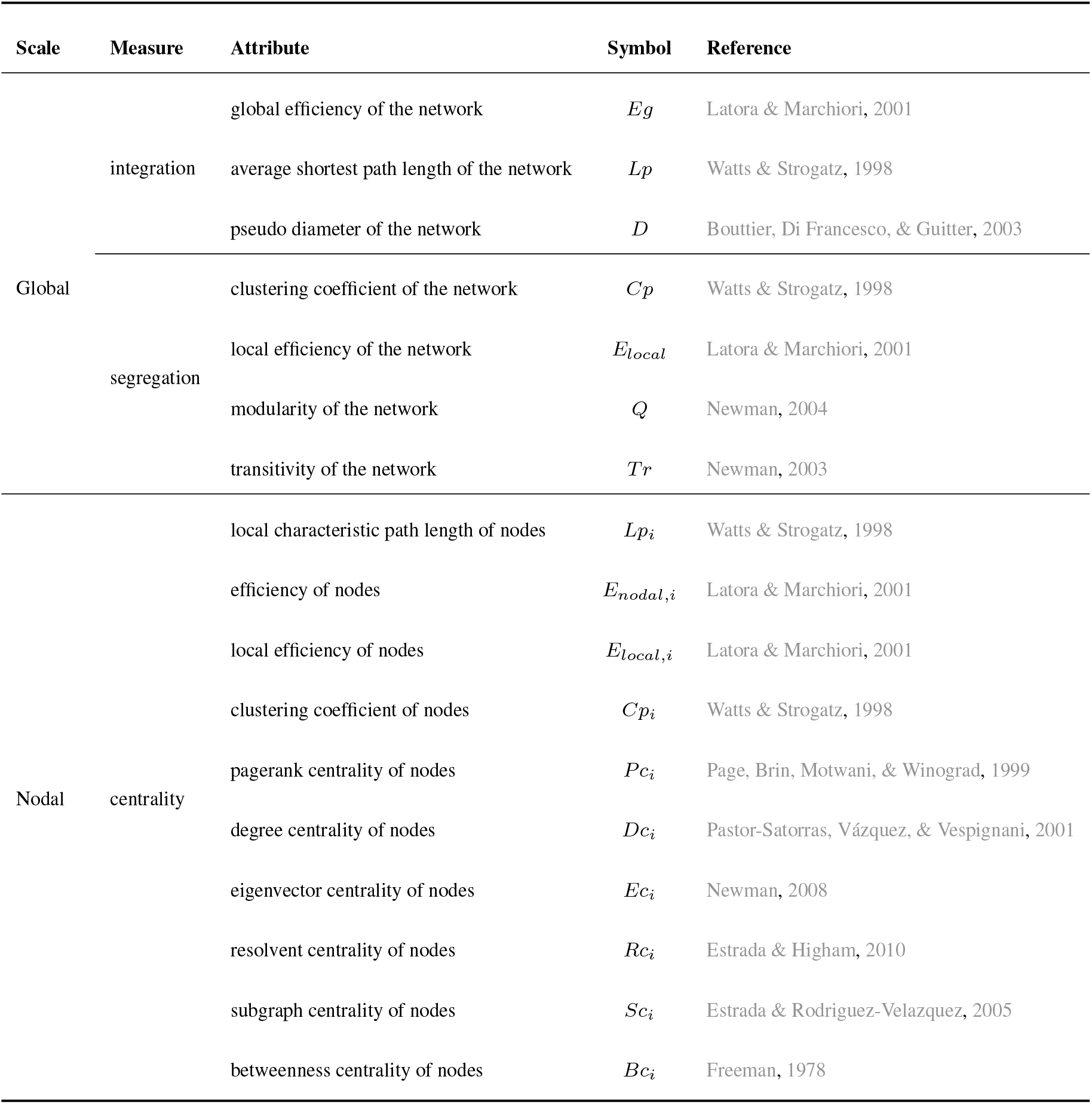
A list of the employed network metrics derived with graph theory

### Reliability Assessments

Measurement reliability is defined as the extent to which measurements can be replicated across multiple repeated measures. Test-retest reliability is the closeness of the agreement between the results of successive measurements of the same measure and carried out under the same conditions of measurement.

#### Linear mixed models

As a group-level statistic, reliability refers to the inter-individual or between-subject variability *V_b_* relative to the intra-individual or within-subject variability *V_w_*. Both the intra- and inter-individual variances can be estimated using linear mixed model (LMM). In this study, given a functional graph metric *ϕ*, we considered a random sample of *P* subjects with N repeated measurements of a continuous variable in *M* visits. *ϕ_ijk_* (for *i* = 1,…, *N* and *j* = 1,…, *M*, and *k* =1,…, *P*) denotes the metric from the *k*^th^ subject’s *j*^th^ visit and *i*^th^ measurement occasions. The three-level LMM models *ϕ_ijk_* as the following equations:

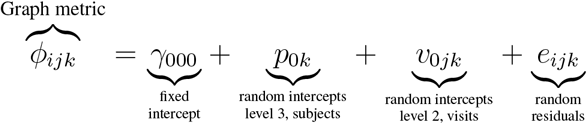

where *γ*_000_ is a fixed parameter (the group mean) and *p*_0*k*_, *v*_0*jk*_ and *e_ijk_* are independent random effects normally distributed with a mean of 0 and variances 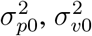, and 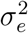. The term *p_0k_* is the subject effect, *v*_0*jk*_ is the visit effect and *e_ijk_* is the measurement residual. Age, gender and interval (Δ*t*) between two visits are covariants.

#### ICC Estimation

These variances are used to calculate the test-retest reliability, which is measured by the dependability coefficient and reflects the absolute agreement of measurements. The dependability coefficient is a form of ICC commonly, which is the ratio of the variances due to the object of measurement versus sources of error. To avoid negative ICC values and obtain more accurate estimation of the sample ICC, the variance components in model are usually estimated with the restricted maximum likelihood (ReML) approach with the covariance structure of an unrestricted symmetrical matrix (Zuo et al., 2013).

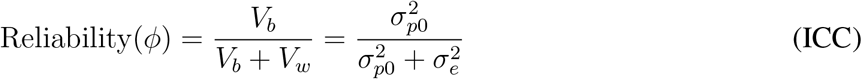

The ICC statistics on the measurement reliability are categorized into five common levels: 0 < ICC ≤ 0.2 (**slight**); 0.2 < ICC ≤ 0.4 (**fair**); 0.4 < ICC ≤ 0.6 (**moderate**); 0.6 < ICC ≤ 0.8 (**substantial**); and 0.8 < ICC < 1.0 (**almost perfect**). A metric with moderate to almost perfect test-retest reliability (ICC ≥ 0.4) is commonly expected in practice. The ICC level should not be judged only based upon the point statistical estimation of ICC but its confidence intervals (CI) (Koo & Li, 2016). We employed the nonparametric conditional bootstrap method for 1000 times to estimate their 95% CIs.

#### Statistics Evaluation

Our analyses can produce big data of 524,160 ICCs (419,328 for the global network metrics). These ICCs are grouped into four categories (parcellation, frequency band, connectivity transformation and edge filtering scheme), each of which has different choices. Given each choice of a category, we estimated its density distributions of ICCs and calculated two descriptive statistics: 1) mean ICC values, which measures the *general reliability* under the given choice; 2) number of almost perfect (noap) ICC values, which measures the *potential reliability* under the given choice.

We further perform Friedman rank sum test to evaluate whether the location parameters of the distribution of ICCs are the same in each choice. Once the Friedman test is significant, we employ the pairwise Wilcoxon signed rank test for post-hoc evaluations to compare ICCs between each pair of the distributions under different choices. The statistical significance levels are corrected with Bonferroni method for controlling the family wise error rate at a level of 0.05. We develop a method to visualize and evaluate the change of ICCs (i.e., reliability gradient) between different choices (Fig. 1c). Specifically, the reliability can be plotted as a function of *V_b_* and *V_w_* in its anatomy plane (Xing & Zuo, 2018; Zuo, Xu, & Milham, 2019). The gradient of reliability between two choices is modeled by the vector (i.e., the black arrow), and decomposed into changes of individual variability. The systematic evaluation on the reliability of the global network metrics determines the optimal network neuroscience by combining the most reliable pipeline choices, which further generated the nodal metrics’ reliability.

## RESULTS

### Whole brain networks are more reliable than cortical networks

We evaluated reliability based on 24 different parcellation choices (Fig. 2a). In the following parts of the paper, we name a parcellation as *‘ParcAbbr-NumberOfParcels’* (e.g., LGP-100 or its whole-brain version wbLGP-458). We found significant differences in ICC distributions across the 24 parcellation choices (Fig. 2b, Friedman rank sum test: *χ*^2^ = 20379.07, *df* = 23, *p* < 2.2 × 10^-16^, effect size *W*_Kendall_ = 0.377). The mean ICCs range from slight (LGP-1000) to substantial (wbLGP-458). Given a particular parcellation and definition of nodes, we illustrate the density distribution of its ICCs under all other strategies (edge definition and metric derivation). Notably, whole-brain parcellations yield higher measurement reliability than parcellations of cerebral cortex on their own (the effect sizes > 0.65). This improvement in reliability seems not simply a bi-product of having more parcels. We chose the parcellations in which the number of parcels (400 ≤ *n* ≤ 1000) almost overlapped between the cortex and the whole brain, and found no correlation between the number of parcels and the median ICCs (*r* = −0.11, *p* = 0.7). We report the mean ICC and the number of almost perfect (noap) ICCs (≥ 0.8) as the descriptive statistics for the density distributions. The wbLGP-458 (mean ICC: 0.671; noap ICC: 519), wbLGP-558 (mean ICC: 0.671; noap ICC: 540) and The wbBNP-568 (mean ICC: 0.664; noap ICC: 511) are the three most reliable choices (see more details of the post-hoc Wilcoxon signed rank test in Table S7). Among the cortical parcellations, the LGP-500 (mean ICC: 0.362; noap ICC: 0), LGP-400 (mean ICC: 0.342; noap ICC: 0) and LGP-600 (mean ICC: 0.340; noap ICC: 0) are the three most reliable choices (Table S3).

To better understand the effect of introducing 358 subcortical parcels into the cortical parcellations, we decomposed the reliability changes into a two-dimensional representation of changes of individual variability (Fig. 2c,d). This idea was motivated by the analysis of reliability derived with individual variability (Xing & Zuo, 2018; Zuo, Xu, & Milham, 2019) as in Fig. 1c. For each ICC under a given parcellation choice, we calculated the related between-subject variability *V_b_* and within-subject variability *V_w_*. Changes in the individual variability associated with the reliability improvements from cortical to whole-brain pipelines were plotted along with Δ*V_b_* and Δ*V_w_* as arrows. These arrows are distributed across the three quadrants (quadI: 0.94%; quadII: 59.99%; quadIII: 39.07%). We noticed that most of these arrows were distributed into the optimal quadrant where the improvements of test-retest reliability by the whole-brain parcellation choices largely attributing to the increases of between-subject variability and decreases of within-subject variability. The decreases of both between-subject and within-subject variability may also strengthen the measurement reliability (the suboptimal quadIII in Fig. 2).

### Spontaneous brain activity portrays more reliable networks in higher slow bands

Brain oscillations are hierarchically organized, and their frequency bands were theoretically driven by the natural logarithm linear law (Buzsaki & Draguhn, 2004). By analogy, rfMRI oscillations can, similarly, be partitioned into distinct frequency bands. Advanced by the fast imaging protocols (TR = 720ms), HCP test-retest data allows to obtain more oscillation classes than traditional rfMRI acquisitions (typical TR = 2s). We incorporate the Buzsaki’s framework with the HCP dataset using the DREAM toolbox (Gong et al., 2021) in the Connectome Computation System to decompose the time series into the six slow bands (Fig. 3a): **slow-6** (0.0069-0.0116 Hz),**slow-5** (0.0116-0.0301 Hz), **slow-4** (0.0301-0.0822 Hz), **slow-3** (0.0822-0.2234 Hz), **slow-2** (0.2234-0.6065 Hz), **slow-1**^-^ (0.6065-0.6944 Hz).

We noticed that, due to the limited sampling rate (TR), this **slow-1**^-^ only covers a small part of the full **slow-1** band (0.6065-1.6487 Hz) – we indicate this above. We also included the frequency band, **slow-emp** (0.01-0.08 Hz) for the sake of comparison, as it is covers a range commonly used in rfMRI studies. A significant effect on order (*χ*^2^ = 9283.536, *df* = 6, *p* < 2.2 × 10^-16^, *W*_Kendall_ = 0.192) across the frequency bands was revealed based on the density distributions of ICC (Fig. 3b): slow-2, slow-1^-^, slow-3, slow-emp, slow-4, slow-5, slow-6. Post-hoc paired tests indicated that any pairs of neighbouring bands are significantly different from one another, with measurement reliability increasing with faster frequency bands. Note, however, that slow-1^-^ (mean ICC: 0.564) did not fit into this trend, possibly due to its limited coverage of the full band. But remarkably, slow-1^-^ exhibited the largest number of almost prefect ICCs for potential reliability (noap ICC: 1746). Slow-emp (mean ICC: 0.519; noap ICC: 434) contains overlapping frequencies with both slow-4 (mean ICC: 0.560; noap ICC: 441) and slow-5 (mean ICC: 0.494; noap ICC: 285), and higher ICCs than the two bands but the effect sizes are small to moderate (slow-emp vs. slow-4: 0.193; slow-emp vs. slow-5: 0.485). Slow-6 is the choice with the lowest ICCs (mean ICC: 0.331; noap ICC: 154) compared to other bands (large effect sizes: *r* > 0.57).

To visualize reliability variation across frequency bands, we plotted a trajectory tracing reliability flow along the five full (slow-6 to slow-2) bands in the reliability plane, whose axes correspond to between-*versus* within-subject variability (Fig. 3c). As expected, this nonlinear trajectory contains two stages of almost linear changes of the network measurement reliability from slow to fast oscillations: whole brain versus cortex. In each case, the reliability improvements attribute to both increases of between-subject variability and decreases of within-subject variability while the improvements of whole-brain network measurement reliability were largely driven by the increased variability between subjects.

### Topological economics individualize highly reliable functional brain networks

Estimating functional connections can be highly challenging due to the absence of a ‘ground truth’ human functional connectome. To provide a reliable way of building candidate edges of the connections, we sampled the 12 schemes on graph edge filtering (Fig. 4a), which turn a fully connected matrix into a sparse graphical representation of the corresponding brain network. These schemes can be categorized into two classes: threshold-based *versus* topology-based schemes. Threshold-based schemes usually use a threshold to preserve those edges whose strengths are above a cutoff value, such as ABS_05_, PROP_10_, PROP_20_, DEG_5_, DEG_15_. Threshold-based schemes are widely used in network neuroscience and ignore the intrinsic topological structure of the entire brain network (e.g, leading to multiple connected components or isolated nodes). In contrast, topology-based schemes such as MST, OMST, PMFG and TMFG come from other scientific disciplines and are optimized based on the entire network topology (see **Materials and Methods**). To combine both the TMFG’s efficiency and OMST’s accuracy, we proposed the OTMFG. All the schemes are plotted in the plane of cost *versus* global-cost efficiency to better visualize the economical properties of the derived networks (Fig. 4b). These plots are fitted into the topographic (contour) maps where the local maxima for each filtering choice is labeled as a circle. The human brain networks achieve higher global efficiency with lower cost using topology-based schemes compared to threshold-based schemes, suggesting increasingly optimal economics.

Significant differences in test-retest reliability were detectable across these 12 edge-filtering schemes (*χ*^2^ = 9784.317, *df* = 11, *p* < 2.2 × 10^-16^, *W*_Kendall_ = 0.189, see Fig. 4c). Among the topology-based schemes, OMST (mean ICC: 0.608; noap ICC: 765), OTMFG (mean ICC: 0.602; noap ICC: 781) and TMFG (mean ICC: 0.570; noap ICC: 767) were the three most reliable choices. They showed significantly greater reliability than the three most reliable threshold-based, respectively: PROP20 (mean ICC: 0.593; noap ICC: 632), PROP_10_ (mean ICC: 549; noap ICC: 445) and GCE (mean ICC: 0.533; noap ICC: 352). Mean reliability of MST are slight to fair (mean ICC: 0.309) but its number of almost perfect reliability (noap ICC:362) is still higher than all threshold-based schemes except PROP_10_ and PROP_20_.

Network measurements are labeled based on topology and threshold groups and projected onto the reliability anatomy plane, whose axes represent between- and within-subject variability (Fig. 4d). The contour maps are reconstructed for each scheme based upon the individual variability of all the related network measurements. The topology-based methods (red) showed overall higher ICCs than the threshold-based methods (blue), improvements that could be attributed to increases in between-subject variability and decreases of within-subject variability. These observations are consistent between cortex and whole brain networks while topology-based whole brain network are almost perfectly reliable (meaning almost perfect reliability, i.e., ICC ≥ 0.8).

We also explored connection transformation and edge weights, two factors included in edge filtering, the choices of connectivity transformation and weighing edges, regarding their measurement reliability. Positive (Eq.pos) (mean ICC: 0.512; noap ICC: 1,031) and exponential (Eq.exp) transformation (mean ICC: 0.509; noap ICC: 1,855) were the two most reliable choices. Comparing to the positive and absolute (Eq.abs) (mean ICC: 0.508; noap ICC: 1,050) transformation, the exponential and distance-inverse (Eq.div) (mean ICC: 0.500; noap ICC: 1,031) transformation show larger number of almost perfect ICCs. Weighted graphs are also more reliable than the binary graphs while the normalized weighted graphs demonstrated the highest ICCs, reflecting both the increased between-subject variability and decreased within-subject variability.

### Network integration and segregation can serve reliable metrics of information flow

The previous extensive data analysis suggests that the optimally reliable pipeline should: 1) define network nodes using a whole-brain parcellation, 2) filter the time series with higher frequency bands, 3) transform the connectivity using positive transformation, 4) construct network edges using topology-based methods and normalized weights. Using the optimal pipelines, we evaluated the reliability levels of various metrics from network neuroscience and their differences across individuals. Focusing on the optimized pipeline with the highest ICCs of the various choices (wbLGP-458, slow-2, pos, normalized weights, OMST), we reported test-retest reliability of the measurements as well as their corresponding individual variability. In Fig. 5a, we found that the global network measurements of information segregation and integration are at the level of almost perfect reliability except for the modularity *Q* (ICC=0.46, 95% CI = [0.252,0.625]). These high-level ICCs are derived with large between-subject variability and small within-subject variability (Fig. 5b). These findings are reproducible across the other two parcellation choices (wbCABP-718, wbBNP-458). In consideration of “Ease-of-Use” for researchers and higher cortical resolution, we mapped the “Out-of-the-Box” Cole-Anticevic Brain-wide Network Partition (wbCABP-718) for nodal metrics visualization.

**Figure 5.**
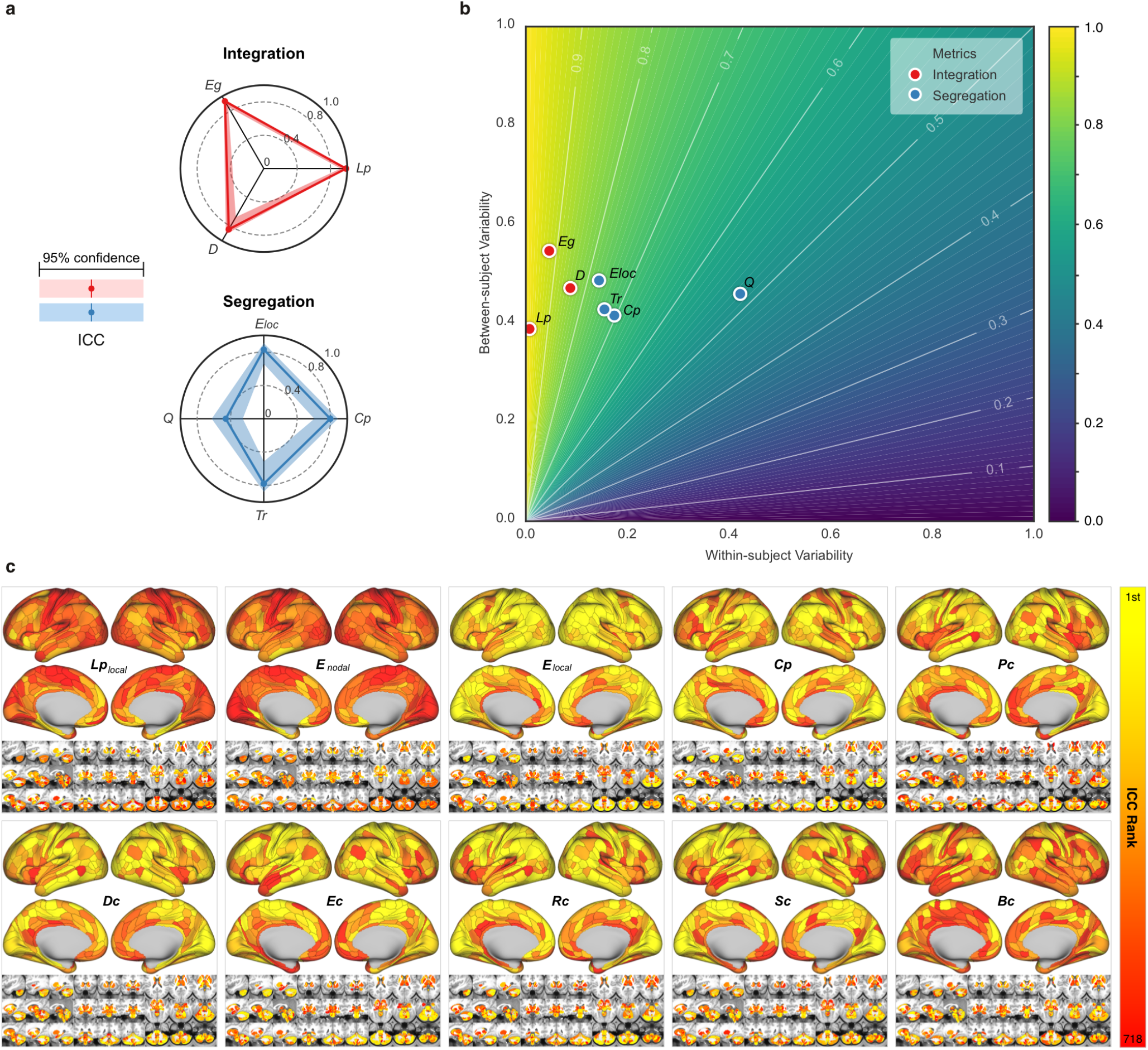
Measurement reliability and variability of global/nodal network metrics under the optimized pipeline. (a) Spider plots are visualized for ICCs (test-retest) with the 95% confidence intervals (CIs, shadow bands) for the global metrics of network integration, segregation. Integrative measures include average shortest path length (*L_p_*), global efficiency (*E_g_*), and pseudo diameter (*D*). Segregation measures include clustering coefficient (*C_p_*), local efficiency (*E*_loc_), transitivity (*T_r_*), modularity (*Q*). (b) The reliability anatomy was plotted as a function of between-subject variability (*V_b_*) and within-subject variability (*V_w_*). (c) Ranks of ICCs across the 360 cortical parcels and the 358 subcortical parcels in the optimal pipeline (wbCABP-718, slow-2, pos, OMST) are depicted. Ten nodal metrics are assessed including local characteristic path length of node (*L*_plcoal_), nodal efficiency (*E*_Eodal_), local efficiency of nodes (*E*_local_), nodal clustering coefficient (*C_p_*), pagerank centrality (*P_c_*), degree centrality (*D_c_*), eigenvector centrality (*E_c_*), resolvent centrality (*R_c_*), subgraph centrality (*S_c_*) and betweeness centrality (*B_c_*).

Similar to the global metrics, shortest path length *L_p_* and nodal efficiency *E_nodal_* exhibited the highest ICCs (almost perfect test-retest reliability) while ICCs of other nodal metrics remained less than 0.6. To visualize node-level network metrics, we reported results derived from the wbCABP-718 choice. To improve spatial contrasts of reliability, we ranked the parcels according to their ICCs and visualized the ranks in Fig. 5c. Most nodal metrics are more reliable across the 360 cortical areas than the 358 subcortical areas (Wilcoxon tests: all p-values less than 0.001, corrected for multiple comparisons). However, *L_p_*, *E*_nodal_ and *B_c_* exhibited higher across subcortical areas than cortical areas (corrected *p* < 0.001). Across the human cerebral cortex, the right hemispheric areas demonstrated more reliable *C_p_* (corrected *p* < 0.0036) than the left hemispheric areas. Interesting patterns of the reliability gradient are also observable along large-scale anatomical directions (dorsal>ventral, posterior>anterior) across the nodal metrics of information segregation and centrality. These spatial configuration profiles on the reliability reflected their correspondence on inter-individual variability of these metrics, characterising the network information flow through the slow-2 band.

### Building an open resource for reliable network neuroscience

The results presented here represent huge costs in terms of computational resources (more than 1,728,000 core-hours on **CNGrid**, supported by Chinese Academy of Sciences (http://cscgrid.cas.cn). Derivations of the ICCs and their linear mixed models were implemented in **R** and **Python**. As our practice in open science, we have started to provide an online platform on the reliability assessments (http://ibraindata.com/research/ifNN/reliabilityassessment). The big reliability data were designed into an online database for providing the community a resource to search reliable choices and help the final decision-making. The website for this online database provided more details of the reliability data use. We shared all the codes, figures and other reliability resources via the website (http://ibraindata.com/research/ifNN/database).

## DISCUSSION

This study examined the series of processing and analysis decisions in constructing graphical representations of brains’ intrinsic spontaneous activity. The focus, here, was on identifying the pipeline that generated reliable, individualized networks and network metrics. The results of our study suggest that to derive reliable global network metrics showing higher inter-individual variances and lower intra-individual variances, one should use whole-brain parcellations to define network nodes, focus on higher frequencies in the slow band for time-series filtering to derive the connectivity, and use the topology-based methods for edge filtering to construct sparse brain graphs. Regarding network metrics, multi-level or multi-modal metrics appear more reliable than single-level or single-model metrics. Derive reliable measurements is critical in network neuroscience, especially for translating network neuroscience into personalized practice. Based on these results, we provide four principles towards optimal functional network neuroscience for reliability of measuring individual differences.

### Principle I: Use a whole brain parcellation to define network nodes

The basic unit of a graph is the node. However, variability across brain parcellations can yield different graphs, distorting network metrics and making it difficult to compare findings across studies (Wang et al., 2009; Zalesky et al., 2010). In many clinical applications (Fornito et al., 2015; Matthews & Hampshire, 2016), researchers aim to identify disease-specific connectivity profiles of the whole brain, including cortical and subcortical structures, as well as cerebellum. A recent review has raised the concern that many studies have focused on restricted sets of nodes, e.g., cortex only, and called a field standard for the best practices in clinical network neuroscience (Hallquist & Hillary, 2019), which requires almost perfectly reliable measurements (Xing & Zuo, 2018). Our meta-reliability assessments revealed such high reliability of measurements made involving functional brain networks can be achieved, through the inclusion of high-resolution subcortical nodes. This provides strong evidences that the whole-brain node use should be part of the standard analysis pipeline for network neuroscience applications. These improvements of reliability can be attributed to increases in between-subject variability coupled with reductions in within-subject variability relative to networks of cortical regions alone. One possible neuroanatomical explanation is that distant areas of cerebral cortex are interconnected by the basal ganglia and thalamus while also communicating with different regions of the cerebellum *via* polysynaptic circuits, forming an integrated connectome (Bostan & Strick, 2018). These subcortical structures have been suggested to play a role in both primary (e.g., motor) and higher-order function (e.g., learning and memory) while studies using rfMRI have delineated the resting-state functional connectivity (RSFC) maps between these subcortical structures and cortical networks of both primary and high-order functions. Interestingly, a recent work revealed that inter-individual variance in cerebellar RSFC networks exceeds that of cortex (Marek et al., 2018). Meanwhile, these RSFC maps are highly individualized and stable within individuals (D. Greene et al., 2020), indicating that they possess reliable characteristics. In line with our observations, we argue that inclusion of the subcortical structures as network nodes can enhance the between-subject variability and stabilize the within-subject variability by providing a more comprehensive measurements on the entirety of the brain connectivity.

### Principle II: Generate functional networks using spontaneous brain activity in multiple slow bands

It has been a common practice in rfMRI research to estimate the RSFC profile based on BOLD time series of the intrinsic spontaneous brain activity from the low-frequency (0.01 - 0.1 Hz or 0.01 - 0.08 Hz). However, the test-retest reliability of RSFC measurements derived from this frequency band has been limited, with ICCs less than 0.4 (Noble et al., 2019; Zuo & Xing, 2014). Still existing studies, however, have advocated adopting a multi-frequency perspective to examine the amplitude of brain activity at rest (Zuo et al., 2010) and its network properties (Achard et al., 2006). This approach has been spurred along by recent advances in multi-banded acquisitions and fast imaging protocols, offering rfMRI studies a way to examine spontaneous brain activity at much higher frequencies that may contain neurobiologically meaningful signals (Gong et al., 2021). Our study provides strong evidence of highly reliable signals across higher slow-frequency bands, which are derived with the hierarchical frequency band theory of neuronal oscillation system (Buzsaki & Draguhn, 2004). Specifically, a spectrum of reliability increases was evident from slow bands to fast bands. This reflects greater variability of the network measurements between subjects and less measurement variability within subject between the higher and lower bands of the slow frequencies. In theory, each frequency band has an independent role in supporting brain function. Lower frequency bands are thought to support more general or global computation with long-distance connections to integrate specific or local computation, which are driven by higher slow bands based on short-distance connections (Buzsáki, 2009). Our findings of high reliability (inter-individual differences) are perfectly consistent with this theory from a perspective of individual variability. Previous findings have found that high-order associative (e.g., default mode and cognitive control) networks are more reliable than the primary (e.g., somatomotor and visual) networks (Noble et al., 2019; Zuo & Xing, 2014; Zuo, Xu, & Milham, 2019). A novel frequency-based perspective on these network-level individual differences can be inspired directly by our observations on the multiple bands.

### Principle III: Optimize topological economy to construct network connections at individual level

There is no gold standard on for human functional connectomes, leading to plurality of approaches for inferring and constructing brain network connections. Threshold-based methods focus on the absolute strength of connectivity, retaining connections that are above some user-defined threshold and oftentimes involve applying the same threshold to all subjects. Although this approach mitigate potential biases in network metrics associated with differences in network density, it may inadvertently also lead to decreased variability between subjects. This is supported by our results showing that threshold-based method yield low reliability of network measurements. On the other hand, the human brain is a complex network that is also near-optimal in terms of connectional economy, balancing tradeoffs of cost with functionality (Bullmore & Sporns, 2012). In line with this view, certain classes of topology-based methods for connection definition may hold promise for individualized network construction.

Specifically, each individual brain optimizes its economic wiring in terms of cost and efficiency, reaching a trade-off between minimizing costs and allowing the emergence of adaptive topology. Our results demonstrate that such highly individualized functional connectomes generated by the topology-based methods are more reliable than those by the threshold-methods. This reflects the increases of individual differences in functional connectomes attributing to the optimal wiring economics at individual level. The topological optimization also brings other benefits such as ensuring that a graph forms a single connected component and preserving weak connections. Indeed, there is increasing evidence supporting the hypothesis that weak connections are neurobiologically meaningful and explain individual differences in mind, behavior and demographics as well as disorders (Santarnecchi, Galli, Polizzotto, Rossi, & Rossi, 2014). Weak connections in a graph may be consistent across datasets and reproducible within the same individual over multiple scanning sessions and therefore be reliable. Weak connections might also play non-trivial roles in transformed versions of the original brain network, e.g. so-called “edge-based functional connectivity” (Faskowitz, Esfahlani, Jo, Sporns, & Betzel, 2020). Among these topology-based methods, MST is the simplest and promising filtering method if computational efficiency is the priority. MST can obtain a graph with the same number of nodes and edges, and it is not sensitive to scaling effects, because its structure only depends on the order rather than the absolute values of the edges. Although MST loses some local network measurements due to the limited number of edges, it has some other unique metrics that can be calculated (e.g., leaf fraction, tree hierarchy). A better alternative might be TMFG which is computationally very efficient and statistically robust, while the OMST and OTMFG are the most reliable choices by prioritizing significant individual differences.

### Principle IV: Characterise information flow with both network integration and segregation metrics

Intrinsic functional networks reflect the outcomes of communication processes and information flows between pairs of brain regions. How the information and other signals propagate between pairs of brain regions can be assayed using network neuroscientific metrics and is essential to understanding normative connectome function and its variation in clinical settings. While the ground truth of functional connectome remains unknown (and may not exist), network models can help validate the imaging-based reconstructions of human functional networks (D. Bassett et al., 2020). From a perspective of individual differences, reliable network measures are the basis of achieving valid ifNN measurements (Zuo, Xu, & Milham, 2019). Our findings indicated that both the brain network segregation and integration could be reliably measured with functional connectomics using rfMRI by the optimized pipelines. At the global level, measures of information integration, e.g. characteristic path length and efficiency, were more reliable than those of information segregation, e.g. modularity and clustering coefficient. Our results also revealed that measures of integration were more stable across different scan sessions (i.e., the test-retest) for an individual subject than the segregation measurements while the inter-individual variability are measured at the similar level for both integration and segregation metrics. At nodal level, mapping reliability of the network measurements revealed interesting spatial patterns. Specifically, we found that cortical areas were generally associated with more reliable local measurements compared to subcortical areas. This may reflect different functional roles for human cortex and subcortex. For example, the differences in reliability of path-based metrics might reflect the fact that there are more cortical within-community paths while between-community paths are more common in subcortex. Beyond this cortical-subcortical gradient, reliability of the nodal information flow also fit the left-right asymmetry and dorsal-ventral as well as posterior-anterior gradient, implying the potential validity of individual differences in information flow attributing to evolutionary, genetic and anatomical factors (Chen et al., 2013; Rakic, 2009). To facilitate the utility of reliable network integration and segregation metrics in ifNN, we integrated all the reliability resources into an online platform for reliability queries on specific metrics of information flow (http://ibraindata.com/research/ifNN).

## REPRODUCIBILITY, GENERALIZABILITY AND CONCLUSION

Both reproducibility and generalizability are cornerstones of modern sciences, and remain challenging as a scientific research frontier (Munafo et al., 2017; Yarkoni, 2022). In this research, we adopt a big data approach by deeply sampling the parameters (more than 524k parametric settings) of various steps in the network construction and analysis pipeline to systematically explore the reliability of functional brain network measurements. This provided robust experimental evidence supporting four key principles that will foster optimal ifNN research and application. These principles can serve as the base for building guidelines on the use of ifNN to map individual differences. Standard guidelines are essential for improvements of reproducibility and generalizability in the research practice, and our work provide basic resources initiating such standardization in future network neuroscience. We note, however, that while our approach was extensive, it was not exhaustive (likely impossible) – the analytical sampling procedure could miss many other existing choices. The processing decisions that yield reliable connectomic measurements may yield the most reliable network statistics, but there may be another way to process data that yields overall a higher level of reliability in network measures. Regarding the statistical benefits of our sampling analytics in the parametric space of the ifNN pipelines, we discuss about the implications of the present research for reproducible and generalizable network neuroscience as following.

The rfMRI datasets minimally preprocessed by the HCP pipeline are employed for our study while many different pipelines are available for rfMRI data preprocessing (see a list of pipelines in (Xu et al., 2015)). These different pipelines vary across parametric settings and orders of various steps of preprocessing, and thus can have different impacts on the reliability of measuring spontaneous brain activity (Li et al., 2022). Therefore, it is very important to validate whether the present findings are reproducible under another preprocessing pipeline. Accordingly, we repeated our analyses by leveraging another widely-accepted preprocessing pipeline, fMRIPrep (Esteban et al., 2019). As documented in the supplementary materials, the major findings supporting the principal guidelines are reproducible while the measurement reliability derived with fMRIPrep are generally lower than those with the HCP pipeline. Various within-pipeline parametric settings also exist other choices not sampled by our experimental design but remain potentials for further investigation. For example, edge filtering methods are commonly used to identify and retain only the most important edges in a graph, based on criteria such as statistical significance or functional relevance. However, this approach has the potential to introduce bias and subjectivity in the selection process, and may not fully capture the higher-order structure of a network system. Algebraic topology, as demonstrated by Giusti, Ghrist, and Bassett (2016) and other recent studies, offers a promising alternative for high-order edge filtering. By representing relationships between objects as higher-dimensional simplices instead of edges, simplicial complexes can characterize polyadic interactions and capture more nuanced aspects of the complex network organization. With the increasing availability of computational tools for the application of algebraic topology to real data, this framework has the potential to surpass graph theory in understanding the complexities of neural systems.

Pipelines of generating highly reliable measurements are central to experimental design of studying individual differences (Matheson, 2019; Zuo, Xu, & Milham, 2019). Given a statistical power, for a fixed sample size, experiments designed with more reliable pipelines can detect bigger effects of interests. On the other side, to detect a fixed effect size, experiments designed with more reliable pipelines would be more powerful or logistically economical (e.g., need less samples). This has very important implications on the recent arguments about ‘big data versus small data’ (Marek et al., 2022; Rosenberg & Finn, 2022; Tibon, Geerligs, & Campbell, 2022), which must take the reliability into account at first place of designing an experiment (Gratton, Nelson, & Gordon, 2022), and has been increasingly appreciated by the field of network neuroscience (Helwegen, Libedinsky, & van den Heuvel, 2023). From a perspective of experimental design, reliability is more straightforward to reproducibility but validity is related to generalizability. Therefore, we clarify that the measurement reliability is not the final goal but the validity (Finn & Rosenberg, 2021; Noble et al., 2021), which is not easily ready for a direct examination as reliability assessment (Zuo, Xu, & Milham, 2019). The reliable pipeline we proposed produced biologically plausible findings according to the four principles as we discussed, likely reflecting its potential validity of measuring individual differences in intrinsic brain functional organization. Validation on the use of our proposed principles represents a promising arena for fostering future network neuroscience studies such as personality (Hilger & Markett, 2021) or brain developmental charts (Bethlehem et al., 2022), with potential novel fMRI paradigms (Elliott, Knodt, & Hariri, 2021; Finn, Glerean, Hasson, & Vanderwal, 2022) or more precise neuroimaging technology (Toi et al., 2022).

Reliability does not necessarily equate to but indeed provides an up bound of validity. In some cases, increasing reliability may cause a decrease in validity, particularly if the sources of reliability are not related to the underlying construct of interests. For example, physiological noise and head motion can be highly reliable as biological traits, but may not be involved in the investigated cognitive processes. Previous studies have shown that head motion can introduce artifacts into the fMRI data, which can affect the reliability of functional connectivity measures (Power et al., 2014). In particular, head motion may have a non-uniform impact on different edge filtering methods and network metrics. Certain methods that rely on the strength of functional connections, such as threshold-based approaches, may be more sensitive to head motion artifacts than topology-based methods that focus on the overall structure of the network. Measures that are highly reliable due to the inclusion of these contaminants may not be valid indicators of the underlying construct. However, head motion may not always be purely noise and may contain some neurobiologically meaningful signals (Zeng et al., 2014; Zhou et al., 2016). Therefore, it is important to carefully consider the potential impact of head motion when choosing an edge filtering method and interpreting the resulting functional connectivity measures, as well as the trade-off between controlling for motion artifacts and preserving potentially meaningful signal in the data. In the context of graph theory, noise can affect both the reliability and validity of graph metrics. For example, noise in the data can result in higher reliability of certain graph metrics, but this may not necessarily reflect the true underlying network structure. This is because noise may lead to inflated correlations between certain regions, resulting in over-estimations of network connectivity and thus higher reliability. However, these measurements may not be valid indicators of the true network structure and may not accurately reflect the underlying cognitive processes being studied. On the other hand, the removal of noise may lead to decreased reliability, but may improve the validity of the measurement by reducing the influence of unrelated sources of variance. The optimal choices for maximizing reliability in our study may also have implications for interpretability and generalizability. For example, the inclusion of subcortical structures in the parcellation scheme may increase the interpretability of the results, as these structures play a key role in the functional organization of the brain. On the other hand, the choice of connectivity transformation and edge weighting may have implications for the generalizability of the results, as different methods may produce different results depending on the specific characteristics of the data. Further research is warranted to fully understand the consequences of these choices on interpretability, generalizability, and other aspects of the measurement process.

The guidelines we proposed for rfMRI-based network neuroscience may also provide insights for network neuroscience computation by leveraging task-fMRI or movie-fMRI. These two paradigms have gained increasing attention in recent years as a means of measuring functional connectomes (Cole, Bassett, Power, Braver, & Petersen, 2014; Cole et al., 2013; Finn et al., 2022). The reliability and predictive power of these measures have been the subject of a number of studies. For instance, a study by Gao et al. (2020) found that the reliability of movie-fMRI connectivity was influenced by the complexity and duration of the movie stimulus, with more complex and longer stimuli resulting in higher test-retest reliability. The results support the notion that task-fMRI and movie-fMRI can produce more reliable connectivity measures with greater predictive power for individual differences in cognitive and mental health measures compared to rfMRI, particularly for tasks and stimuli that elicit strong and sustained activation. According to the relationships among rest, task and movie as well as other naturalistic states of the human brain as a systems entity (Cole, Ito, Bassett, & Schultz, 2016; Finn, 2021; McCormick, Arnemann, Ito, Hanson, & Cole, 2022), we speculate that the four principles are generalizable to functional network neuroscience based on these non-rest brain states. However, we note that more research is warranted to fully understand the underlying mechanisms and generalizability of these findings to different task and movie paradigms as well as their translational applications (Eickhoff, Milham, & Vanderwal, 2020; Finn & Rosenberg, 2021).

Population diversity plays a critical factor of assuring the generalizability in studying individual differences (A. S. Greene et al., 2022; Ricard et al., 2023). This has been responded by the emerging new stage of cognitive neuroscience, namely population neuroscience (Falk et al., 2013; Paus, 2010). Psychometric studies are particularly required for population neuroscience due to the core aim of measuring individual differences in brain and mind developmental during the life span (Zuo et al., 2017, 2018). The design of a psychometric study is normally recommended to recruit a group of participants who are stable across the duration of investigation. This makes the interpretation of within-subject variability straightforward as the subject-independent random noise, and the reliability assessment more precisely; and also is why most psychometric studies were done in adults although some in children (but with very short duration). When studying the lifespan development, one must consider the specific research aims and the underlying assumptions of the study for addressing the reliability trade-off between maximizing between-individual variability and minimizing within-individual variability. For example, if the goal of the study is to identify developmental or lifespan-related trajectories, it may be more important to prioritize maximizing between-individual variability in order to capture the full range of individual differences. In this case, techniques such as motion scrubbing or outlier detection may be employed to minimize within-individual variance, even if this leads to a decrease in overall reliability. On the other hand, if the focus of the study is on assessing within-subject changes over time, it may be more important to minimize within-individual variance in order to accurately capture changes in brain function. In this case, techniques such as temporal smoothing or denoising may be employed to increase reliability, even if this leads to a decrease in between-individual variability. It is also important to consider the potential impacts of these choices on the validity of the measurement. For example, if motion scrubbing or outlier detection leads to the exclusion of a large number of subjects or time points, this may introduce bias and reduce the generalizability of the results regarding the limited diversity. Careful consideration of these trade-offs is therefore essential in order to ensure that the chosen approach is appropriate for the specific research aims and assumptions of the study, especially the population neuroscience research.

## ACKNOWLEDGMENTS

The neuroimaging data were provided by the HCP WU-Minn Consortium, which is funded by the 16 NIH institutes and centers that support the NIH Blueprint for Neuroscience Research 1U54MH091657 (PIs: David Van Essen and Kamil Ugurbil), the McDonnell Center for Systems Neuroscience at Washington University. The institutional Research Ethics Boards approved their HCP WU-Minn Consortium studies. Written informed consent was obtained from all participants.

## COMPETING INTERESTS

The authors declare that they have no conflicts of interest in relation to this manuscript.

## AUTHOR CONTRIBUTION

Chao Jiang: Conceptualization; Data curation; Formal analysis; Methodology; Software; Visualization; Writing - original draft; Writing - review & editing. Richard F. Betzel: Funding acquisition; Writing - review & editing. Ye He: Writing - review & editing; Conceptualization. Yin-Shan Wang: Conceptualization; Writing - review & editing. Xiu-Xia Xing: Conceptualization; Software; Resources; Supervision; Writing - review & editing. Xi-Nian Zuo: Conceptualization; Funding acquisition; Resources; Supervision; Visualization; Writing - original draft; Writing - review & editing.

## FUNDING INFORMATION

This study is supported by the STI 2030 - Major Project 2021ZD0200500. We thank the *Chinese Data-sharing Warehouse for In-vivo Imaging Brain* at National Basic Science Data Center for informatics resources, the *Research Program on Discipline Direction Prediction and Technology Roadmap* of China Association for Science and Technology, and *Learning: Brains, Machines and Children* from the Indiana University Office of the Vice President for Research Emerging Area of Research Initiative.

## Notes

http://ibraindata.com/research/ifNN

### Competing Interest Statement

The authors have declared no competing interest.

### Summary of Updates

Sections on Introduction and Discussions are updated to clarify both reproducibility and generalizability; Figures 1,2 and 5 are revised; author affiliations and order are updated; Supplemental files are updated.

