## Supplementary Materials for "Optimizing network neuroscience computation of individual differences in human spontaneous brain activity for test-retest reliability"

**7 fMRIPrep pipeline** **67**

**References** **71**

### 1 Parcellations - Node Definition

#### 1.1 ICC Density distribution

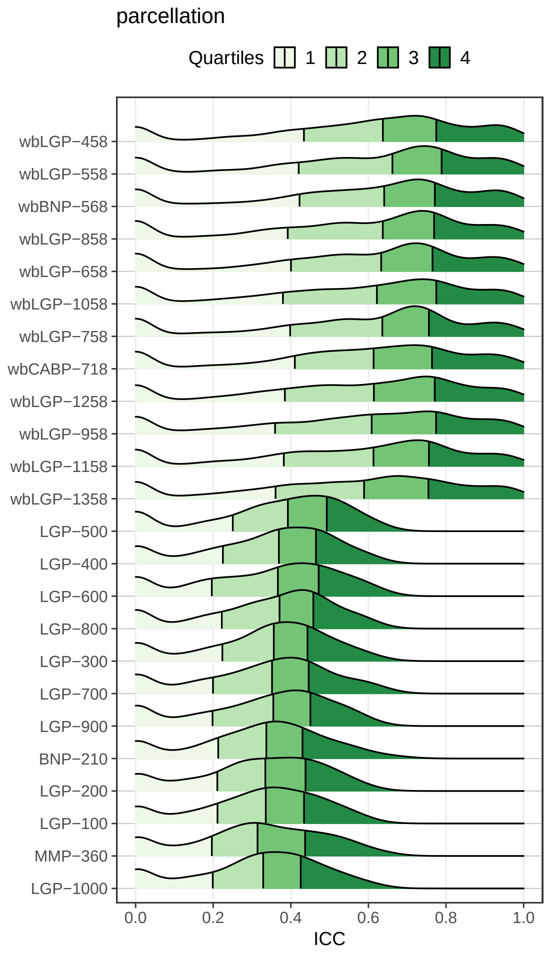

**Fig. S1.** Parcellation - Density distribution

#### 1.2 ICC Almost Perfect ( ICCs > 0.8 )

**Table S1.** Parcellation - Number of ICCs > 0.8

| parcellation | n | ratio |
|---|---|---|
| wbLGP-558 | 540 | 0.2295918 |
| wbLGP-458 | 519 | 0.2206633 |
| wbBNP-568 | 511 | 0.2172619 |
| wbLGP-958 | 486 | 0.2066327 |
| wbLGP-1058 | 484 | 0.2057823 |
| wbLGP-1258 | 482 | 0.2049320 |
| wbLGP-858 | 478 | 0.2032313 |
| wbCABP-718 | 463 | 0.1968537 |
| wbLGP-658 | 444 | 0.1887755 |
| wbLGP-1358 | 440 | 0.1870748 |
| wbLGP-1158 | 416 | 0.1768707 |
| wbLGP-758 | 413 | 0.1755952 |

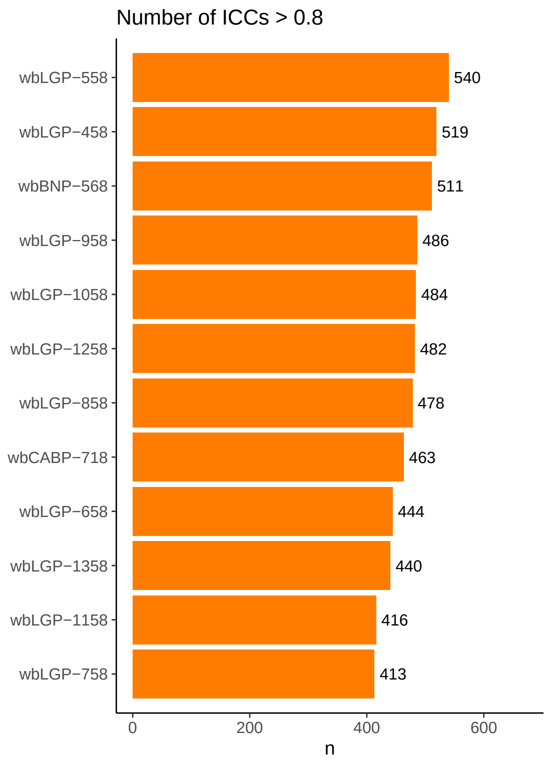

**Fig. S2.** Parcellation - Number of ICCs > 0.8

#### 1.3 Substantial or Above ( ICCs > 0.6 )

**Table S2.** Parcellation - Number of ICCs > 0.6

| parcellation | n | ratio |
|---|---|---|
| wbLGP-458 | 1328 | 0.5646259 |
| wbLGP-558 | 1320 | 0.5612245 |
| wbBNP-568 | 1298 | 0.5518707 |
| wbLGP-858 | 1263 | 0.5369898 |
| wbLGP-758 | 1248 | 0.5306122 |
| wbLGP-658 | 1245 | 0.5293367 |
| wbLGP-1058 | 1239 | 0.5267857 |
| wbCABP-718 | 1224 | 0.5204082 |
| wbLGP-1258 | 1216 | 0.5170068 |
| wbLGP-1158 | 1211 | 0.5148810 |
| wbLGP-958 | 1209 | 0.5140306 |
| wbLGP-1358 | 1141 | 0.4851190 |
| LGP-500 | 97 | 0.0412415 |
| LGP-700 | 92 | 0.0391156 |
| LGP-600 | 85 | 0.0361395 |
| BNP-210 | 71 | 0.0301871 |
| MMP-360 | 66 | 0.0280612 |
| LGP-400 | 51 | 0.0216837 |
| LGP-300 | 47 | 0.0199830 |
| LGP-800 | 46 | 0.0195578 |
| LGP-900 | 38 | 0.0161565 |
| LGP-1000 | 37 | 0.0157313 |
| LGP-200 | 24 | 0.0102041 |
| LGP-100 | 21 | 0.0089286 |

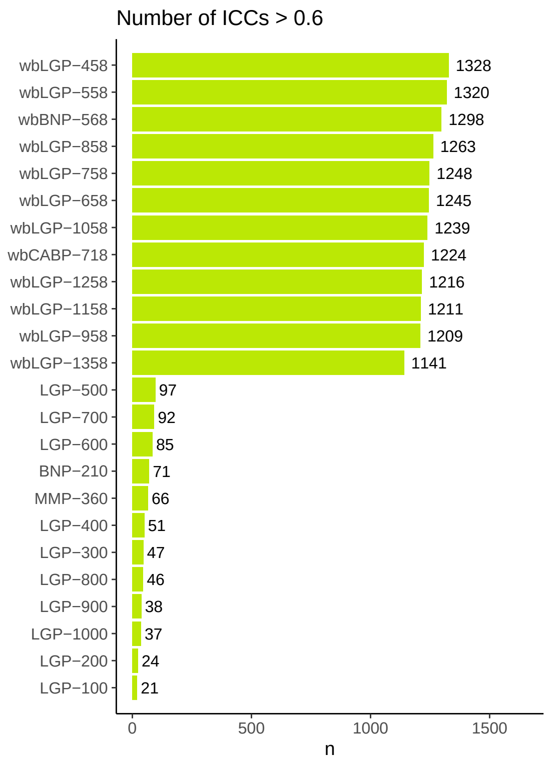

**Fig. S3.** Parcellation - Number of ICCs > 0.6

#### 1.4 Descriptive statistics Mean

**Table S3.** Parcellation - ICC Mean

| parcellation | variable | n | mean | mean_z2r |
|---|---|---|---|---|
| wbLGP-458 | ICC.z | 2352 | 0.813 | 0.6712419 |
| wbLGP-558 | ICC.z | 2352 | 0.812 | 0.6706921 |
| wbBNP-568 | ICC.z | 2352 | 0.800 | 0.6640368 |
| wbLGP-858 | ICC.z | 2352 | 0.777 | 0.6509814 |
| wbLGP-658 | ICC.z | 2352 | 0.775 | 0.6498275 |
| wbLGP-1058 | ICC.z | 2352 | 0.770 | 0.6469295 |
| wbCABP-718 | ICC.z | 2352 | 0.765 | 0.6440126 |
| wbLGP-758 | ICC.z | 2352 | 0.765 | 0.6440126 |
| wbLGP-1258 | ICC.z | 2352 | 0.764 | 0.6434270 |
| wbLGP-958 | ICC.z | 2352 | 0.760 | 0.6410770 |
| wbLGP-1158 | ICC.z | 2352 | 0.746 | 0.6327565 |
| wbLGP-1358 | ICC.z | 2352 | 0.731 | 0.6236768 |
| LGP-500 | ICC.z | 2352 | 0.379 | 0.3618387 |
| LGP-400 | ICC.z | 2352 | 0.356 | 0.3416859 |
| LGP-600 | ICC.z | 2352 | 0.354 | 0.3399182 |
| LGP-800 | ICC.z | 2352 | 0.354 | 0.3399182 |
| LGP-300 | ICC.z | 2352 | 0.347 | 0.3337123 |
| LGP-700 | ICC.z | 2352 | 0.344 | 0.3310437 |
| LGP-900 | ICC.z | 2352 | 0.340 | 0.3274774 |
| BNP-210 | ICC.z | 2352 | 0.335 | 0.3230063 |
| LGP-200 | ICC.z | 2352 | 0.330 | 0.3185208 |
| LGP-100 | ICC.z | 2352 | 0.329 | 0.3176219 |
| MMP-360 | ICC.z | 2352 | 0.326 | 0.3149220 |
| LGP-1000 | ICC.z | 2352 | 0.324 | 0.3131193 |

#### 1.5 Descriptive statistics Median

**Table S4.** Parcellation - ICC Median

| parcellation | variable | n | median |
|---|---|---|---|
| wbLGP-558 | ICC | 2352 | 0.662 |
| wbBNP-568 | ICC | 2352 | 0.641 |
| wbLGP-458 | ICC | 2352 | 0.637 |
| wbLGP-858 | ICC | 2352 | 0.637 |
| wbLGP-758 | ICC | 2352 | 0.636 |
| wbLGP-658 | ICC | 2352 | 0.633 |
| wbLGP-1058 | ICC | 2352 | 0.622 |
| wbLGP-1258 | ICC | 2352 | 0.614 |
| wbCABP-718 | ICC | 2352 | 0.613 |
| wbLGP-1158 | ICC | 2352 | 0.613 |
| wbLGP-958 | ICC | 2352 | 0.608 |
| wbLGP-1358 | ICC | 2352 | 0.589 |
| LGP-500 | ICC | 2352 | 0.392 |
| LGP-800 | ICC | 2352 | 0.371 |
| LGP-400 | ICC | 2352 | 0.369 |
| LGP-600 | ICC | 2352 | 0.367 |
| LGP-300 | ICC | 2352 | 0.356 |
| LGP-900 | ICC | 2352 | 0.355 |
| LGP-700 | ICC | 2352 | 0.352 |
| BNP-210 | ICC | 2352 | 0.337 |
| LGP-100 | ICC | 2352 | 0.335 |
| LGP-200 | ICC | 2352 | 0.334 |
| LGP-1000 | ICC | 2352 | 0.329 |
| MMP-360 | ICC | 2352 | 0.314 |

#### 1.6 Friedman Test

**Table S5.** Parcellation - Friedman Test

| .y. | n | statistic | df | p | method |
|---|---|---|---|---|---|
| ICC.z | 2352 | 20379.07 | 23 | 0 | Friedman test |

#### 1.7 Friedman Test Effect size

**Table S6.** Parcellation - Friedman Test Effect size

| .y. | n | effsize | method | magnitude |
|---|---|---|---|---|
| ICC.z | 2352 | 0.3767205 | Kendall W | moderate |

#### 1.8 Paired Wilcoxon signed rank test

**Table S7.** Parcellation - Paired Wilcoxon signed rank test

| group1 | group2 | n1 | statistic | alternative | p | p.adj | p.adj.signif |
|---|---|---|---|---|---|---|---|
| BNP-210 | LGP-100 | 2352 | 1302001 | two.sided | 1.60e-02 | 1.0000000 | ns |
| BNP-210 | LGP-1000 | 2352 | 1379617 | two.sided | 0.00e+00 | 0.0000030 | **** |
| BNP-210 | LGP-200 | 2352 | 1330962 | two.sided | 2.00e-03 | 0.5520000 | ns |
| BNP-210 | LGP-300 | 2352 | 1047149 | two.sided | 0.00e+00 | 0.0000037 | **** |
| BNP-210 | LGP-400 | 2352 | 928029 | two.sided | 0.00e+00 | 0.0000000 | **** |
| BNP-210 | LGP-500 | 2352 | 686156 | two.sided | 0.00e+00 | 0.0000000 | **** |
| BNP-210 | LGP-600 | 2352 | 1004541 | two.sided | 0.00e+00 | 0.0000000 | **** |
| BNP-210 | LGP-700 | 2352 | 1105948 | two.sided | 1.76e-04 | 0.0485760 | * |
| BNP-210 | LGP-800 | 2352 | 945406 | two.sided | 0.00e+00 | 0.0000000 | **** |
| BNP-210 | LGP-900 | 2352 | 1144573 | two.sided | 8.00e-03 | 1.0000000 | ns |
| BNP-210 | MMP-360 | 2352 | 1340035 | two.sided | 2.40e-04 | 0.0662400 | ns |
| BNP-210 | wbBNP-568 | 2352 | 160500 | two.sided | 0.00e+00 | 0.0000000 | **** |
| BNP-210 | wbCABP-718 | 2352 | 162989 | two.sided | 0.00e+00 | 0.0000000 | **** |
| BNP-210 | wbLGP-1058 | 2352 | 176518 | two.sided | 0.00e+00 | 0.0000000 | **** |
| BNP-210 | wbLGP-1158 | 2352 | 199995 | two.sided | 0.00e+00 | 0.0000000 | **** |
| BNP-210 | wbLGP-1258 | 2352 | 180282 | two.sided | 0.00e+00 | 0.0000000 | **** |
| BNP-210 | wbLGP-1358 | 2352 | 232751 | two.sided | 0.00e+00 | 0.0000000 | **** |
| BNP-210 | wbLGP-458 | 2352 | 145336 | two.sided | 0.00e+00 | 0.0000000 | **** |
| BNP-210 | wbLGP-558 | 2352 | 148873 | two.sided | 0.00e+00 | 0.0000000 | **** |
| BNP-210 | wbLGP-658 | 2352 | 175789 | two.sided | 0.00e+00 | 0.0000000 | **** |
| BNP-210 | wbLGP-758 | 2352 | 180496 | two.sided | 0.00e+00 | 0.0000000 | **** |
| BNP-210 | wbLGP-858 | 2352 | 170819 | two.sided | 0.00e+00 | 0.0000000 | **** |
| BNP-210 | wbLGP-958 | 2352 | 189206 | two.sided | 0.00e+00 | 0.0000000 | **** |
| LGP-100 | LGP-1000 | 2352 | 1315278 | two.sided | 6.00e-03 | 1.0000000 | ns |
| LGP-100 | LGP-200 | 2352 | 1255194 | two.sided | 5.26e-01 | 1.0000000 | ns |
| LGP-100 | LGP-300 | 2352 | 1010235 | two.sided | 0.00e+00 | 0.0000000 | **** |
| LGP-100 | LGP-400 | 2352 | 912286 | two.sided | 0.00e+00 | 0.0000000 | **** |
| LGP-100 | LGP-500 | 2352 | 659721 | two.sided | 0.00e+00 | 0.0000000 | **** |
| LGP-100 | LGP-600 | 2352 | 981916 | two.sided | 0.00e+00 | 0.0000000 | **** |
| LGP-100 | LGP-700 | 2352 | 1129951 | two.sided | 4.59e-04 | 0.1266840 | ns |
| LGP-100 | LGP-800 | 2352 | 994724 | two.sided | 0.00e+00 | 0.0000000 | **** |
| LGP-100 | LGP-900 | 2352 | 1156218 | two.sided | 1.40e-02 | 1.0000000 | ns |
| LGP-100 | MMP-360 | 2352 | 1275769 | two.sided | 1.06e-01 | 1.0000000 | ns |
| LGP-100 | wbBNP-568 | 2352 | 159540 | two.sided | 0.00e+00 | 0.0000000 | **** |
| LGP-100 | wbCABP-718 | 2352 | 161073 | two.sided | 0.00e+00 | 0.0000000 | **** |
| LGP-100 | wbLGP-1058 | 2352 | 170565 | two.sided | 0.00e+00 | 0.0000000 | **** |
| LGP-100 | wbLGP-1158 | 2352 | 187780 | two.sided | 0.00e+00 | 0.0000000 | **** |
| LGP-100 | wbLGP-1258 | 2352 | 168428 | two.sided | 0.00e+00 | 0.0000000 | **** |
| LGP-100 | wbLGP-1358 | 2352 | 215782 | two.sided | 0.00e+00 | 0.0000000 | **** |
| LGP-100 | wbLGP-458 | 2352 | 130296 | two.sided | 0.00e+00 | 0.0000000 | **** |

**Table S7.** Parcellation - Paired Wilcoxon signed rank test *(continued)*

| group1 | group2 | n1 | statistic | alternative | p | p.adj | p.adj.signif |
|---|---|---|---|---|---|---|---|
| LGP-100 | wbLGP-558 | 2352 | 136606 | two.sided | 0.00e+00 | 0.0000000 | **** |
| LGP-100 | wbLGP-658 | 2352 | 172094 | two.sided | 0.00e+00 | 0.0000000 | **** |
| LGP-100 | wbLGP-758 | 2352 | 179105 | two.sided | 0.00e+00 | 0.0000000 | **** |
| LGP-100 | wbLGP-858 | 2352 | 167470 | two.sided | 0.00e+00 | 0.0000000 | **** |
| LGP-100 | wbLGP-958 | 2352 | 185024 | two.sided | 0.00e+00 | 0.0000000 | **** |
| LGP-1000 | LGP-200 | 2352 | 1144843 | two.sided | 5.00e-03 | 1.0000000 | ns |
| LGP-1000 | LGP-300 | 2352 | 907429 | two.sided | 0.00e+00 | 0.0000000 | **** |
| LGP-1000 | LGP-400 | 2352 | 756937 | two.sided | 0.00e+00 | 0.0000000 | **** |
| LGP-1000 | LGP-500 | 2352 | 564982 | two.sided | 0.00e+00 | 0.0000000 | **** |
| LGP-1000 | LGP-600 | 2352 | 766240 | two.sided | 0.00e+00 | 0.0000000 | **** |
| LGP-1000 | LGP-700 | 2352 | 813046 | two.sided | 0.00e+00 | 0.0000000 | **** |
| LGP-1000 | LGP-800 | 2352 | 724898 | two.sided | 0.00e+00 | 0.0000000 | **** |
| LGP-1000 | LGP-900 | 2352 | 862540 | two.sided | 0.00e+00 | 0.0000000 | **** |
| LGP-1000 | MMP-360 | 2352 | 1194004 | two.sided | 6.30e-01 | 1.0000000 | ns |
| LGP-1000 | wbBNP-568 | 2352 | 146449 | two.sided | 0.00e+00 | 0.0000000 | **** |
| LGP-1000 | wbCABP-718 | 2352 | 133629 | two.sided | 0.00e+00 | 0.0000000 | **** |
| LGP-1000 | wbLGP-1058 | 2352 | 143488 | two.sided | 0.00e+00 | 0.0000000 | **** |
| LGP-1000 | wbLGP-1158 | 2352 | 152381 | two.sided | 0.00e+00 | 0.0000000 | **** |
| LGP-1000 | wbLGP-1258 | 2352 | 140457 | two.sided | 0.00e+00 | 0.0000000 | **** |
| LGP-1000 | wbLGP-1358 | 2352 | 180243 | two.sided | 0.00e+00 | 0.0000000 | **** |
| LGP-1000 | wbLGP-458 | 2352 | 126403 | two.sided | 0.00e+00 | 0.0000000 | **** |
| LGP-1000 | wbLGP-558 | 2352 | 130477 | two.sided | 0.00e+00 | 0.0000000 | **** |
| LGP-1000 | wbLGP-658 | 2352 | 146959 | two.sided | 0.00e+00 | 0.0000000 | **** |
| LGP-1000 | wbLGP-758 | 2352 | 146560 | two.sided | 0.00e+00 | 0.0000000 | **** |
| LGP-1000 | wbLGP-858 | 2352 | 144497 | two.sided | 0.00e+00 | 0.0000000 | **** |
| LGP-1000 | wbLGP-958 | 2352 | 151257 | two.sided | 0.00e+00 | 0.0000000 | **** |
| LGP-200 | LGP-300 | 2352 | 925471 | two.sided | 0.00e+00 | 0.0000000 | **** |
| LGP-200 | LGP-400 | 2352 | 832641 | two.sided | 0.00e+00 | 0.0000000 | **** |
| LGP-200 | LGP-500 | 2352 | 576900 | two.sided | 0.00e+00 | 0.0000000 | **** |
| LGP-200 | LGP-600 | 2352 | 932917 | two.sided | 0.00e+00 | 0.0000000 | **** |
| LGP-200 | LGP-700 | 2352 | 1091274 | two.sided | 3.40e-06 | 0.0009301 | *** |
| LGP-200 | LGP-800 | 2352 | 917094 | two.sided | 0.00e+00 | 0.0000000 | **** |
| LGP-200 | LGP-900 | 2352 | 1112972 | two.sided | 1.30e-04 | 0.0358800 | * |
| LGP-200 | MMP-360 | 2352 | 1268709 | two.sided | 2.33e-01 | 1.0000000 | ns |
| LGP-200 | wbBNP-568 | 2352 | 144847 | two.sided | 0.00e+00 | 0.0000000 | **** |
| LGP-200 | wbCABP-718 | 2352 | 149133 | two.sided | 0.00e+00 | 0.0000000 | **** |
| LGP-200 | wbLGP-1058 | 2352 | 160610 | two.sided | 0.00e+00 | 0.0000000 | **** |
| LGP-200 | wbLGP-1158 | 2352 | 180943 | two.sided | 0.00e+00 | 0.0000000 | **** |
| LGP-200 | wbLGP-1258 | 2352 | 164714 | two.sided | 0.00e+00 | 0.0000000 | **** |
| LGP-200 | wbLGP-1358 | 2352 | 207338 | two.sided | 0.00e+00 | 0.0000000 | **** |
| LGP-200 | wbLGP-458 | 2352 | 130665 | two.sided | 0.00e+00 | 0.0000000 | **** |

**Table S7.** Parcellation - Paired Wilcoxon signed rank test *(continued)*

| group1 | group2 | n1 | statistic | alternative | p | p.adj | p.adj.signif |
|---|---|---|---|---|---|---|---|
| LGP-200 | wbLGP-558 | 2352 | 131597 | two.sided | 0.00e+00 | 0.0000000 | **** |
| LGP-200 | wbLGP-658 | 2352 | 154264 | two.sided | 0.00e+00 | 0.0000000 | **** |
| LGP-200 | wbLGP-758 | 2352 | 164041 | two.sided | 0.00e+00 | 0.0000000 | **** |
| LGP-200 | wbLGP-858 | 2352 | 159660 | two.sided | 0.00e+00 | 0.0000000 | **** |
| LGP-200 | wbLGP-958 | 2352 | 176411 | two.sided | 0.00e+00 | 0.0000000 | **** |
| LGP-300 | LGP-400 | 2352 | 999180 | two.sided | 0.00e+00 | 0.0000000 | **** |
| LGP-300 | LGP-500 | 2352 | 629869 | two.sided | 0.00e+00 | 0.0000000 | **** |
| LGP-300 | LGP-600 | 2352 | 1060070 | two.sided | 2.00e-07 | 0.0000602 | **** |
| LGP-300 | LGP-700 | 2352 | 1259128 | two.sided | 1.03e-01 | 1.0000000 | ns |
| LGP-300 | LGP-800 | 2352 | 1069448 | two.sided | 2.00e-07 | 0.0000538 | **** |
| LGP-300 | LGP-900 | 2352 | 1299549 | two.sided | 6.00e-03 | 1.0000000 | ns |
| LGP-300 | MMP-360 | 2352 | 1483967 | two.sided | 0.00e+00 | 0.0000000 | **** |
| LGP-300 | wbBNP-568 | 2352 | 164589 | two.sided | 0.00e+00 | 0.0000000 | **** |
| LGP-300 | wbCABP-718 | 2352 | 158079 | two.sided | 0.00e+00 | 0.0000000 | **** |
| LGP-300 | wbLGP-1058 | 2352 | 172650 | two.sided | 0.00e+00 | 0.0000000 | **** |
| LGP-300 | wbLGP-1158 | 2352 | 196899 | two.sided | 0.00e+00 | 0.0000000 | **** |
| LGP-300 | wbLGP-1258 | 2352 | 176738 | two.sided | 0.00e+00 | 0.0000000 | **** |
| LGP-300 | wbLGP-1358 | 2352 | 227257 | two.sided | 0.00e+00 | 0.0000000 | **** |
| LGP-300 | wbLGP-458 | 2352 | 143518 | two.sided | 0.00e+00 | 0.0000000 | **** |
| LGP-300 | wbLGP-558 | 2352 | 144041 | two.sided | 0.00e+00 | 0.0000000 | **** |
| LGP-300 | wbLGP-658 | 2352 | 173281 | two.sided | 0.00e+00 | 0.0000000 | **** |
| LGP-300 | wbLGP-758 | 2352 | 170528 | two.sided | 0.00e+00 | 0.0000000 | **** |
| LGP-300 | wbLGP-858 | 2352 | 163732 | two.sided | 0.00e+00 | 0.0000000 | **** |
| LGP-300 | wbLGP-958 | 2352 | 186563 | two.sided | 0.00e+00 | 0.0000000 | **** |
| LGP-400 | LGP-500 | 2352 | 767221 | two.sided | 0.00e+00 | 0.0000000 | **** |
| LGP-400 | LGP-600 | 2352 | 1251633 | two.sided | 2.34e-01 | 1.0000000 | ns |
| LGP-400 | LGP-700 | 2352 | 1517947 | two.sided | 0.00e+00 | 0.0000000 | **** |
| LGP-400 | LGP-800 | 2352 | 1303871 | two.sided | 1.20e-02 | 1.0000000 | ns |
| LGP-400 | LGP-900 | 2352 | 1477786 | two.sided | 0.00e+00 | 0.0000000 | **** |
| LGP-400 | MMP-360 | 2352 | 1583190 | two.sided | 0.00e+00 | 0.0000000 | **** |
| LGP-400 | wbBNP-568 | 2352 | 178030 | two.sided | 0.00e+00 | 0.0000000 | **** |
| LGP-400 | wbCABP-718 | 2352 | 168997 | two.sided | 0.00e+00 | 0.0000000 | **** |
| LGP-400 | wbLGP-1058 | 2352 | 190089 | two.sided | 0.00e+00 | 0.0000000 | **** |
| LGP-400 | wbLGP-1158 | 2352 | 212433 | two.sided | 0.00e+00 | 0.0000000 | **** |
| LGP-400 | wbLGP-1258 | 2352 | 188919 | two.sided | 0.00e+00 | 0.0000000 | **** |
| LGP-400 | wbLGP-1358 | 2352 | 246801 | two.sided | 0.00e+00 | 0.0000000 | **** |
| LGP-400 | wbLGP-458 | 2352 | 151198 | two.sided | 0.00e+00 | 0.0000000 | **** |
| LGP-400 | wbLGP-558 | 2352 | 159132 | two.sided | 0.00e+00 | 0.0000000 | **** |
| LGP-400 | wbLGP-658 | 2352 | 190283 | two.sided | 0.00e+00 | 0.0000000 | **** |
| LGP-400 | wbLGP-758 | 2352 | 185190 | two.sided | 0.00e+00 | 0.0000000 | **** |
| LGP-400 | wbLGP-858 | 2352 | 182103 | two.sided | 0.00e+00 | 0.0000000 | **** |
| LGP-400 | wbLGP-958 | 2352 | 201977 | two.sided | 0.00e+00 | 0.0000000 | **** |

**Table S7.** Parcellation - Paired Wilcoxon signed rank test *(continued)*

| group1 | group2 | n1 | statistic | alternative | p | p.adj | p.adj.signif |
|---|---|---|---|---|---|---|---|
| LGP-500 | LGP-600 | 2352 | 1636990 | two.sided | 0.00e+00 | 0.0000000 | **** |
| LGP-500 | LGP-700 | 2352 | 1775407 | two.sided | 0.00e+00 | 0.0000000 | **** |
| LGP-500 | LGP-800 | 2352 | 1732442 | two.sided | 0.00e+00 | 0.0000000 | **** |
| LGP-500 | LGP-900 | 2352 | 1731306 | two.sided | 0.00e+00 | 0.0000000 | **** |
| LGP-500 | MMP-360 | 2352 | 1758329 | two.sided | 0.00e+00 | 0.0000000 | **** |
| LGP-500 | wbBNP-568 | 2352 | 195928 | two.sided | 0.00e+00 | 0.0000000 | **** |
| LGP-500 | wbCABP-718 | 2352 | 187217 | two.sided | 0.00e+00 | 0.0000000 | **** |
| LGP-500 | wbLGP-1058 | 2352 | 202651 | two.sided | 0.00e+00 | 0.0000000 | **** |
| LGP-500 | wbLGP-1158 | 2352 | 224307 | two.sided | 0.00e+00 | 0.0000000 | **** |
| LGP-500 | wbLGP-1258 | 2352 | 206925 | two.sided | 0.00e+00 | 0.0000000 | **** |
| LGP-500 | wbLGP-1358 | 2352 | 273700 | two.sided | 0.00e+00 | 0.0000000 | **** |
| LGP-500 | wbLGP-458 | 2352 | 179170 | two.sided | 0.00e+00 | 0.0000000 | **** |
| LGP-500 | wbLGP-558 | 2352 | 167854 | two.sided | 0.00e+00 | 0.0000000 | **** |
| LGP-500 | wbLGP-658 | 2352 | 207577 | two.sided | 0.00e+00 | 0.0000000 | **** |
| LGP-500 | wbLGP-758 | 2352 | 205183 | two.sided | 0.00e+00 | 0.0000000 | **** |
| LGP-500 | wbLGP-858 | 2352 | 190097 | two.sided | 0.00e+00 | 0.0000000 | **** |
| LGP-500 | wbLGP-958 | 2352 | 223872 | two.sided | 0.00e+00 | 0.0000000 | **** |
| LGP-600 | LGP-700 | 2352 | 1517047 | two.sided | 0.00e+00 | 0.0000000 | **** |
| LGP-600 | LGP-800 | 2352 | 1268374 | two.sided | 2.70e-02 | 1.0000000 | ns |
| LGP-600 | LGP-900 | 2352 | 1454702 | two.sided | 0.00e+00 | 0.0000000 | **** |
| LGP-600 | MMP-360 | 2352 | 1486050 | two.sided | 0.00e+00 | 0.0000000 | **** |
| LGP-600 | wbBNP-568 | 2352 | 151395 | two.sided | 0.00e+00 | 0.0000000 | **** |
| LGP-600 | wbCABP-718 | 2352 | 146032 | two.sided | 0.00e+00 | 0.0000000 | **** |
| LGP-600 | wbLGP-1058 | 2352 | 151532 | two.sided | 0.00e+00 | 0.0000000 | **** |
| LGP-600 | wbLGP-1158 | 2352 | 174886 | two.sided | 0.00e+00 | 0.0000000 | **** |
| LGP-600 | wbLGP-1258 | 2352 | 164221 | two.sided | 0.00e+00 | 0.0000000 | **** |
| LGP-600 | wbLGP-1358 | 2352 | 210481 | two.sided | 0.00e+00 | 0.0000000 | **** |
| LGP-600 | wbLGP-458 | 2352 | 142445 | two.sided | 0.00e+00 | 0.0000000 | **** |
| LGP-600 | wbLGP-558 | 2352 | 138450 | two.sided | 0.00e+00 | 0.0000000 | **** |
| LGP-600 | wbLGP-658 | 2352 | 156775 | two.sided | 0.00e+00 | 0.0000000 | **** |
| LGP-600 | wbLGP-758 | 2352 | 158441 | two.sided | 0.00e+00 | 0.0000000 | **** |
| LGP-600 | wbLGP-858 | 2352 | 148288 | two.sided | 0.00e+00 | 0.0000000 | **** |
| LGP-600 | wbLGP-958 | 2352 | 145886 | two.sided | 0.00e+00 | 0.0000000 | **** |
| LGP-700 | LGP-800 | 2352 | 994539 | two.sided | 0.00e+00 | 0.0000000 | **** |
| LGP-700 | LGP-900 | 2352 | 1232646 | two.sided | 3.15e-01 | 1.0000000 | ns |
| LGP-700 | MMP-360 | 2352 | 1426998 | two.sided | 0.00e+00 | 0.0000000 | **** |
| LGP-700 | wbBNP-568 | 2352 | 161398 | two.sided | 0.00e+00 | 0.0000000 | **** |
| LGP-700 | wbCABP-718 | 2352 | 152205 | two.sided | 0.00e+00 | 0.0000000 | **** |
| LGP-700 | wbLGP-1058 | 2352 | 163976 | two.sided | 0.00e+00 | 0.0000000 | **** |
| LGP-700 | wbLGP-1158 | 2352 | 185951 | two.sided | 0.00e+00 | 0.0000000 | **** |
| LGP-700 | wbLGP-1258 | 2352 | 167287 | two.sided | 0.00e+00 | 0.0000000 | **** |
| LGP-700 | wbLGP-1358 | 2352 | 219346 | two.sided | 0.00e+00 | 0.0000000 | **** |

**Table S7.** Parcellation - Paired Wilcoxon signed rank test *(continued)*

| group1 | group2 | n1 | statistic | alternative | p | p.adj | p.adj.signif |
|---|---|---|---|---|---|---|---|
| LGP-700 | wbLGP-458 | 2352 | 142001 | two.sided | 0.00e+00 | 0.0000000 | **** |
| LGP-700 | wbLGP-558 | 2352 | 145607 | two.sided | 0.00e+00 | 0.0000000 | **** |
| LGP-700 | wbLGP-658 | 2352 | 169299 | two.sided | 0.00e+00 | 0.0000000 | **** |
| LGP-700 | wbLGP-758 | 2352 | 169104 | two.sided | 0.00e+00 | 0.0000000 | **** |
| LGP-700 | wbLGP-858 | 2352 | 164010 | two.sided | 0.00e+00 | 0.0000000 | **** |
| LGP-700 | wbLGP-958 | 2352 | 173569 | two.sided | 0.00e+00 | 0.0000000 | **** |
| LGP-800 | LGP-900 | 2352 | 1418708 | two.sided | 0.00e+00 | 0.0000000 | **** |
| LGP-800 | MMP-360 | 2352 | 1553788 | two.sided | 0.00e+00 | 0.0000000 | **** |
| LGP-800 | wbBNP-568 | 2352 | 181300 | two.sided | 0.00e+00 | 0.0000000 | **** |
| LGP-800 | wbCABP-718 | 2352 | 172356 | two.sided | 0.00e+00 | 0.0000000 | **** |
| LGP-800 | wbLGP-1058 | 2352 | 188254 | two.sided | 0.00e+00 | 0.0000000 | **** |
| LGP-800 | wbLGP-1158 | 2352 | 193741 | two.sided | 0.00e+00 | 0.0000000 | **** |
| LGP-800 | wbLGP-1258 | 2352 | 188270 | two.sided | 0.00e+00 | 0.0000000 | **** |
| LGP-800 | wbLGP-1358 | 2352 | 242146 | two.sided | 0.00e+00 | 0.0000000 | **** |
| LGP-800 | wbLGP-458 | 2352 | 162709 | two.sided | 0.00e+00 | 0.0000000 | **** |
| LGP-800 | wbLGP-558 | 2352 | 160927 | two.sided | 0.00e+00 | 0.0000000 | **** |
| LGP-800 | wbLGP-658 | 2352 | 189416 | two.sided | 0.00e+00 | 0.0000000 | **** |
| LGP-800 | wbLGP-758 | 2352 | 190433 | two.sided | 0.00e+00 | 0.0000000 | **** |
| LGP-800 | wbLGP-858 | 2352 | 183382 | two.sided | 0.00e+00 | 0.0000000 | **** |
| LGP-800 | wbLGP-958 | 2352 | 201777 | two.sided | 0.00e+00 | 0.0000000 | **** |
| LGP-900 | MMP-360 | 2352 | 1411598 | two.sided | 0.00e+00 | 0.0000000 | **** |
| LGP-900 | wbBNP-568 | 2352 | 161762 | two.sided | 0.00e+00 | 0.0000000 | **** |
| LGP-900 | wbCABP-718 | 2352 | 150635 | two.sided | 0.00e+00 | 0.0000000 | **** |
| LGP-900 | wbLGP-1058 | 2352 | 167674 | two.sided | 0.00e+00 | 0.0000000 | **** |
| LGP-900 | wbLGP-1158 | 2352 | 175152 | two.sided | 0.00e+00 | 0.0000000 | **** |
| LGP-900 | wbLGP-1258 | 2352 | 162633 | two.sided | 0.00e+00 | 0.0000000 | **** |
| LGP-900 | wbLGP-1358 | 2352 | 210734 | two.sided | 0.00e+00 | 0.0000000 | **** |
| LGP-900 | wbLGP-458 | 2352 | 143322 | two.sided | 0.00e+00 | 0.0000000 | **** |
| LGP-900 | wbLGP-558 | 2352 | 145640 | two.sided | 0.00e+00 | 0.0000000 | **** |
| LGP-900 | wbLGP-658 | 2352 | 166904 | two.sided | 0.00e+00 | 0.0000000 | **** |
| LGP-900 | wbLGP-758 | 2352 | 172439 | two.sided | 0.00e+00 | 0.0000000 | **** |
| LGP-900 | wbLGP-858 | 2352 | 161857 | two.sided | 0.00e+00 | 0.0000000 | **** |
| LGP-900 | wbLGP-958 | 2352 | 177763 | two.sided | 0.00e+00 | 0.0000000 | **** |
| MMP-360 | wbBNP-568 | 2352 | 153083 | two.sided | 0.00e+00 | 0.0000000 | **** |
| MMP-360 | wbCABP-718 | 2352 | 137815 | two.sided | 0.00e+00 | 0.0000000 | **** |
| MMP-360 | wbLGP-1058 | 2352 | 158332 | two.sided | 0.00e+00 | 0.0000000 | **** |
| MMP-360 | wbLGP-1158 | 2352 | 185386 | two.sided | 0.00e+00 | 0.0000000 | **** |
| MMP-360 | wbLGP-1258 | 2352 | 166968 | two.sided | 0.00e+00 | 0.0000000 | **** |
| MMP-360 | wbLGP-1358 | 2352 | 219183 | two.sided | 0.00e+00 | 0.0000000 | **** |
| MMP-360 | wbLGP-458 | 2352 | 137964 | two.sided | 0.00e+00 | 0.0000000 | **** |
| MMP-360 | wbLGP-558 | 2352 | 138865 | two.sided | 0.00e+00 | 0.0000000 | **** |

**Table S7.** Parcellation - Paired Wilcoxon signed rank test *(continued)*

| group1 | group2 | n1 | statistic | alternative | p | p.adj | p.adj.signif |
|---|---|---|---|---|---|---|---|
| MMP-360 | wbLGP-658 | 2352 | 156009 | two.sided | 0.00e+00 | 0.0000000 | **** |
| MMP-360 | wbLGP-758 | 2352 | 164978 | two.sided | 0.00e+00 | 0.0000000 | **** |
| MMP-360 | wbLGP-858 | 2352 | 157870 | two.sided | 0.00e+00 | 0.0000000 | **** |
| MMP-360 | wbLGP-958 | 2352 | 182605 | two.sided | 0.00e+00 | 0.0000000 | **** |
| wbBNP-568 | wbCABP-718 | 2352 | 1639241 | two.sided | 0.00e+00 | 0.0000000 | **** |
| wbBNP-568 | wbLGP-1058 | 2352 | 1547859 | two.sided | 0.00e+00 | 0.0000000 | **** |
| wbBNP-568 | wbLGP-1158 | 2352 | 1533796 | two.sided | 0.00e+00 | 0.0000000 | **** |
| wbBNP-568 | wbLGP-1258 | 2352 | 1447228 | two.sided | 0.00e+00 | 0.0000000 | **** |
| wbBNP-568 | wbLGP-1358 | 2352 | 1650972 | two.sided | 0.00e+00 | 0.0000000 | **** |
| wbBNP-568 | wbLGP-458 | 2352 | 1125170 | two.sided | 1.22e-05 | 0.0033672 | ** |
| wbBNP-568 | wbLGP-558 | 2352 | 1098196 | two.sided | 2.40e-06 | 0.0006486 | *** |
| wbBNP-568 | wbLGP-658 | 2352 | 1550059 | two.sided | 0.00e+00 | 0.0000000 | **** |
| wbBNP-568 | wbLGP-758 | 2352 | 1595774 | two.sided | 0.00e+00 | 0.0000000 | **** |
| wbBNP-568 | wbLGP-858 | 2352 | 1458694 | two.sided | 0.00e+00 | 0.0000000 | **** |
| wbBNP-568 | wbLGP-958 | 2352 | 1661566 | two.sided | 0.00e+00 | 0.0000000 | **** |
| wbCABP-718 | wbLGP-1058 | 2352 | 1166229 | two.sided | 8.90e-02 | 1.0000000 | ns |
| wbCABP-718 | wbLGP-1158 | 2352 | 1266087 | two.sided | 3.20e-01 | 1.0000000 | ns |
| wbCABP-718 | wbLGP-1258 | 2352 | 1176085 | two.sided | 5.60e-02 | 1.0000000 | ns |
| wbCABP-718 | wbLGP-1358 | 2352 | 1400531 | two.sided | 1.00e-07 | 0.0000293 | **** |
| wbCABP-718 | wbLGP-458 | 2352 | 831718 | two.sided | 0.00e+00 | 0.0000000 | **** |
| wbCABP-718 | wbLGP-558 | 2352 | 757451 | two.sided | 0.00e+00 | 0.0000000 | **** |
| wbCABP-718 | wbLGP-658 | 2352 | 1045849 | two.sided | 0.00e+00 | 0.0000006 | **** |
| wbCABP-718 | wbLGP-758 | 2352 | 1166760 | two.sided | 5.80e-02 | 1.0000000 | ns |
| wbCABP-718 | wbLGP-858 | 2352 | 1068928 | two.sided | 4.00e-07 | 0.0001190 | *** |
| wbCABP-718 | wbLGP-958 | 2352 | 1278722 | two.sided | 8.60e-02 | 1.0000000 | ns |
| wbLGP-1058 | wbLGP-1158 | 2352 | 1310431 | two.sided | 1.00e-02 | 1.0000000 | ns |
| wbLGP-1058 | wbLGP-1258 | 2352 | 1068697 | two.sided | 1.00e-07 | 0.0000157 | **** |
| wbLGP-1058 | wbLGP-1358 | 2352 | 1503855 | two.sided | 0.00e+00 | 0.0000000 | **** |
| wbLGP-1058 | wbLGP-458 | 2352 | 921542 | two.sided | 0.00e+00 | 0.0000000 | **** |
| wbLGP-1058 | wbLGP-558 | 2352 | 830108 | two.sided | 0.00e+00 | 0.0000000 | **** |
| wbLGP-1058 | wbLGP-658 | 2352 | 1070610 | two.sided | 3.00e-07 | 0.0000944 | **** |
| wbLGP-1058 | wbLGP-758 | 2352 | 1177300 | two.sided | 2.97e-01 | 1.0000000 | ns |
| wbLGP-1058 | wbLGP-858 | 2352 | 975194 | two.sided | 0.00e+00 | 0.0000000 | **** |
| wbLGP-1058 | wbLGP-958 | 2352 | 1341281 | two.sided | 5.59e-05 | 0.0154284 | * |
| wbLGP-1158 | wbLGP-1258 | 2352 | 1053122 | two.sided | 0.00e+00 | 0.0000005 | **** |
| wbLGP-1158 | wbLGP-1358 | 2352 | 1451123 | two.sided | 0.00e+00 | 0.0000000 | **** |
| wbLGP-1158 | wbLGP-458 | 2352 | 908290 | two.sided | 0.00e+00 | 0.0000000 | **** |
| wbLGP-1158 | wbLGP-558 | 2352 | 892111 | two.sided | 0.00e+00 | 0.0000000 | **** |
| wbLGP-1158 | wbLGP-658 | 2352 | 1078962 | two.sided | 3.00e-07 | 0.0000836 | **** |
| wbLGP-1158 | wbLGP-758 | 2352 | 1178685 | two.sided | 1.16e-01 | 1.0000000 | ns |
| wbLGP-1158 | wbLGP-858 | 2352 | 1008073 | two.sided | 0.00e+00 | 0.0000000 | **** |
| wbLGP-1158 | wbLGP-958 | 2352 | 1295606 | two.sided | 3.40e-02 | 1.0000000 | ns |

**Table S7.** Parcellation - Paired Wilcoxon signed rank test *(continued)*

| group1 | group2 | n1 | statistic | alternative | p | p.adj | p.adj.signif |
|---|---|---|---|---|---|---|---|
| wbLGP-1258 | wbLGP-1358 | 2352 | 1612256 | two.sided | 0.00e+00 | 0.0000000 | **** |
| wbLGP-1258 | wbLGP-458 | 2352 | 994969 | two.sided | 0.00e+00 | 0.0000000 | **** |
| wbLGP-1258 | wbLGP-558 | 2352 | 957447 | two.sided | 0.00e+00 | 0.0000000 | **** |
| wbLGP-1258 | wbLGP-658 | 2352 | 1172824 | two.sided | 4.40e-02 | 1.0000000 | ns |
| wbLGP-1258 | wbLGP-758 | 2352 | 1273752 | two.sided | 1.74e-01 | 1.0000000 | ns |
| wbLGP-1258 | wbLGP-858 | 2352 | 1150313 | two.sided | 8.00e-03 | 1.0000000 | ns |
| wbLGP-1258 | wbLGP-958 | 2352 | 1390511 | two.sided | 1.00e-07 | 0.0000197 | **** |
| wbLGP-1358 | wbLGP-458 | 2352 | 824258 | two.sided | 0.00e+00 | 0.0000000 | **** |
| wbLGP-1358 | wbLGP-558 | 2352 | 808992 | two.sided | 0.00e+00 | 0.0000000 | **** |
| wbLGP-1358 | wbLGP-658 | 2352 | 968209 | two.sided | 0.00e+00 | 0.0000000 | **** |
| wbLGP-1358 | wbLGP-758 | 2352 | 1031212 | two.sided | 0.00e+00 | 0.0000000 | **** |
| wbLGP-1358 | wbLGP-858 | 2352 | 936260 | two.sided | 0.00e+00 | 0.0000000 | **** |
| wbLGP-1358 | wbLGP-958 | 2352 | 1134304 | two.sided | 6.89e-04 | 0.1901640 | ns |
| wbLGP-458 | wbLGP-558 | 2352 | 1264741 | two.sided | 5.63e-01 | 1.0000000 | ns |
| wbLGP-458 | wbLGP-658 | 2352 | 1598752 | two.sided | 0.00e+00 | 0.0000000 | **** |
| wbLGP-458 | wbLGP-758 | 2352 | 1629670 | two.sided | 0.00e+00 | 0.0000000 | **** |
| wbLGP-458 | wbLGP-858 | 2352 | 1517308 | two.sided | 0.00e+00 | 0.0000000 | **** |
| wbLGP-458 | wbLGP-958 | 2352 | 1684990 | two.sided | 0.00e+00 | 0.0000000 | **** |
| wbLGP-558 | wbLGP-658 | 2352 | 1702013 | two.sided | 0.00e+00 | 0.0000000 | **** |
| wbLGP-558 | wbLGP-758 | 2352 | 1693929 | two.sided | 0.00e+00 | 0.0000000 | **** |
| wbLGP-558 | wbLGP-858 | 2352 | 1557712 | two.sided | 0.00e+00 | 0.0000000 | **** |
| wbLGP-558 | wbLGP-958 | 2352 | 1716972 | two.sided | 0.00e+00 | 0.0000000 | **** |
| wbLGP-658 | wbLGP-758 | 2352 | 1388101 | two.sided | 0.00e+00 | 0.0000049 | **** |
| wbLGP-658 | wbLGP-858 | 2352 | 1241424 | two.sided | 5.82e-01 | 1.0000000 | ns |
| wbLGP-658 | wbLGP-958 | 2352 | 1515613 | two.sided | 0.00e+00 | 0.0000000 | **** |
| wbLGP-758 | wbLGP-858 | 2352 | 970045 | two.sided | 0.00e+00 | 0.0000000 | **** |
| wbLGP-758 | wbLGP-958 | 2352 | 1351987 | two.sided | 3.70e-06 | 0.0010295 | ** |
| wbLGP-858 | wbLGP-958 | 2352 | 1494529 | two.sided | 0.00e+00 | 0.0000000 | **** |

#### 1.9 Kullback-leibler divergence Map

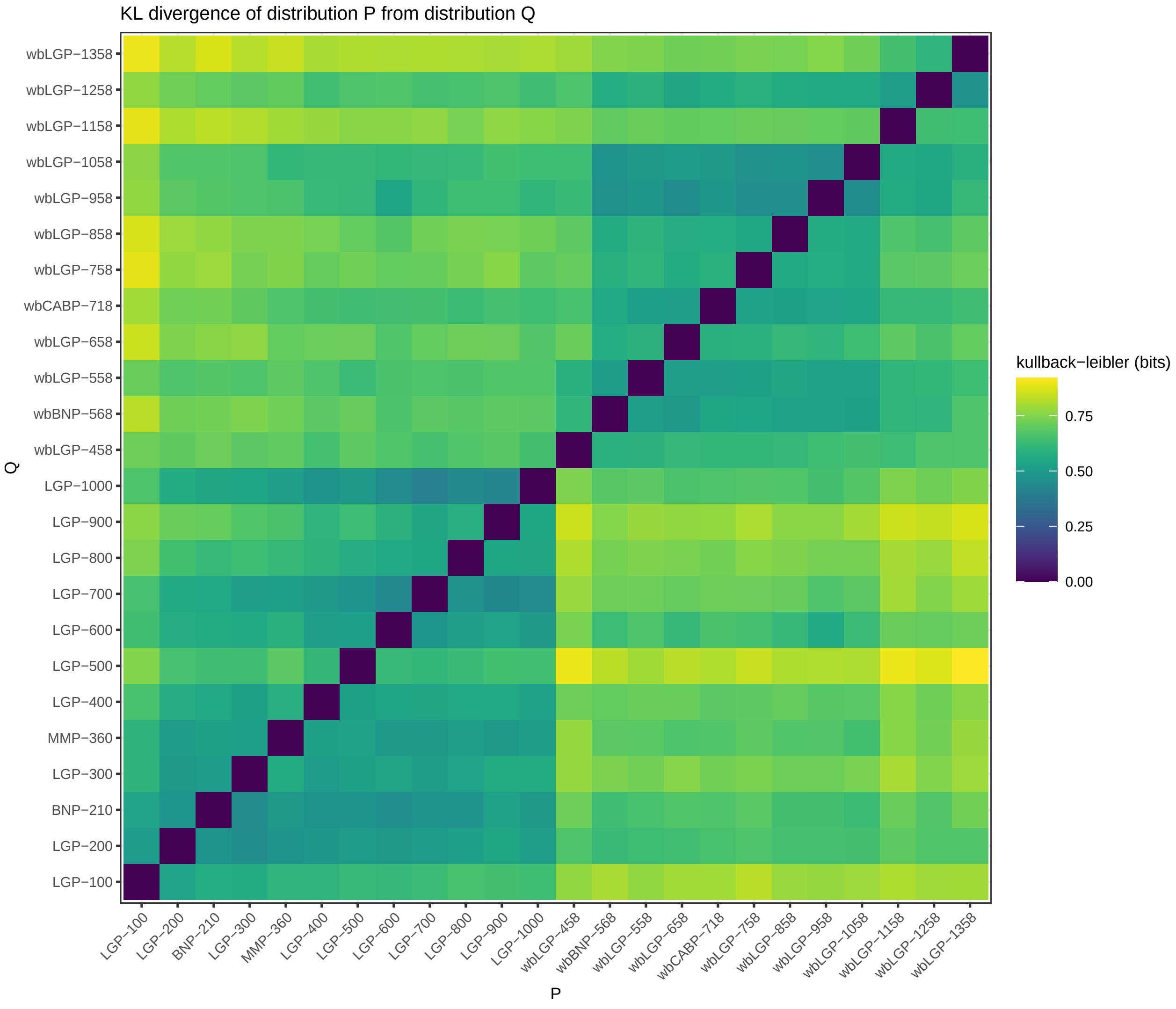

**Fig. S4.** Parcellation - KL divergence Map

#### 1.10 Significance Map

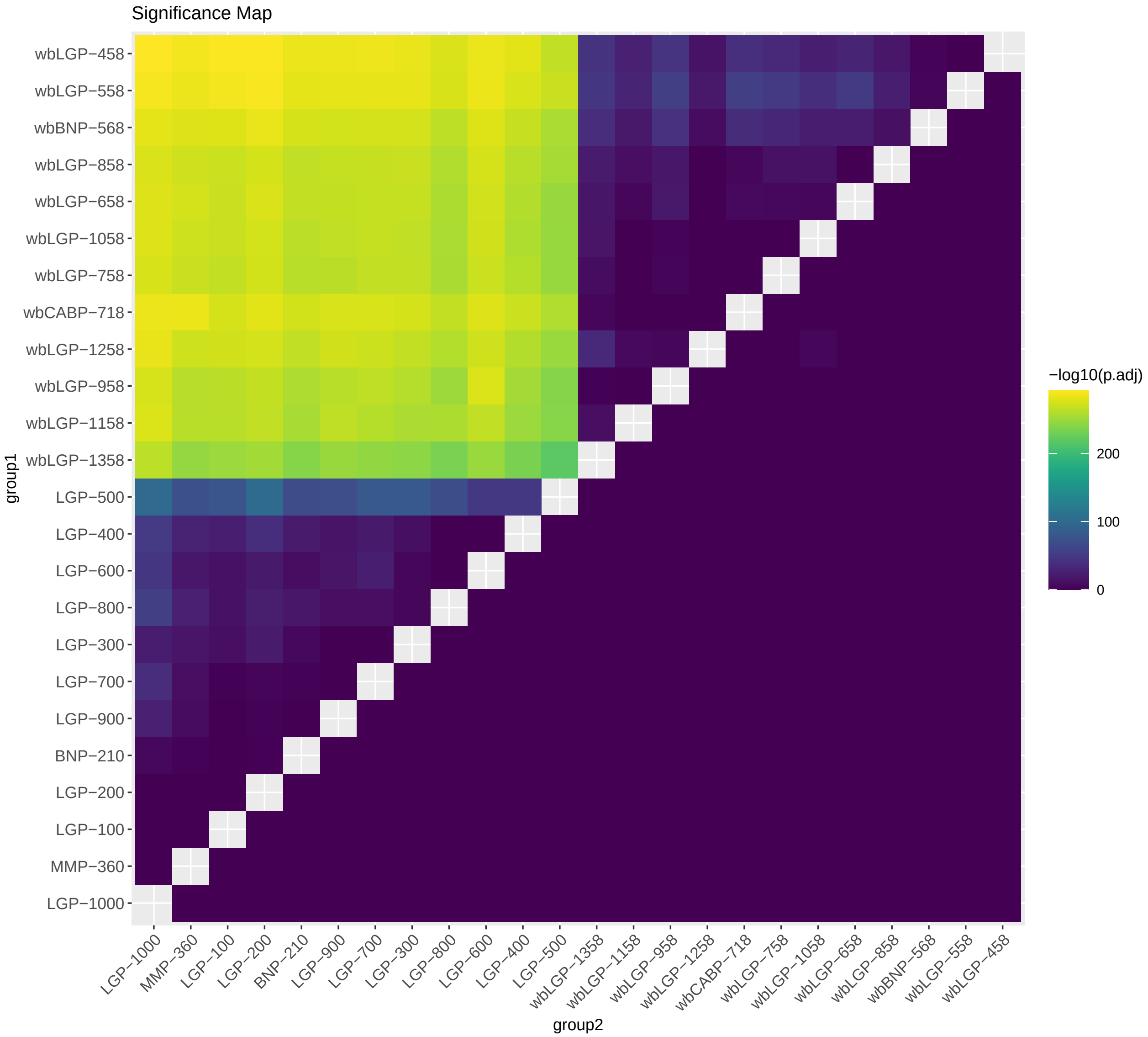

**Fig. S5.** Parcellation - Significance Map

#### 1.11 Effect size

**Table S8.** Parcellation - Effect size

| group1 | group2 | effsize | n1 | n2 | magnitude |
|---|---|---|---|---|---|
| BNP-210 | LGP-100 | 0.0512884 | 2352 | 2352 | small |
| BNP-210 | LGP-1000 | 0.1207652 | 2352 | 2352 | small |
| BNP-210 | LGP-200 | 0.0626895 | 2352 | 2352 | small |
| BNP-210 | LGP-300 | 0.1141191 | 2352 | 2352 | small |
| BNP-210 | LGP-400 | 0.2082967 | 2352 | 2352 | small |
| BNP-210 | LGP-500 | 0.3653563 | 2352 | 2352 | moderate |
| BNP-210 | LGP-600 | 0.1408717 | 2352 | 2352 | small |

**Table S8.** Parcellation - Effect size *(continued)*

| group1 | group2 | effsize | n1 | n2 | magnitude |
|---|---|---|---|---|---|
| BNP-210 | LGP-700 | 0.0779897 | 2352 | 2352 | small |
| BNP-210 | LGP-800 | 0.1893357 | 2352 | 2352 | small |
| BNP-210 | LGP-900 | 0.0543280 | 2352 | 2352 | small |
| BNP-210 | MMP-360 | 0.0731563 | 2352 | 2352 | small |
| BNP-210 | wbBNP-568 | 0.7389896 | 2352 | 2352 | large |
| BNP-210 | wbCABP-718 | 0.7379236 | 2352 | 2352 | large |
| BNP-210 | wbLGP-1058 | 0.7216681 | 2352 | 2352 | large |
| BNP-210 | wbLGP-1158 | 0.7104950 | 2352 | 2352 | large |
| BNP-210 | wbLGP-1258 | 0.7231289 | 2352 | 2352 | large |
| BNP-210 | wbLGP-1358 | 0.6888160 | 2352 | 2352 | large |
| BNP-210 | wbLGP-458 | 0.7536559 | 2352 | 2352 | large |
| BNP-210 | wbLGP-558 | 0.7475750 | 2352 | 2352 | large |
| BNP-210 | wbLGP-658 | 0.7266246 | 2352 | 2352 | large |
| BNP-210 | wbLGP-758 | 0.7214303 | 2352 | 2352 | large |
| BNP-210 | wbLGP-858 | 0.7258208 | 2352 | 2352 | large |
| BNP-210 | wbLGP-958 | 0.7141604 | 2352 | 2352 | large |
| LGP-100 | LGP-1000 | 0.0539583 | 2352 | 2352 | small |
| LGP-100 | LGP-200 | 0.0129528 | 2352 | 2352 | small |
| LGP-100 | LGP-300 | 0.1500861 | 2352 | 2352 | small |
| LGP-100 | LGP-400 | 0.2208211 | 2352 | 2352 | small |
| LGP-100 | LGP-500 | 0.3913895 | 2352 | 2352 | moderate |
| LGP-100 | LGP-600 | 0.1695900 | 2352 | 2352 | small |
| LGP-100 | LGP-700 | 0.0698175 | 2352 | 2352 | small |
| LGP-100 | LGP-800 | 0.1689070 | 2352 | 2352 | small |
| LGP-100 | LGP-900 | 0.0495689 | 2352 | 2352 | small |
| LGP-100 | MMP-360 | 0.0317933 | 2352 | 2352 | small |
| LGP-100 | wbBNP-568 | 0.7436624 | 2352 | 2352 | large |
| LGP-100 | wbCABP-718 | 0.7394118 | 2352 | 2352 | large |
| LGP-100 | wbLGP-1058 | 0.7296547 | 2352 | 2352 | large |
| LGP-100 | wbLGP-1158 | 0.7220587 | 2352 | 2352 | large |
| LGP-100 | wbLGP-1258 | 0.7355677 | 2352 | 2352 | large |
| LGP-100 | wbLGP-1358 | 0.7044764 | 2352 | 2352 | large |
| LGP-100 | wbLGP-458 | 0.7640762 | 2352 | 2352 | large |
| LGP-100 | wbLGP-558 | 0.7569985 | 2352 | 2352 | large |
| LGP-100 | wbLGP-658 | 0.7310191 | 2352 | 2352 | large |
| LGP-100 | wbLGP-758 | 0.7275245 | 2352 | 2352 | large |
| LGP-100 | wbLGP-858 | 0.7326421 | 2352 | 2352 | large |
| LGP-100 | wbLGP-958 | 0.7209303 | 2352 | 2352 | large |
| LGP-1000 | LGP-200 | 0.0578365 | 2352 | 2352 | small |
| LGP-1000 | LGP-300 | 0.2105271 | 2352 | 2352 | small |
| LGP-1000 | LGP-400 | 0.3192842 | 2352 | 2352 | moderate |
| LGP-1000 | LGP-500 | 0.4487333 | 2352 | 2352 | moderate |
| LGP-1000 | LGP-600 | 0.3020832 | 2352 | 2352 | moderate |
| LGP-1000 | LGP-700 | 0.2719757 | 2352 | 2352 | small |
| LGP-1000 | LGP-800 | 0.3321805 | 2352 | 2352 | moderate |
| LGP-1000 | LGP-900 | 0.2340567 | 2352 | 2352 | small |
| LGP-1000 | MMP-360 | 0.0081145 | 2352 | 2352 | small |

**Table S8.** Parcellation - Effect size *(continued)*

| group1 | group2 | effsize | n1 | n2 | magnitude |
|---|---|---|---|---|---|
| LGP-1000 | wbBNP-568 | 0.7500285 | 2352 | 2352 | large |
| LGP-1000 | wbCABP-718 | 0.7568211 | 2352 | 2352 | large |
| LGP-1000 | wbLGP-1058 | 0.7435657 | 2352 | 2352 | large |
| LGP-1000 | wbLGP-1158 | 0.7443931 | 2352 | 2352 | large |
| LGP-1000 | wbLGP-1258 | 0.7509503 | 2352 | 2352 | large |
| LGP-1000 | wbLGP-1358 | 0.7255476 | 2352 | 2352 | large |
| LGP-1000 | wbLGP-458 | 0.7662544 | 2352 | 2352 | large |
| LGP-1000 | wbLGP-558 | 0.7623347 | 2352 | 2352 | large |
| LGP-1000 | wbLGP-658 | 0.7456084 | 2352 | 2352 | large |
| LGP-1000 | wbLGP-758 | 0.7432770 | 2352 | 2352 | large |
| LGP-1000 | wbLGP-858 | 0.7434569 | 2352 | 2352 | large |
| LGP-1000 | wbLGP-958 | 0.7395782 | 2352 | 2352 | large |
| LGP-200 | LGP-300 | 0.2081011 | 2352 | 2352 | small |
| LGP-200 | LGP-400 | 0.2759045 | 2352 | 2352 | small |
| LGP-200 | LGP-500 | 0.4511007 | 2352 | 2352 | moderate |
| LGP-200 | LGP-600 | 0.1986975 | 2352 | 2352 | small |
| LGP-200 | LGP-700 | 0.0956037 | 2352 | 2352 | small |
| LGP-200 | LGP-800 | 0.2215336 | 2352 | 2352 | small |
| LGP-200 | LGP-900 | 0.0807512 | 2352 | 2352 | small |
| LGP-200 | MMP-360 | 0.0232331 | 2352 | 2352 | small |
| LGP-200 | wbBNP-568 | 0.7524502 | 2352 | 2352 | large |
| LGP-200 | wbCABP-718 | 0.7494454 | 2352 | 2352 | large |
| LGP-200 | wbLGP-1058 | 0.7351284 | 2352 | 2352 | large |
| LGP-200 | wbLGP-1158 | 0.7272050 | 2352 | 2352 | large |
| LGP-200 | wbLGP-1258 | 0.7374204 | 2352 | 2352 | large |
| LGP-200 | wbLGP-1358 | 0.7088809 | 2352 | 2352 | large |
| LGP-200 | wbLGP-458 | 0.7639591 | 2352 | 2352 | large |
| LGP-200 | wbLGP-558 | 0.7610277 | 2352 | 2352 | large |
| LGP-200 | wbLGP-658 | 0.7432334 | 2352 | 2352 | large |
| LGP-200 | wbLGP-758 | 0.7366071 | 2352 | 2352 | large |
| LGP-200 | wbLGP-858 | 0.7386798 | 2352 | 2352 | large |
| LGP-200 | wbLGP-958 | 0.7282388 | 2352 | 2352 | large |
| LGP-300 | LGP-400 | 0.1569344 | 2352 | 2352 | small |
| LGP-300 | LGP-500 | 0.4014572 | 2352 | 2352 | moderate |
| LGP-300 | LGP-600 | 0.1086461 | 2352 | 2352 | small |
| LGP-300 | LGP-700 | 0.0305094 | 2352 | 2352 | small |
| LGP-300 | LGP-800 | 0.1105415 | 2352 | 2352 | small |
| LGP-300 | LGP-900 | 0.0588944 | 2352 | 2352 | small |
| LGP-300 | MMP-360 | 0.1802777 | 2352 | 2352 | small |
| LGP-300 | wbBNP-568 | 0.7393016 | 2352 | 2352 | large |
| LGP-300 | wbCABP-718 | 0.7409097 | 2352 | 2352 | large |
| LGP-300 | wbLGP-1058 | 0.7254031 | 2352 | 2352 | large |
| LGP-300 | wbLGP-1158 | 0.7146360 | 2352 | 2352 | large |
| LGP-300 | wbLGP-1258 | 0.7276375 | 2352 | 2352 | large |
| LGP-300 | wbLGP-1358 | 0.6950338 | 2352 | 2352 | large |
| LGP-300 | wbLGP-458 | 0.7545390 | 2352 | 2352 | large |

**Table S8.** Parcellation - Effect size *(continued)*

| group1 | group2 | effsize | n1 | n2 | magnitude |
|---|---|---|---|---|---|
| LGP-300 | wbLGP-558 | 0.7518696 | 2352 | 2352 | large |
| LGP-300 | wbLGP-658 | 0.7305186 | 2352 | 2352 | large |
| LGP-300 | wbLGP-758 | 0.7288353 | 2352 | 2352 | large |
| LGP-300 | wbLGP-858 | 0.7315947 | 2352 | 2352 | large |
| LGP-300 | wbLGP-958 | 0.7184744 | 2352 | 2352 | large |
| LGP-400 | LGP-500 | 0.3106815 | 2352 | 2352 | moderate |
| LGP-400 | LGP-600 | 0.0239758 | 2352 | 2352 | small |
| LGP-400 | LGP-700 | 0.2024827 | 2352 | 2352 | small |
| LGP-400 | LGP-800 | 0.0547714 | 2352 | 2352 | small |
| LGP-400 | LGP-900 | 0.1808123 | 2352 | 2352 | small |
| LGP-400 | MMP-360 | 0.2340978 | 2352 | 2352 | small |
| LGP-400 | wbBNP-568 | 0.7295883 | 2352 | 2352 | large |
| LGP-400 | wbCABP-718 | 0.7324939 | 2352 | 2352 | large |
| LGP-400 | wbLGP-1058 | 0.7127931 | 2352 | 2352 | large |
| LGP-400 | wbLGP-1158 | 0.7034513 | 2352 | 2352 | large |
| LGP-400 | wbLGP-1258 | 0.7163478 | 2352 | 2352 | large |
| LGP-400 | wbLGP-1358 | 0.6809360 | 2352 | 2352 | large |
| LGP-400 | wbLGP-458 | 0.7489080 | 2352 | 2352 | large |
| LGP-400 | wbLGP-558 | 0.7418199 | 2352 | 2352 | large |
| LGP-400 | wbLGP-658 | 0.7177687 | 2352 | 2352 | large |
| LGP-400 | wbLGP-758 | 0.7184835 | 2352 | 2352 | large |
| LGP-400 | wbLGP-858 | 0.7209195 | 2352 | 2352 | large |
| LGP-400 | wbLGP-958 | 0.7072074 | 2352 | 2352 | large |
| LGP-500 | LGP-600 | 0.3060236 | 2352 | 2352 | moderate |
| LGP-500 | LGP-700 | 0.4001344 | 2352 | 2352 | moderate |
| LGP-500 | LGP-800 | 0.3747741 | 2352 | 2352 | moderate |
| LGP-500 | LGP-900 | 0.3739231 | 2352 | 2352 | moderate |
| LGP-500 | MMP-360 | 0.3762905 | 2352 | 2352 | moderate |
| LGP-500 | wbBNP-568 | 0.7165713 | 2352 | 2352 | large |
| LGP-500 | wbCABP-718 | 0.7182115 | 2352 | 2352 | large |
| LGP-500 | wbLGP-1058 | 0.7002399 | 2352 | 2352 | large |
| LGP-500 | wbLGP-1158 | 0.6914885 | 2352 | 2352 | large |
| LGP-500 | wbLGP-1258 | 0.7026320 | 2352 | 2352 | large |
| LGP-500 | wbLGP-1358 | 0.6590763 | 2352 | 2352 | large |
| LGP-500 | wbLGP-458 | 0.7298496 | 2352 | 2352 | large |
| LGP-500 | wbLGP-558 | 0.7343705 | 2352 | 2352 | large |
| LGP-500 | wbLGP-658 | 0.7035363 | 2352 | 2352 | large |
| LGP-500 | wbLGP-758 | 0.7031792 | 2352 | 2352 | large |
| LGP-500 | wbLGP-858 | 0.7123423 | 2352 | 2352 | large |
| LGP-500 | wbLGP-958 | 0.6868862 | 2352 | 2352 | large |
| LGP-600 | LGP-700 | 0.2268428 | 2352 | 2352 | small |
| LGP-600 | LGP-800 | 0.0462445 | 2352 | 2352 | small |
| LGP-600 | LGP-900 | 0.1816469 | 2352 | 2352 | small |
| LGP-600 | MMP-360 | 0.1886473 | 2352 | 2352 | small |
| LGP-600 | wbBNP-568 | 0.7481082 | 2352 | 2352 | large |
| LGP-600 | wbCABP-718 | 0.7481815 | 2352 | 2352 | large |

**Table S8.** Parcellation - Effect size *(continued)*

| group1 | group2 | effsize | n1 | n2 | magnitude |
|---|---|---|---|---|---|
| LGP-600 | wbLGP-1058 | 0.7383695 | 2352 | 2352 | large |
| LGP-600 | wbLGP-1158 | 0.7294442 | 2352 | 2352 | large |
| LGP-600 | wbLGP-1258 | 0.7369489 | 2352 | 2352 | large |
| LGP-600 | wbLGP-1358 | 0.7042384 | 2352 | 2352 | large |
| LGP-600 | wbLGP-458 | 0.7557913 | 2352 | 2352 | large |
| LGP-600 | wbLGP-558 | 0.7562789 | 2352 | 2352 | large |
| LGP-600 | wbLGP-658 | 0.7398462 | 2352 | 2352 | large |
| LGP-600 | wbLGP-758 | 0.7355434 | 2352 | 2352 | large |
| LGP-600 | wbLGP-858 | 0.7421715 | 2352 | 2352 | large |
| LGP-600 | wbLGP-958 | 0.7452103 | 2352 | 2352 | large |
| LGP-700 | LGP-800 | 0.1482687 | 2352 | 2352 | small |
| LGP-700 | LGP-900 | 0.0223819 | 2352 | 2352 | small |
| LGP-700 | MMP-360 | 0.1459144 | 2352 | 2352 | small |
| LGP-700 | wbBNP-568 | 0.7385190 | 2352 | 2352 | large |
| LGP-700 | wbCABP-718 | 0.7428081 | 2352 | 2352 | large |
| LGP-700 | wbLGP-1058 | 0.7280828 | 2352 | 2352 | large |
| LGP-700 | wbLGP-1158 | 0.7198417 | 2352 | 2352 | large |
| LGP-700 | wbLGP-1258 | 0.7322622 | 2352 | 2352 | large |
| LGP-700 | wbLGP-1358 | 0.6979349 | 2352 | 2352 | large |
| LGP-700 | wbLGP-458 | 0.7555548 | 2352 | 2352 | large |
| LGP-700 | wbLGP-558 | 0.7506720 | 2352 | 2352 | large |
| LGP-700 | wbLGP-658 | 0.7296047 | 2352 | 2352 | large |
| LGP-700 | wbLGP-758 | 0.7284709 | 2352 | 2352 | large |
| LGP-700 | wbLGP-858 | 0.7297612 | 2352 | 2352 | large |
| LGP-700 | wbLGP-958 | 0.7241542 | 2352 | 2352 | large |
| LGP-800 | LGP-900 | 0.1523587 | 2352 | 2352 | small |
| LGP-800 | MMP-360 | 0.2294773 | 2352 | 2352 | small |
| LGP-800 | wbBNP-568 | 0.7247815 | 2352 | 2352 | large |
| LGP-800 | wbCABP-718 | 0.7278916 | 2352 | 2352 | large |
| LGP-800 | wbLGP-1058 | 0.7110976 | 2352 | 2352 | large |
| LGP-800 | wbLGP-1158 | 0.7148132 | 2352 | 2352 | large |
| LGP-800 | wbLGP-1258 | 0.7177082 | 2352 | 2352 | large |
| LGP-800 | wbLGP-1358 | 0.6815238 | 2352 | 2352 | large |
| LGP-800 | wbLGP-458 | 0.7424878 | 2352 | 2352 | large |
| LGP-800 | wbLGP-558 | 0.7403174 | 2352 | 2352 | large |
| LGP-800 | wbLGP-658 | 0.7160305 | 2352 | 2352 | large |
| LGP-800 | wbLGP-758 | 0.7131063 | 2352 | 2352 | large |
| LGP-800 | wbLGP-858 | 0.7184349 | 2352 | 2352 | large |
| LGP-800 | wbLGP-958 | 0.7032708 | 2352 | 2352 | large |
| LGP-900 | MMP-360 | 0.1404051 | 2352 | 2352 | small |
| LGP-900 | wbBNP-568 | 0.7387104 | 2352 | 2352 | large |
| LGP-900 | wbCABP-718 | 0.7433471 | 2352 | 2352 | large |
| LGP-900 | wbLGP-1058 | 0.7243857 | 2352 | 2352 | large |
| LGP-900 | wbLGP-1158 | 0.7255897 | 2352 | 2352 | large |
| LGP-900 | wbLGP-1258 | 0.7343608 | 2352 | 2352 | large |
| LGP-900 | wbLGP-1358 | 0.7026858 | 2352 | 2352 | large |
| LGP-900 | wbLGP-458 | 0.7545988 | 2352 | 2352 | large |

**Table S8.** Parcellation - Effect size *(continued)*

| group1 | group2 | effsize | n1 | n2 | magnitude |
|---|---|---|---|---|---|
| LGP-900 | wbLGP-558 | 0.7503933 | 2352 | 2352 | large |
| LGP-900 | wbLGP-658 | 0.7302955 | 2352 | 2352 | large |
| LGP-900 | wbLGP-758 | 0.7241196 | 2352 | 2352 | large |
| LGP-900 | wbLGP-858 | 0.7301608 | 2352 | 2352 | large |
| LGP-900 | wbLGP-958 | 0.7201412 | 2352 | 2352 | large |
| MMP-360 | wbBNP-568 | 0.7444282 | 2352 | 2352 | large |
| MMP-360 | wbCABP-718 | 0.7535082 | 2352 | 2352 | large |
| MMP-360 | wbLGP-1058 | 0.7311101 | 2352 | 2352 | large |
| MMP-360 | wbLGP-1158 | 0.7201675 | 2352 | 2352 | large |
| MMP-360 | wbLGP-1258 | 0.7319867 | 2352 | 2352 | large |
| MMP-360 | wbLGP-1358 | 0.6975871 | 2352 | 2352 | large |
| MMP-360 | wbLGP-458 | 0.7584484 | 2352 | 2352 | large |
| MMP-360 | wbLGP-558 | 0.7548697 | 2352 | 2352 | large |
| MMP-360 | wbLGP-658 | 0.7382470 | 2352 | 2352 | large |
| MMP-360 | wbLGP-758 | 0.7320983 | 2352 | 2352 | large |
| MMP-360 | wbLGP-858 | 0.7345617 | 2352 | 2352 | large |
| MMP-360 | wbLGP-958 | 0.7174438 | 2352 | 2352 | large |
| wbBNP-568 | wbCABP-718 | 0.2728574 | 2352 | 2352 | small |
| wbBNP-568 | wbLGP-1058 | 0.2162613 | 2352 | 2352 | small |
| wbBNP-568 | wbLGP-1158 | 0.1995484 | 2352 | 2352 | small |
| wbBNP-568 | wbLGP-1258 | 0.1386707 | 2352 | 2352 | small |
| wbBNP-568 | wbLGP-1358 | 0.2717290 | 2352 | 2352 | small |
| wbBNP-568 | wbLGP-458 | 0.0877119 | 2352 | 2352 | small |
| wbBNP-568 | wbLGP-558 | 0.0947171 | 2352 | 2352 | small |
| wbBNP-568 | wbLGP-658 | 0.2160494 | 2352 | 2352 | small |
| wbBNP-568 | wbLGP-758 | 0.2540569 | 2352 | 2352 | small |
| wbBNP-568 | wbLGP-858 | 0.1592633 | 2352 | 2352 | small |
| wbBNP-568 | wbLGP-958 | 0.2937065 | 2352 | 2352 | small |
| wbCABP-718 | wbLGP-1058 | 0.0359091 | 2352 | 2352 | small |
| wbCABP-718 | wbLGP-1158 | 0.0233259 | 2352 | 2352 | small |
| wbCABP-718 | wbLGP-1258 | 0.0388589 | 2352 | 2352 | small |
| wbCABP-718 | wbLGP-1358 | 0.1106258 | 2352 | 2352 | small |
| wbCABP-718 | wbLGP-458 | 0.2848362 | 2352 | 2352 | small |
| wbCABP-718 | wbLGP-558 | 0.3284945 | 2352 | 2352 | moderate |
| wbCABP-718 | wbLGP-658 | 0.1230697 | 2352 | 2352 | small |
| wbCABP-718 | wbLGP-758 | 0.0370190 | 2352 | 2352 | small |
| wbCABP-718 | wbLGP-858 | 0.0991309 | 2352 | 2352 | small |
| wbCABP-718 | wbLGP-958 | 0.0362152 | 2352 | 2352 | small |
| wbLGP-1058 | wbLGP-1158 | 0.0561301 | 2352 | 2352 | small |
| wbLGP-1058 | wbLGP-1258 | 0.1116554 | 2352 | 2352 | small |
| wbLGP-1058 | wbLGP-1358 | 0.1822538 | 2352 | 2352 | small |
| wbLGP-1058 | wbLGP-458 | 0.2180510 | 2352 | 2352 | small |
| wbLGP-1058 | wbLGP-558 | 0.2778507 | 2352 | 2352 | small |
| wbLGP-1058 | wbLGP-658 | 0.1051919 | 2352 | 2352 | small |
| wbLGP-1058 | wbLGP-758 | 0.0228152 | 2352 | 2352 | small |
| wbLGP-1058 | wbLGP-858 | 0.1661968 | 2352 | 2352 | small |
| wbLGP-1058 | wbLGP-958 | 0.0818575 | 2352 | 2352 | small |

**Table S8.** Parcellation - Effect size *(continued)*

| group1 | group2 | effsize | n1 | n2 | magnitude |
|---|---|---|---|---|---|
| wbLGP-1158 | wbLGP-1258 | 0.1252645 | 2352 | 2352 | small |
| wbLGP-1158 | wbLGP-1358 | 0.1480026 | 2352 | 2352 | small |
| wbLGP-1158 | wbLGP-458 | 0.2274392 | 2352 | 2352 | small |
| wbLGP-1158 | wbLGP-558 | 0.2439547 | 2352 | 2352 | small |
| wbLGP-1158 | wbLGP-658 | 0.1045024 | 2352 | 2352 | small |
| wbLGP-1158 | wbLGP-758 | 0.0322159 | 2352 | 2352 | small |
| wbLGP-1158 | wbLGP-858 | 0.1551948 | 2352 | 2352 | small |
| wbLGP-1158 | wbLGP-958 | 0.0414800 | 2352 | 2352 | small |
| wbLGP-1258 | wbLGP-1358 | 0.2590570 | 2352 | 2352 | small |
| wbLGP-1258 | wbLGP-458 | 0.1735670 | 2352 | 2352 | small |
| wbLGP-1258 | wbLGP-558 | 0.1977710 | 2352 | 2352 | small |
| wbLGP-1258 | wbLGP-658 | 0.0419666 | 2352 | 2352 | small |
| wbLGP-1258 | wbLGP-758 | 0.0287042 | 2352 | 2352 | small |
| wbLGP-1258 | wbLGP-858 | 0.0559503 | 2352 | 2352 | small |
| wbLGP-1258 | wbLGP-958 | 0.1063742 | 2352 | 2352 | small |
| wbLGP-1358 | wbLGP-458 | 0.2918365 | 2352 | 2352 | small |
| wbLGP-1358 | wbLGP-558 | 0.3015406 | 2352 | 2352 | moderate |
| wbLGP-1358 | wbLGP-658 | 0.1847120 | 2352 | 2352 | small |
| wbLGP-1358 | wbLGP-758 | 0.1378691 | 2352 | 2352 | small |
| wbLGP-1358 | wbLGP-858 | 0.2044047 | 2352 | 2352 | small |
| wbLGP-1358 | wbLGP-958 | 0.0714880 | 2352 | 2352 | small |
| wbLGP-458 | wbLGP-558 | 0.0115092 | 2352 | 2352 | small |
| wbLGP-458 | wbLGP-658 | 0.2420159 | 2352 | 2352 | small |
| wbLGP-458 | wbLGP-758 | 0.2603953 | 2352 | 2352 | small |
| wbLGP-458 | wbLGP-858 | 0.1864424 | 2352 | 2352 | small |
| wbLGP-458 | wbLGP-958 | 0.2951134 | 2352 | 2352 | small |
| wbLGP-558 | wbLGP-658 | 0.3159288 | 2352 | 2352 | moderate |
| wbLGP-558 | wbLGP-758 | 0.3114833 | 2352 | 2352 | moderate |
| wbLGP-558 | wbLGP-858 | 0.2210771 | 2352 | 2352 | small |
| wbLGP-558 | wbLGP-958 | 0.3280744 | 2352 | 2352 | moderate |
| wbLGP-658 | wbLGP-758 | 0.1151963 | 2352 | 2352 | small |
| wbLGP-658 | wbLGP-858 | 0.0129152 | 2352 | 2352 | small |
| wbLGP-658 | wbLGP-958 | 0.1975321 | 2352 | 2352 | small |
| wbLGP-758 | wbLGP-858 | 0.1643324 | 2352 | 2352 | small |
| wbLGP-758 | wbLGP-958 | 0.0960455 | 2352 | 2352 | small |
| wbLGP-858 | wbLGP-958 | 0.1928274 | 2352 | 2352 | small |

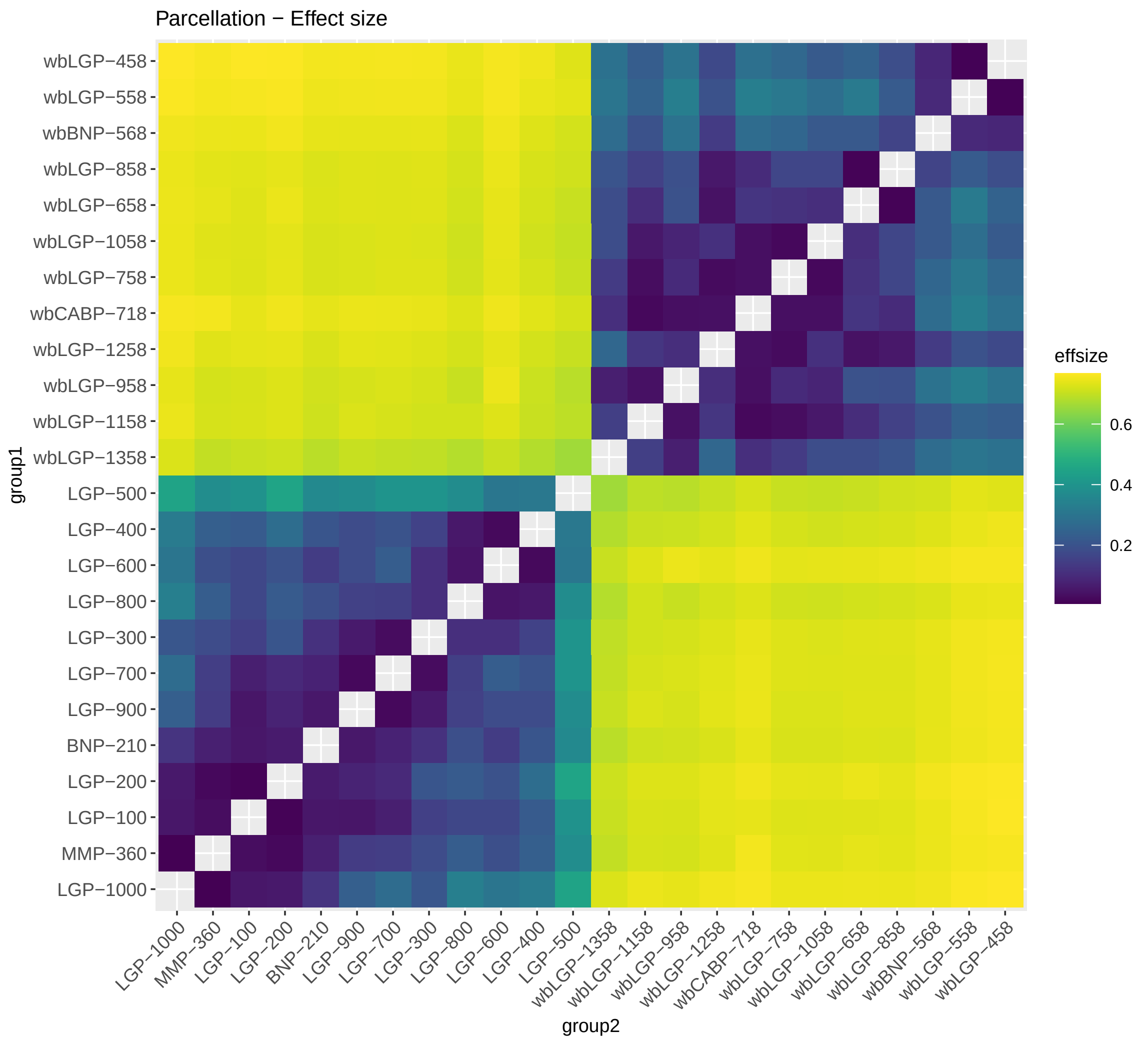

**Fig. S6.** Parcellation - Effect size

#### 1.12 Variability Changes

Whole Brain vs. Cortex

- Quadrant 1: $V_w \uparrow, V_b \uparrow$: 0.0093684
- Quadrant 2: $V_w \downarrow, V_b \uparrow$: 0.5999534
- Quadrant 3: $V_w \downarrow, V_b \downarrow$: 0.3906782
- Quadrant 4: $V_w \uparrow, V_b \downarrow$: 0

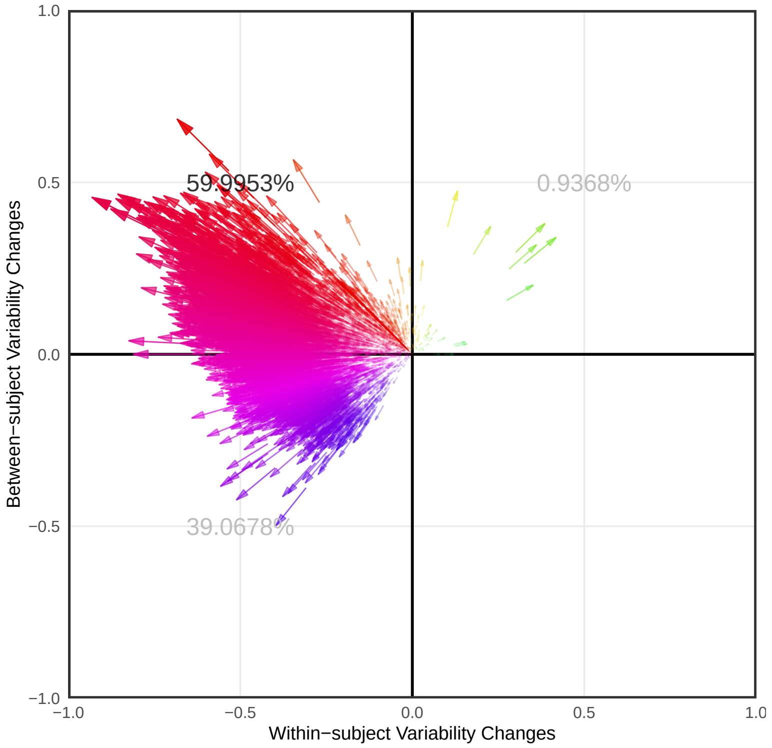

**Fig. S7.** Parcellation - Variability Changes

### 2 Frequency Bands - Edge Construction

#### 2.1 ICC Density distribution

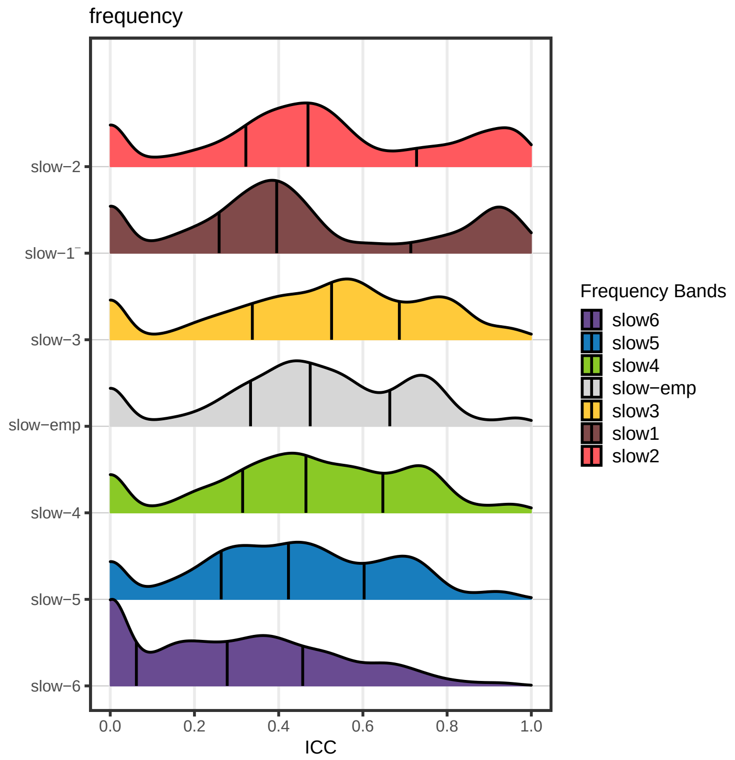

**Fig. S8.** Frequency - ICC Density distribution

#### 2.2 Almost Perfect ( ICCs > 0.8 )

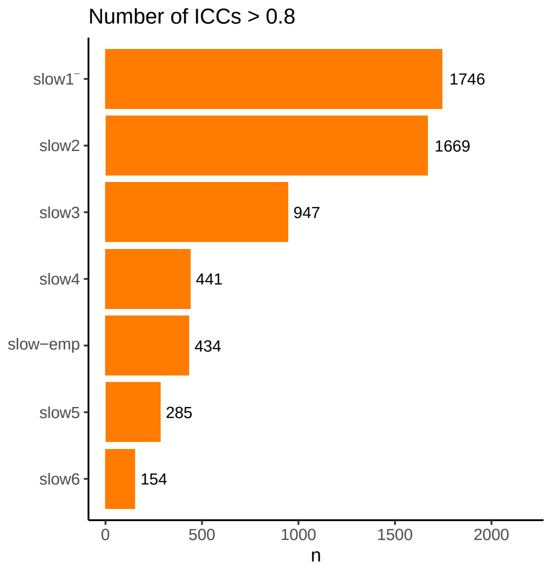

**Fig. S9.** Frequency - Number of ICC >0.8

#### 2.3 Substantial or Above ( ICCs > 0.6 )

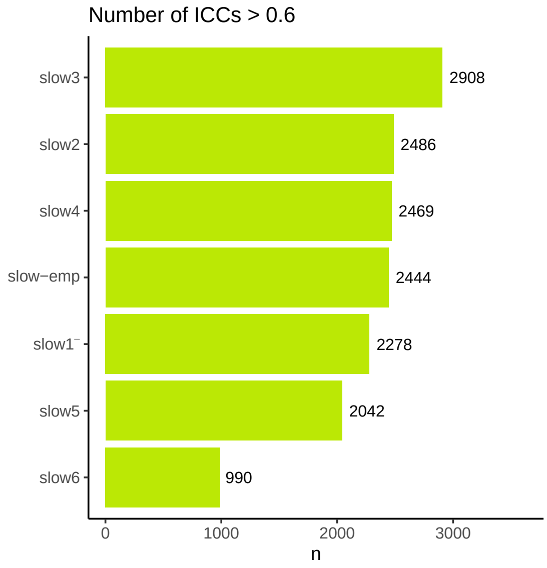

**Fig. S10.** Frequency - Number of ICC >0.6

#### 2.4 Variability Changes

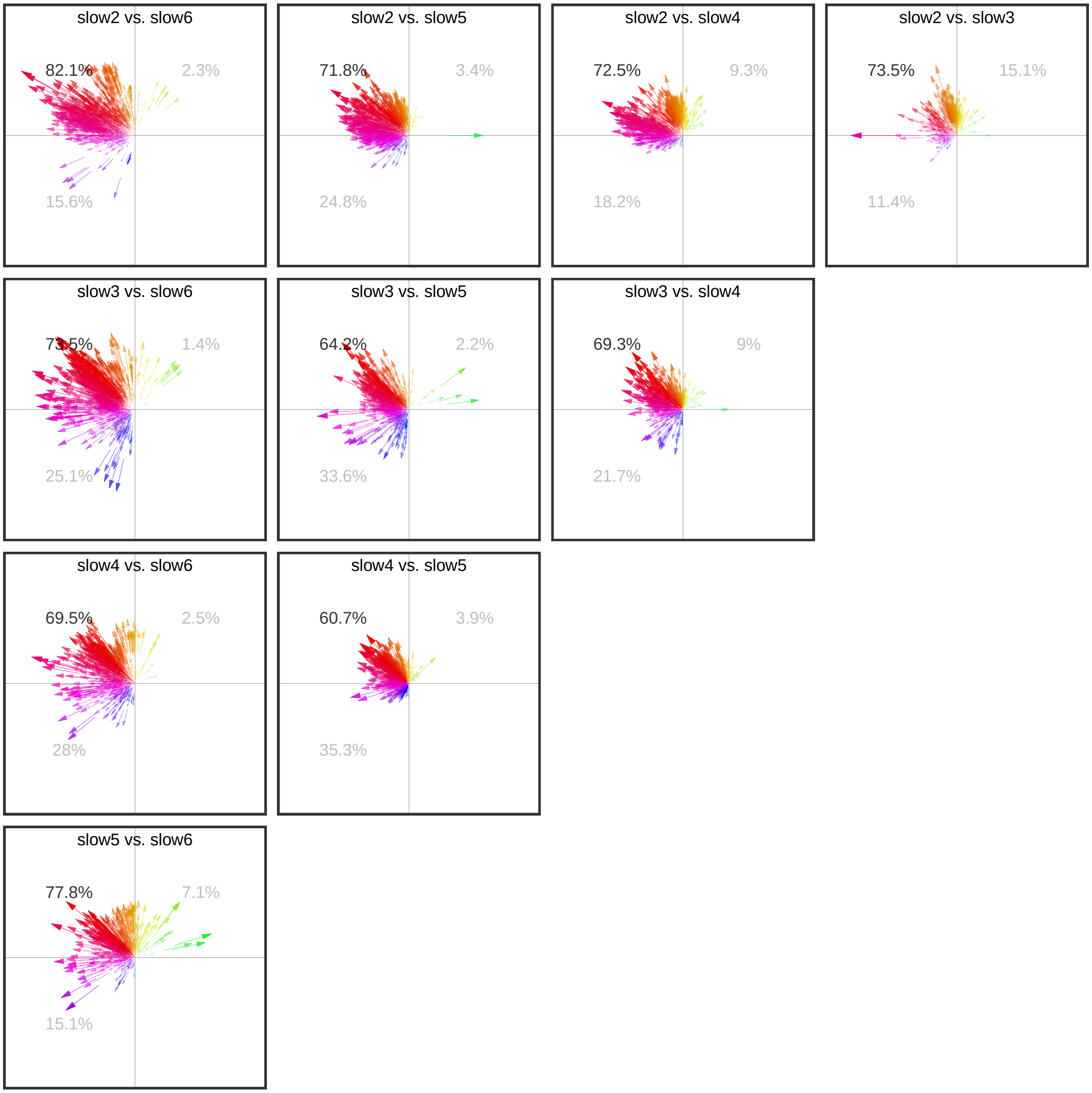

**Fig. S11.** Frequency - Variability Changes

#### 2.5 Descriptive statistics Mean

**Table S9.** Frequency - ICC Mean

| freq | variable | n | mean | mean_z2r |
|---|---|---|---|---|
| slow2 | ICC.z | 8064 | 0.689 | 0.5973392 |
| slow1 | ICC.z | 8064 | 0.638 | 0.5635362 |
| slow3 | ICC.z | 8064 | 0.621 | 0.5518239 |
| slow-emp | ICC.z | 8064 | 0.575 | 0.5190218 |
| slow4 | ICC.z | 8064 | 0.560 | 0.5079774 |
| slow5 | ICC.z | 8064 | 0.494 | 0.4573854 |
| slow6 | ICC.z | 8064 | 0.331 | 0.3194190 |

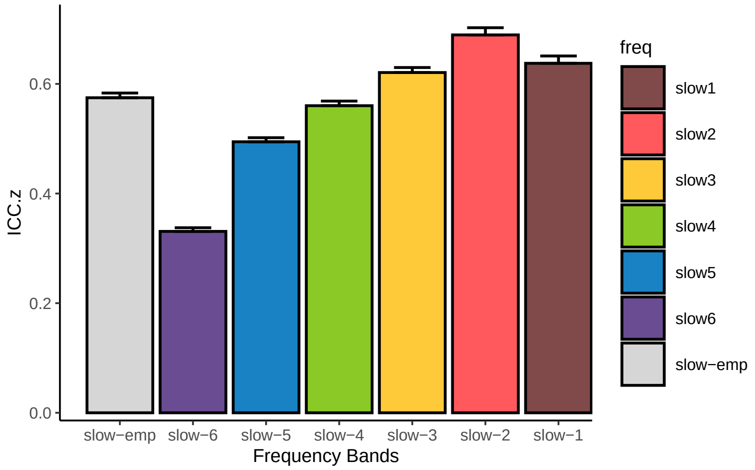

**Fig. S12.** Frequency - ICC Mean Bar plot

#### 2.6 Descriptive statistics Median

**Table S10.** Frequency - ICC Median

| freq | variable | n | median |
|---|---|---|---|
| slow3 | ICC | 8064 | 0.526 |
| slow-emp | ICC | 8064 | 0.475 |
| slow2 | ICC | 8064 | 0.470 |
| slow4 | ICC | 8064 | 0.465 |
| slow5 | ICC | 8064 | 0.423 |
| slow1 | ICC | 8064 | 0.395 |
| slow6 | ICC | 8064 | 0.278 |

#### 2.7 Friedman Test

**Table S11.** Frequency - Friedman Test

| .y. | n | statistic | df | p | method |
|---|---|---|---|---|---|
| ICC.z | 8064 | 9283.536 | 6 | 0 | Friedman test |

#### 2.8 Friedman Test Effect size

**Table S12.** Frequency - Friedman Test Effect size

| .y. | n | effsize | method | magnitude |
|---|---|---|---|---|
| ICC.z | 8064 | 0.191872 | Kendall W | small |

#### 2.9 Paired Wilcoxon signed rank test

**Table S13.** Frequency - ICC group1 vs. group2

| group1 | group2 | n1 | estimate | statistic | alternative | p | p.adj | p.adj.signif |
|---|---|---|---|---|---|---|---|---|
| slow-emp | slow1 | 8064 | -0.0162227 | 14054394 | two.sided | 9.68e-04 | 0.0203280 | * |
| slow-emp | slow2 | 8064 | -0.0864131 | 10505882 | two.sided | 0.00e+00 | 0.0000000 | **** |
| slow-emp | slow3 | 8064 | -0.0539695 | 10625450 | two.sided | 0.00e+00 | 0.0000000 | **** |
| slow-emp | slow4 | 8064 | 0.0150546 | 17102898 | two.sided | 0.00e+00 | 0.0000000 | **** |
| slow-emp | slow5 | 8064 | 0.0841908 | 23531772 | two.sided | 0.00e+00 | 0.0000000 | **** |
| slow-emp | slow6 | 8064 | 0.2231625 | 26254891 | two.sided | 0.00e+00 | 0.0000000 | **** |
| slow1 | slow2 | 8064 | -0.0624239 | 9900068 | two.sided | 0.00e+00 | 0.0000000 | **** |
| slow1 | slow3 | 8064 | -0.0200081 | 13838087 | two.sided | 8.77e-05 | 0.0018417 | ** |
| slow1 | slow4 | 8064 | 0.0325830 | 15946425 | two.sided | 0.00e+00 | 0.0000000 | **** |
| slow1 | slow5 | 8064 | 0.1039074 | 18945816 | two.sided | 0.00e+00 | 0.0000000 | **** |
| slow1 | slow6 | 8064 | 0.2451552 | 23735075 | two.sided | 0.00e+00 | 0.0000000 | **** |
| slow2 | slow3 | 8064 | 0.0367993 | 16494499 | two.sided | 0.00e+00 | 0.0000000 | **** |
| slow2 | slow4 | 8064 | 0.1038401 | 19454026 | two.sided | 0.00e+00 | 0.0000000 | **** |
| slow2 | slow5 | 8064 | 0.1643515 | 22530367 | two.sided | 0.00e+00 | 0.0000000 | **** |
| slow2 | slow6 | 8064 | 0.2906672 | 25875758 | two.sided | 0.00e+00 | 0.0000000 | **** |
| slow3 | slow4 | 8064 | 0.0703217 | 20070190 | two.sided | 0.00e+00 | 0.0000000 | **** |
| slow3 | slow5 | 8064 | 0.1355815 | 23423580 | two.sided | 0.00e+00 | 0.0000000 | **** |
| slow3 | slow6 | 8064 | 0.2698008 | 26652991 | two.sided | 0.00e+00 | 0.0000000 | **** |
| slow4 | slow5 | 8064 | 0.0707338 | 21276606 | two.sided | 0.00e+00 | 0.0000000 | **** |
| slow4 | slow6 | 8064 | 0.2083264 | 25815007 | two.sided | 0.00e+00 | 0.0000000 | **** |
| slow5 | slow6 | 8064 | 0.1410985 | 24873960 | two.sided | 0.00e+00 | 0.0000000 | **** |

#### 2.10 Kullback-leibler divergence Map

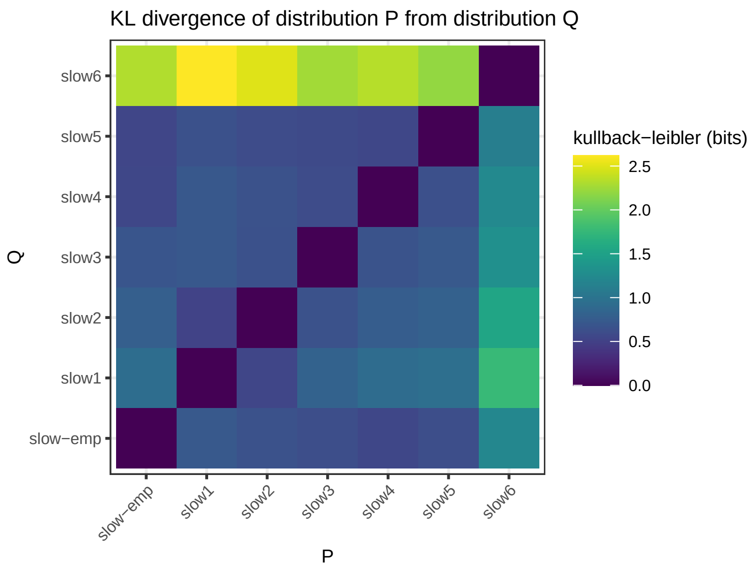

**Fig. S13.** Frequency - KL divergence Map

#### 2.11 Significance Map

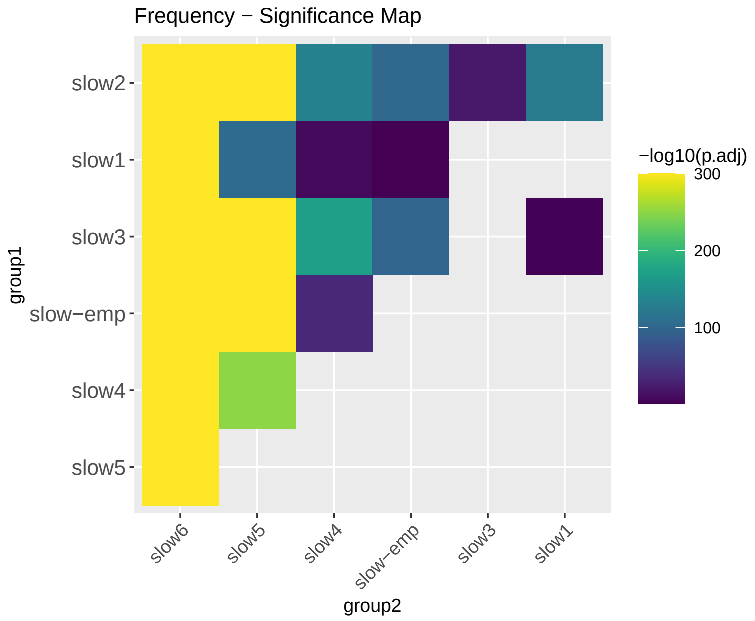

**Fig. S14.** Frequency - Significance Map

#### 2.12 Effect size

**Table S14.** Frequency - Effect size

| group1 | group2 | effsize | n1 | n2 | magnitude |
|---|---|---|---|---|---|
| slow-emp | slow1 | 0.0296722 | 8064 | 8064 | small |
| slow-emp | slow2 | 0.2374189 | 8064 | 8064 | small |
| slow-emp | slow3 | 0.2357191 | 8064 | 8064 | small |
| slow-emp | slow4 | 0.1417849 | 8064 | 8064 | small |
| slow-emp | slow5 | 0.5094890 | 8064 | 8064 | large |
| slow-emp | slow6 | 0.6519342 | 8064 | 8064 | large |
| slow1 | slow2 | 0.2699083 | 8064 | 8064 | small |
| slow1 | slow3 | 0.0520287 | 8064 | 8064 | small |
| slow1 | slow4 | 0.0646878 | 8064 | 8064 | small |
| slow1 | slow5 | 0.2408327 | 8064 | 8064 | small |
| slow1 | slow6 | 0.5023808 | 8064 | 8064 | large |
| slow2 | slow3 | 0.1000050 | 8064 | 8064 | small |
| slow2 | slow4 | 0.2699834 | 8064 | 8064 | small |
| slow2 | slow5 | 0.4490800 | 8064 | 8064 | moderate |
| slow2 | slow6 | 0.6292308 | 8064 | 8064 | large |
| slow3 | slow4 | 0.3096116 | 8064 | 8064 | moderate |
| slow3 | slow5 | 0.5028784 | 8064 | 8064 | large |
| slow3 | slow6 | 0.6802808 | 8064 | 8064 | large |
| slow4 | slow5 | 0.3770518 | 8064 | 8064 | moderate |
| slow4 | slow6 | 0.6310338 | 8064 | 8064 | large |
| slow5 | slow6 | 0.5700624 | 8064 | 8064 | large |

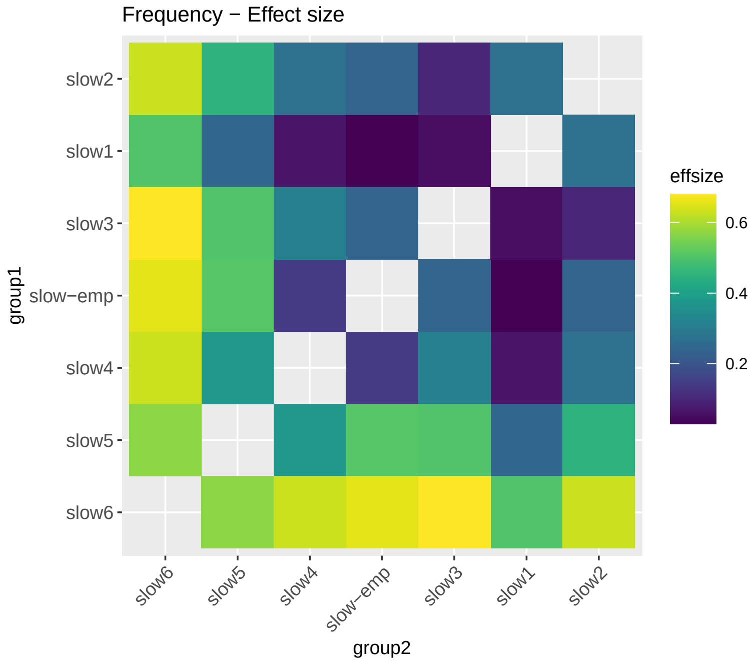

**Fig. S15.** Frequency - Effect size

### 3 R Tranforms - Edge Construction

Many network metrics are not well defined for negatively weighted connections. In order to ensure that the connection weights are positive only, we applied four types of transformations to the symmetric correlation matrix: the **positive** (Eq.pos), **absolute** (Eq.abs), **exponential** (Eq.exp) and **distance-inverse** (Eq.div) functions, respectively. This avoids the negative values in the inter-node connectivity matrix $W = (w_{ij})$ where $z_{ij} = \tanh^{-1}(r_{ij})$ is Fisher's $z$—transformation.

$$w_{ij} = \frac{z_{ij} + |z_{ij}|}{2} \in [0, \infty) \tag{pos}$$

$$w_{ij} = |z_{ij}| \in [0, \infty) \tag{abs}$$

$$w_{ij} = \mathbf{e}^{z_{ij}} \in [0, \infty) \tag{exp}$$

$$w_{ij} = \frac{2}{\sqrt{2 \times (1 - r_{ij})}} \in (0, \infty) \tag{div}$$

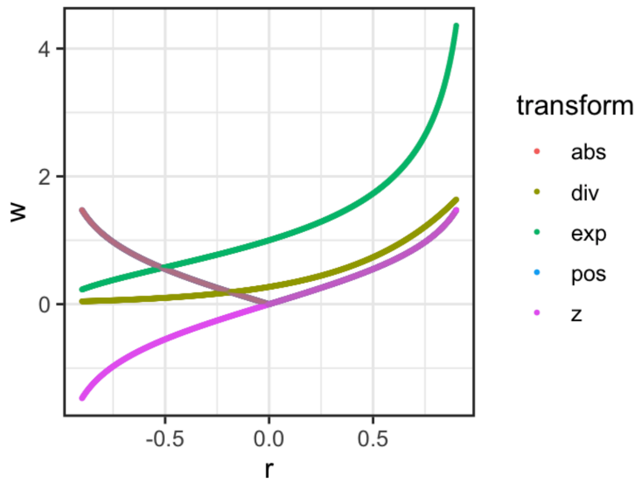

**Fig. S16.** Transform r to weight

#### 3.1 ICC Density distribution

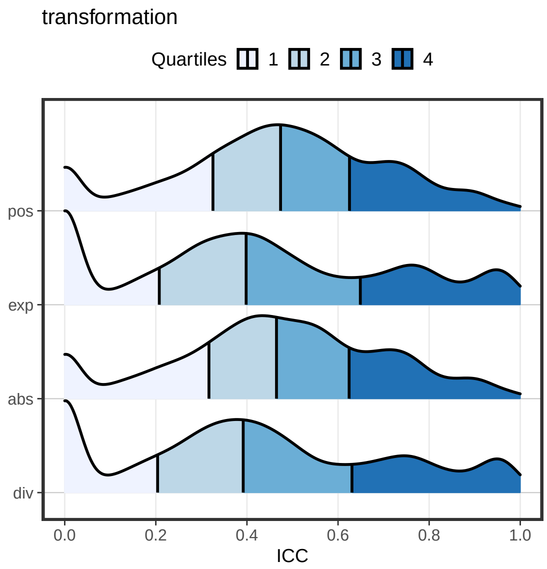

**Fig. S17.** Transforms - ICC Density distribution

#### 3.2 Almost Perfect ( ICCs > 0.8 )

**Table S15.** Transforms - Number of ICCs > 0.8

| transform | n |
|---|---|
| exp | 1855 |
| div | 1740 |
| abs | 1050 |
| pos | 1031 |

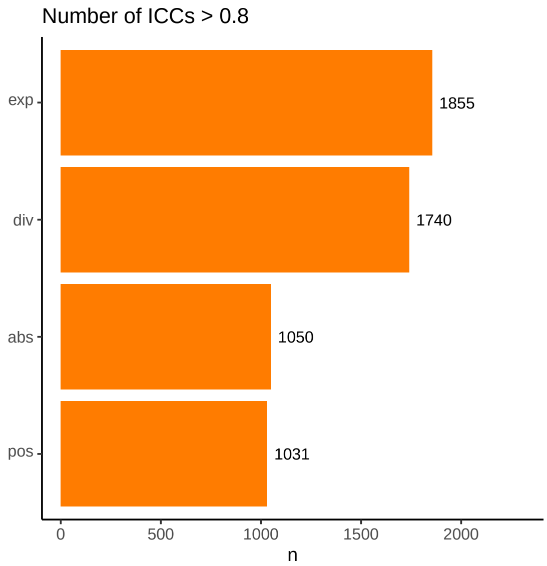

**Fig. S18.** Transforms - Number of ICC > 0.8

#### 3.3 Substantial or Above ( ICCs > 0.6 )

**Table S16.** Transforms - Number of ICCs > 0.6

| transform | n |
|---|---|
| pos | 3977 |
| exp | 3930 |
| abs | 3912 |
| div | 3798 |

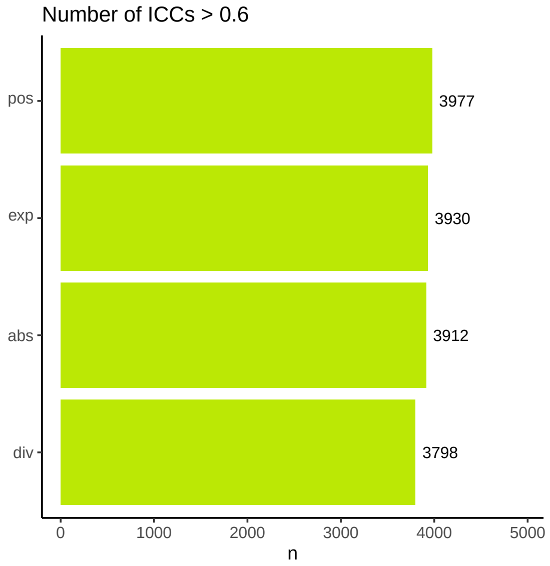

**Fig. S19.** Transforms - Number of ICC > 0.8

#### 3.4 Descriptive statistics Mean

**Table S17.** Transforms - ICC Mean

| transform | variable | n | mean | mean_z2r |
|---|---|---|---|---|
| pos | ICC.z | 14112 | 0.564 | 0.5109392 |
| exp | ICC.z | 14112 | 0.561 | 0.5087190 |
| abs | ICC.z | 14112 | 0.560 | 0.5079774 |
| div | ICC.z | 14112 | 0.548 | 0.4990198 |

#### 3.5 Descriptive statistics Median

**Table S18.** Transforms - ICC Median

| transform | variable | n | median |
|---|---|---|---|
| pos | ICC | 14112 | 0.474 |
| abs | ICC | 14112 | 0.465 |
| exp | ICC | 14112 | 0.398 |
| div | ICC | 14112 | 0.392 |

#### 3.6 Friedman Test

**Table S19.** Transforms - Friedman Test

| .y. | n | statistic | df | p | method |
|---|---|---|---|---|---|
| ICC.z | 14112 | 480.0639 | 3 | 0 | Friedman test |

#### 3.7 Friedman Test Effect size

**Table S20.** Transforms - Friedman Test Effect size

| .y. | n | effsize | method | magnitude |
|---|---|---|---|---|
| ICC.z | 14112 | 0.0113394 | Kendall W | small |

#### 3.8 Paired Wilcoxon signed rank test

**Table S21.** Transforms - ICC group1 vs. group2

| group1 | group2 | n1 | statistic | alternative | p | p.adj | p.adj.signif |
|---|---|---|---|---|---|---|---|
| abs | div | 14112 | 50949564 | two.sided | 0e+00 | 0.0e+00 | **** |
| abs | exp | 14112 | 47946915 | two.sided | 8e-07 | 4.8e-06 | **** |
| abs | pos | 14112 | 44430310 | two.sided | 3e-03 | 1.8e-02 | * |
| div | exp | 14112 | 35746898 | two.sided | 0e+00 | 0.0e+00 | **** |
| div | pos | 14112 | 39669204 | two.sided | 0e+00 | 0.0e+00 | **** |
| exp | pos | 14112 | 42638630 | two.sided | 0e+00 | 0.0e+00 | **** |

##### 3.9 Kullback-leibler divergence Map

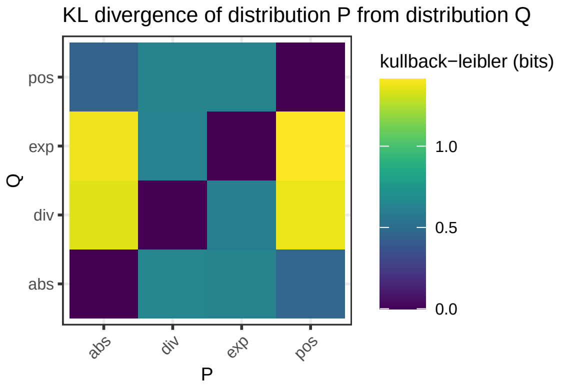

**Fig. S20.** Transforms - KL divergence Map

##### 3.10 Significance map

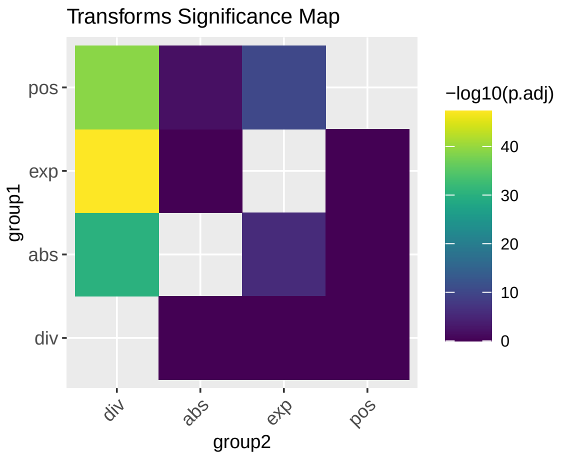

**Fig. S21.** Transforms - Significance Map

##### 3.11 Effect size

**Table S22.** Transforms - Effect size

| group1 | group2 | effsize | n1 | n2 | magnitude |
|---|---|---|---|---|---|
| abs | div | 0.1009229 | 14112 | 14112 | small |
| abs | exp | 0.0439603 | 14112 | 14112 | small |
| abs | pos | 0.0232195 | 14112 | 14112 | small |
| div | exp | 0.1257376 | 14112 | 14112 | small |
| div | pos | 0.1139965 | 14112 | 14112 | small |
| exp | pos | 0.0582916 | 14112 | 14112 | small |

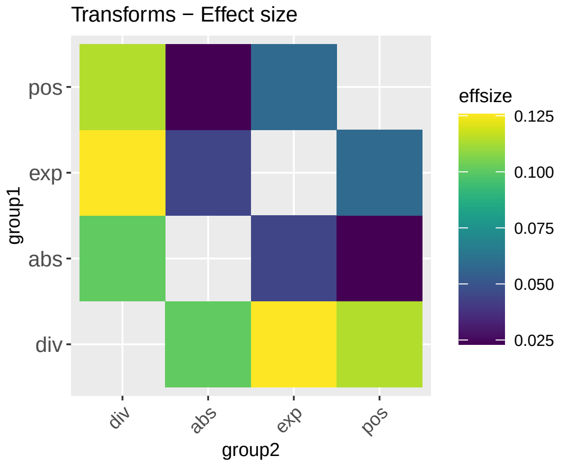

**Fig. S22.** Transforms - Effect size

### 4 Schemes - Edge Construction

#### 4.1 ICC Density distribution

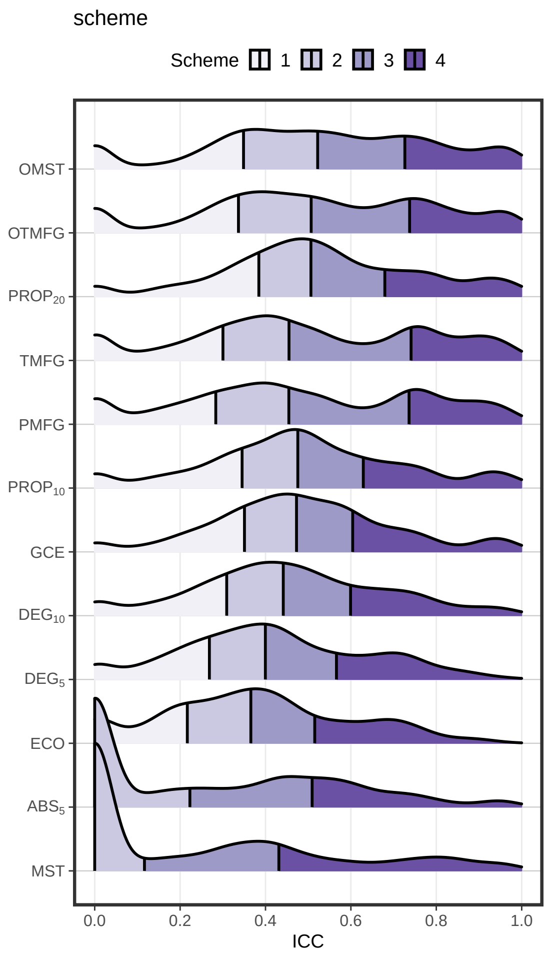

**Fig. S23.** Schemes - ICC Density distribution

#### 4.2 Almost Perfect ( ICCs > 0.8 )

**Table S23.** Schemes - Number of ICCs > 0.8

| scheme | n |
|---|---|
| OTMFG | 781 |
| TMFG | 767 |
| OMST | 765 |
| PMFG | 737 |
| $PROP_{20}$ | 632 |
| $PROP_{10}$ | 445 |
| MST | 362 |
| GCE | 352 |
| $DEG_{15}$ | 306 |
| $DEG_{5}$ | 213 |
| $ABS_{05}$ | 189 |
| ECO | 127 |

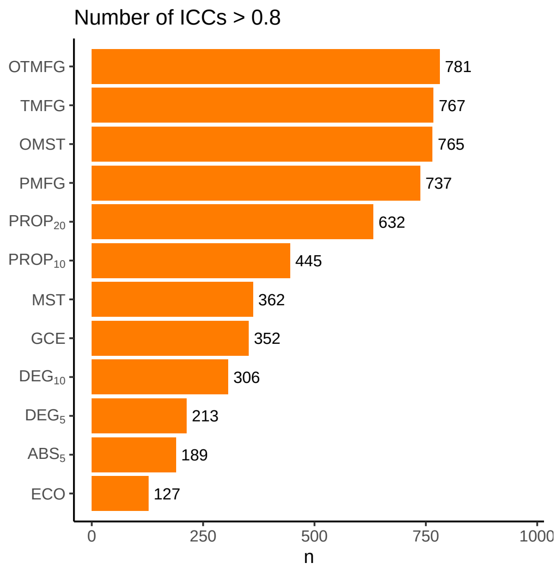

**Fig. S24.** Schemes - Number of ICC > 0.8

#### 4.3 Substantial or Above ( ICCs > 0.6 )

**Table S24.** Schemes - Number of ICCs > 0.8

| scheme | n |
|---|---|
| OMST | 1860 |
| OTMFG | 1832 |
| TMFG | 1670 |
| PMFG | 1659 |
| $PROP_{20}$ | 1564 |
| $PROP_{10}$ | 1345 |
| GCE | 1210 |
| $DEG_{15}$ | 1175 |
| $DEG_{5}$ | 1035 |
| ECO | 837 |
| MST | 719 |
| $ABS_{05}$ | 711 |

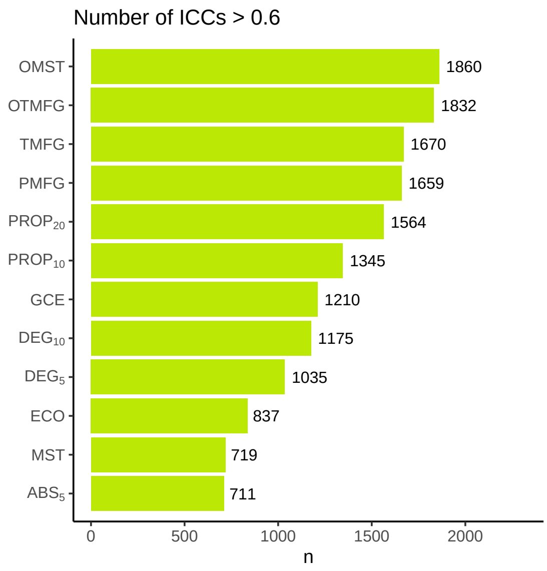

**Fig. S25.** Schemes - Number of ICC > 0.6

#### 4.4 Varibility Changes

OMST, OTMFG, $PROP_{20}$, TMFG, PMFG, $PROP_{10}$

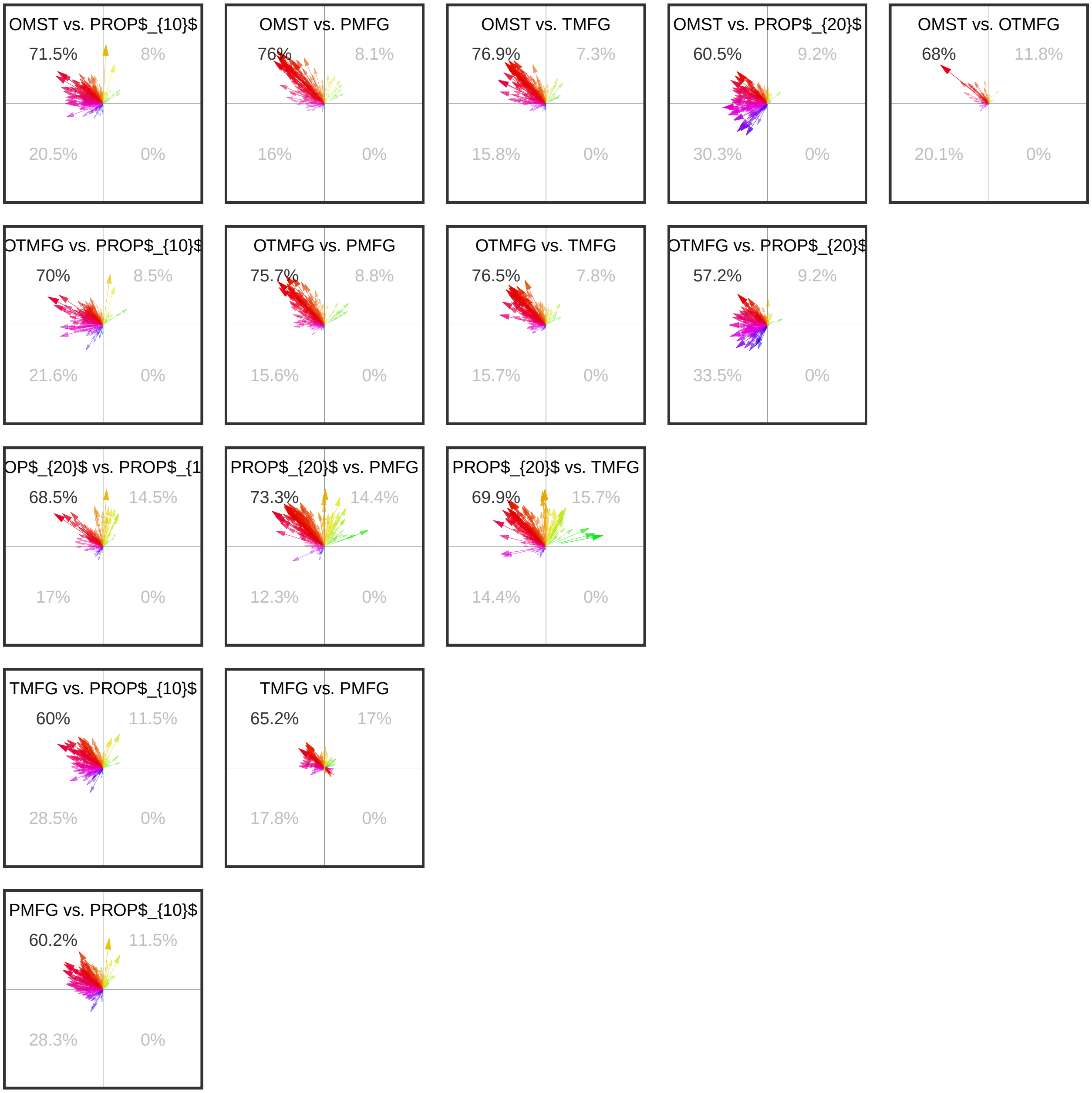

**Fig. S26.** Schemes - Variability Changes

#### 4.5 Descriptive statistics Mean

**Table S25.** Schemes - ICC Mean

| scheme | variable | n | mean | meanz2r |
|---|---|---|---|---|
| OMST | ICC.z | 4704 | 0.706 | 0.6081624 |
| OTMFG | ICC.z | 4704 | 0.697 | 0.6024601 |
| $\text{PROP}_{20}$ | ICC.z | 4704 | 0.683 | 0.5934662 |
| TMFG | ICC.z | 4704 | 0.647 | 0.5696469 |
| PMFG | ICC.z | 4704 | 0.635 | 0.5614855 |
| $\text{PROP}_{10}$ | ICC.z | 4704 | 0.617 | 0.5490358 |
| GCE | ICC.z | 4704 | 0.595 | 0.5334821 |
| $\text{DEG}_{15}$ | ICC.z | 4704 | 0.547 | 0.4982684 |
| $\text{DEG}_{5}$ | ICC.z | 4704 | 0.486 | 0.4510359 |
| ECO | ICC.z | 4704 | 0.431 | 0.4061567 |
| $\text{ABS}_{05}$ | ICC.z | 4704 | 0.337 | 0.3247965 |
| MST | ICC.z | 4704 | 0.319 | 0.3086024 |

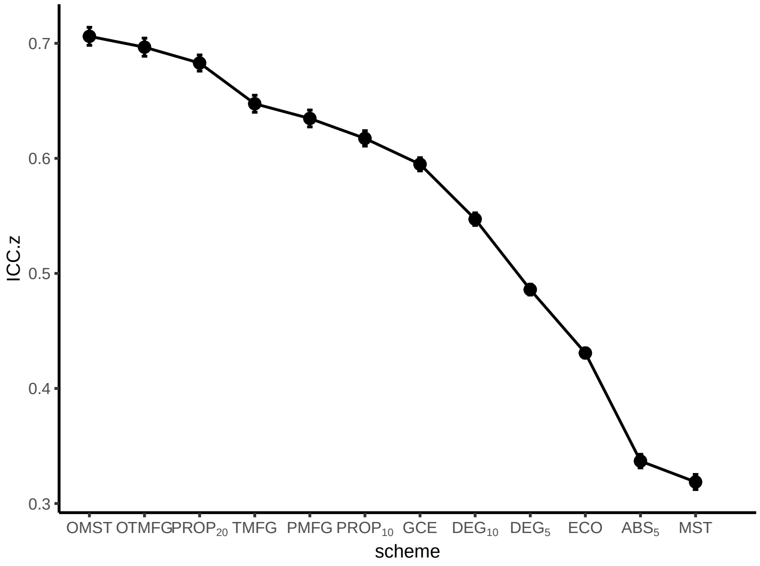

**Fig. S27.** Schemes - ICC mean and se

#### 4.6 Descriptive statistics Median

**Table S26.** Schemes - ICC Median

| scheme | variable | n | median |
|---|---|---|---|
| $PROP_{20}$ | ICC | 4704 | 0.506 |
| OMST | ICC | 4704 | 0.522 |
| OTMFG | ICC | 4704 | 0.507 |
| $PROP_{10}$ | ICC | 4704 | 0.476 |
| GCE | ICC | 4704 | 0.473 |
| PMFG | ICC | 4704 | 0.455 |
| TMFG | ICC | 4704 | 0.455 |
| $DEG_{15}$ | ICC | 4704 | 0.442 |
| $DEG_{5}$ | ICC | 4704 | 0.400 |
| ECO | ICC | 4704 | 0.366 |
| $ABS_{05}$ | ICC | 4704 | 0.223 |
| MST | ICC | 4704 | 0.117 |

#### 4.7 Friedman Test

**Table S27.** Schemes - Friedman Test

| .y. | n | statistic | df | p | method |
|---|---|---|---|---|---|
| ICC.z | 4704 | 9784.317 | 11 | 0 | Friedman test |

#### 4.8 Friedman Test Effect size

**Table S28.** Schemes - Friedman Test Effect size

| .y. | n | effsize | method | magnitude |
|---|---|---|---|---|
| ICC.z | 4704 | 0.1890909 | Kendall W | small |

#### 4.9 Paired Wilcoxon signed rank test

**Table S29.** Schemes - ICC group1 vs. group2

| group1 | group2 | n1 | statistic | alternative | p | p.adj | p.adj.signif |
|---|---|---|---|---|---|---|---|
| $ABS_{05}$ | $DEG_{15}$ | 4704 | 2694254 | two.sided | 0.00e+00 | 0.0000000 | **** |
| $ABS_{05}$ | $DEG_{5}$ | 4704 | 3270500 | two.sided | 0.00e+00 | 0.0000000 | **** |
| $ABS_{05}$ | ECO | 4704 | 3943115 | two.sided | 0.00e+00 | 0.0000000 | **** |
| $ABS_{05}$ | GCE | 4704 | 2271646 | two.sided | 0.00e+00 | 0.0000000 | **** |
| $ABS_{05}$ | MST | 4704 | 4965999 | two.sided | 0.00e+00 | 0.0000003 | **** |
| $ABS_{05}$ | OMST | 4704 | 1467195 | two.sided | 0.00e+00 | 0.0000000 | **** |
| $ABS_{05}$ | OTMFG | 4704 | 1591487 | two.sided | 0.00e+00 | 0.0000000 | **** |
| $ABS_{05}$ | PMFG | 4704 | 2360874 | two.sided | 0.00e+00 | 0.0000000 | **** |

**Table S29.** Schemes - ICC group1 vs. group2 *(continued)*

| group1 | group2 | n1 | statistic | alternative | p | p.adj | p.adj.signif |
|---|---|---|---|---|---|---|---|
| $ABS_{05}$ | $PROP_{10}$ | 4704 | 2285372 | two.sided | 0.00e+00 | 0.0000000 | **** |
| $ABS_{05}$ | $PROP_{20}$ | 4704 | 1672112 | two.sided | 0.00e+00 | 0.0000000 | **** |
| $ABS_{05}$ | TMFG | 4704 | 2232779 | two.sided | 0.00e+00 | 0.0000000 | **** |
| $DEG_{15}$ | $DEG_{5}$ | 4704 | 7803402 | two.sided | 0.00e+00 | 0.0000000 | **** |
| $DEG_{15}$ | ECO | 4704 | 8444214 | two.sided | 0.00e+00 | 0.0000000 | **** |
| $DEG_{15}$ | GCE | 4704 | 4108796 | two.sided | 0.00e+00 | 0.0000000 | **** |
| $DEG_{15}$ | MST | 4704 | 8200729 | two.sided | 0.00e+00 | 0.0000000 | **** |
| $DEG_{15}$ | OMST | 4704 | 2665961 | two.sided | 0.00e+00 | 0.0000000 | **** |
| $DEG_{15}$ | OTMFG | 4704 | 2810801 | two.sided | 0.00e+00 | 0.0000000 | **** |
| $DEG_{15}$ | PMFG | 4704 | 4054470 | two.sided | 0.00e+00 | 0.0000000 | **** |
| $DEG_{15}$ | $PROP_{10}$ | 4704 | 3445204 | two.sided | 0.00e+00 | 0.0000000 | **** |
| $DEG_{15}$ | $PROP_{20}$ | 4704 | 2504887 | two.sided | 0.00e+00 | 0.0000000 | **** |
| $DEG_{15}$ | TMFG | 4704 | 3834624 | two.sided | 0.00e+00 | 0.0000000 | **** |
| $DEG_{5}$ | ECO | 4704 | 8040590 | two.sided | 0.00e+00 | 0.0000000 | **** |
| $DEG_{5}$ | GCE | 4704 | 3328222 | two.sided | 0.00e+00 | 0.0000000 | **** |
| $DEG_{5}$ | MST | 4704 | 7483490 | two.sided | 0.00e+00 | 0.0000000 | **** |
| $DEG_{5}$ | OMST | 4704 | 2135420 | two.sided | 0.00e+00 | 0.0000000 | **** |
| $DEG_{5}$ | OTMFG | 4704 | 2269997 | two.sided | 0.00e+00 | 0.0000000 | **** |
| $DEG_{5}$ | PMFG | 4704 | 2996624 | two.sided | 0.00e+00 | 0.0000000 | **** |
| $DEG_{5}$ | $PROP_{10}$ | 4704 | 2832475 | two.sided | 0.00e+00 | 0.0000000 | **** |
| $DEG_{5}$ | $PROP_{20}$ | 4704 | 1965712 | two.sided | 0.00e+00 | 0.0000000 | **** |
| $DEG_{5}$ | TMFG | 4704 | 2747205 | two.sided | 0.00e+00 | 0.0000000 | **** |
| ECO | GCE | 4704 | 2438845 | two.sided | 0.00e+00 | 0.0000000 | **** |
| ECO | MST | 4704 | 6602202 | two.sided | 0.00e+00 | 0.0000000 | **** |
| ECO | OMST | 4704 | 1445475 | two.sided | 0.00e+00 | 0.0000000 | **** |
| ECO | OTMFG | 4704 | 1561756 | two.sided | 0.00e+00 | 0.0000000 | **** |
| ECO | PMFG | 4704 | 2072641 | two.sided | 0.00e+00 | 0.0000000 | **** |
| ECO | $PROP_{10}$ | 4704 | 2273516 | two.sided | 0.00e+00 | 0.0000000 | **** |
| ECO | $PROP_{20}$ | 4704 | 1494010 | two.sided | 0.00e+00 | 0.0000000 | **** |
| ECO | TMFG | 4704 | 1864579 | two.sided | 0.00e+00 | 0.0000000 | **** |
| GCE | MST | 4704 | 8955851 | two.sided | 0.00e+00 | 0.0000000 | **** |
| GCE | OMST | 4704 | 3276387 | two.sided | 0.00e+00 | 0.0000000 | **** |
| GCE | OTMFG | 4704 | 3459105 | two.sided | 0.00e+00 | 0.0000000 | **** |
| GCE | PMFG | 4704 | 5127650 | two.sided | 2.85e-05 | 0.0018810 | ** |
| GCE | $PROP_{10}$ | 4704 | 5081947 | two.sided | 3.30e-06 | 0.0002158 | *** |
| GCE | $PROP_{20}$ | 4704 | 3034877 | two.sided | 0.00e+00 | 0.0000000 | **** |
| GCE | TMFG | 4704 | 4875617 | two.sided | 0.00e+00 | 0.0000000 | **** |
| MST | OMST | 4704 | 712172 | two.sided | 0.00e+00 | 0.0000000 | **** |
| MST | OTMFG | 4704 | 795758 | two.sided | 0.00e+00 | 0.0000000 | **** |
| MST | PMFG | 4704 | 1340430 | two.sided | 0.00e+00 | 0.0000000 | **** |
| MST | $PROP_{10}$ | 4704 | 1771350 | two.sided | 0.00e+00 | 0.0000000 | **** |
| MST | $PROP_{20}$ | 4704 | 1149079 | two.sided | 0.00e+00 | 0.0000000 | **** |
| MST | TMFG | 4704 | 1164787 | two.sided | 0.00e+00 | 0.0000000 | **** |
| OMST | OTMFG | 4704 | 5863222 | two.sided | 0.00e+00 | 0.0000000 | **** |
| OMST | PMFG | 4704 | 7183474 | two.sided | 0.00e+00 | 0.0000000 | **** |
| OMST | $PROP_{10}$ | 4704 | 7386941 | two.sided | 0.00e+00 | 0.0000000 | **** |
| OMST | $PROP_{20}$ | 4704 | 6196410 | two.sided | 0.00e+00 | 0.0000000 | **** |

**Table S29.** Schemes - ICC group1 vs. group2 *(continued)*

| group1 | group2 | n1 | statistic | alternative | p | p.adj | p.adj.signif |
|---|---|---|---|---|---|---|---|
| OMST | TMFG | 4704 | 6887505 | two.sided | 0.00e+00 | 0.0000000 | **** |
| OTMFG | PMFG | 4704 | 7021466 | two.sided | 0.00e+00 | 0.0000000 | **** |
| OTMFG | $\text{PROP}_{10}$ | 4704 | 7154543 | two.sided | 0.00e+00 | 0.0000000 | **** |
| OTMFG | $\text{PROP}_{20}$ | 4704 | 5995825 | two.sided | 0.00e+00 | 0.0000028 | **** |
| OTMFG | TMFG | 4704 | 6682826 | two.sided | 0.00e+00 | 0.0000000 | **** |
| PMFG | $\text{PROP}_{10}$ | 4704 | 5618652 | two.sided | 1.45e-01 | 1.0000000 | ns |
| PMFG | $\text{PROP}_{20}$ | 4704 | 4515273 | two.sided | 0.00e+00 | 0.0000000 | **** |
| PMFG | TMFG | 4704 | 4119850 | two.sided | 0.00e+00 | 0.0000000 | **** |
| $\text{PROP}_{10}$ | $\text{PROP}_{20}$ | 4704 | 2886243 | two.sided | 0.00e+00 | 0.0000000 | **** |
| $\text{PROP}_{10}$ | TMFG | 4704 | 5169273 | two.sided | 4.31e-04 | 0.0284460 | * |
| $\text{PROP}_{20}$ | TMFG | 4704 | 6289407 | two.sided | 0.00e+00 | 0.0000000 | **** |

#### 4.10 Kullback-leibler divergence Map

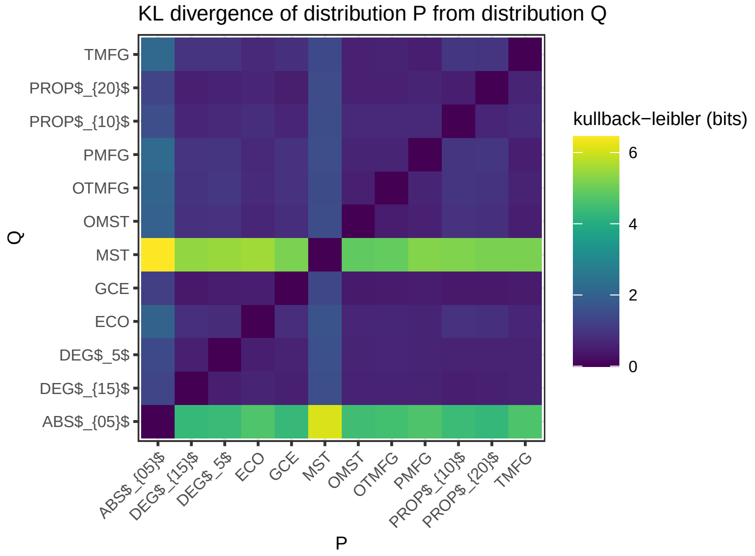

**Fig. S28.** Schemes - KL divergence Map

#### 4.11 Significance map

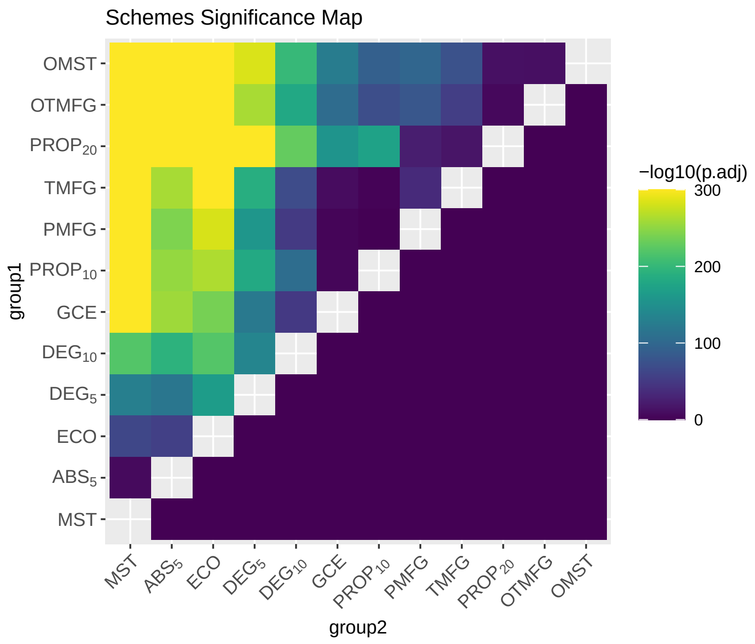

**Fig. S29.** Schemes - Significance Map

#### 4.12 Effect size

**Table S30.** Schemes - Effect size

| group1 | group2 | effsize | n1 | n2 | magnitude |
|---|---|---|---|---|---|
| $\text{ABS}_{05}$ | $\text{DEG}_{15}$ | 0.4336085 | 4704 | 4704 | moderate |
| $\text{ABS}_{05}$ | $\text{DEG}_{5}$ | 0.3385623 | 4704 | 4704 | moderate |
| $\text{ABS}_{05}$ | ECO | 0.2339104 | 4704 | 4704 | small |
| $\text{ABS}_{05}$ | GCE | 0.5006219 | 4704 | 4704 | large |
| $\text{ABS}_{05}$ | MST | 0.1010940 | 4704 | 4704 | small |
| $\text{ABS}_{05}$ | OMST | 0.6272196 | 4704 | 4704 | large |
| $\text{ABS}_{05}$ | OTMFG | 0.6064118 | 4704 | 4704 | large |
| $\text{ABS}_{05}$ | PMFG | 0.4861088 | 4704 | 4704 | moderate |
| $\text{ABS}_{05}$ | $\text{PROP}_{10}$ | 0.4962328 | 4704 | 4704 | moderate |
| $\text{ABS}_{05}$ | $\text{PROP}_{20}$ | 0.5953801 | 4704 | 4704 | large |
| $\text{ABS}_{05}$ | TMFG | 0.5043800 | 4704 | 4704 | large |
| $\text{DEG}_{15}$ | $\text{DEG}_{5}$ | 0.3656085 | 4704 | 4704 | moderate |
| $\text{DEG}_{15}$ | ECO | 0.4643143 | 4704 | 4704 | moderate |
| $\text{DEG}_{15}$ | GCE | 0.2213037 | 4704 | 4704 | small |
| $\text{DEG}_{15}$ | MST | 0.4629302 | 4704 | 4704 | moderate |
| $\text{DEG}_{15}$ | OMST | 0.4439416 | 4704 | 4704 | moderate |
| $\text{DEG}_{15}$ | OTMFG | 0.4196640 | 4704 | 4704 | moderate |
| $\text{DEG}_{15}$ | PMFG | 0.2243101 | 4704 | 4704 | small |
| $\text{DEG}_{15}$ | $\text{PROP}_{10}$ | 0.3230340 | 4704 | 4704 | moderate |

**Table S30.** Schemes - Effect size *(continued)*

| group1 | group2 | effsize | n1 | n2 | magnitude |
|---|---|---|---|---|---|
| $DEG_{15}$ | $PROP_{20}$ | 0.4729914 | 4704 | 4704 | moderate |
| $DEG_{15}$ | TMFG | 0.2592393 | 4704 | 4704 | small |
| $DEG_{5}$ | ECO | 0.4011082 | 4704 | 4704 | moderate |
| $DEG_{5}$ | GCE | 0.3443975 | 4704 | 4704 | moderate |
| $DEG_{5}$ | MST | 0.3530459 | 4704 | 4704 | moderate |
| $DEG_{5}$ | OMST | 0.5266713 | 4704 | 4704 | large |
| $DEG_{5}$ | OTMFG | 0.5058619 | 4704 | 4704 | large |
| $DEG_{5}$ | PMFG | 0.3921309 | 4704 | 4704 | moderate |
| $DEG_{5}$ | $PROP_{10}$ | 0.4209417 | 4704 | 4704 | moderate |
| $DEG_{5}$ | $PROP_{20}$ | 0.5574678 | 4704 | 4704 | large |
| $DEG_{5}$ | TMFG | 0.4294553 | 4704 | 4704 | moderate |
| ECO | GCE | 0.4829184 | 4704 | 4704 | moderate |
| ECO | MST | 0.2472343 | 4704 | 4704 | small |
| ECO | OMST | 0.6246376 | 4704 | 4704 | large |
| ECO | OTMFG | 0.6073007 | 4704 | 4704 | large |
| ECO | PMFG | 0.5251605 | 4704 | 4704 | large |
| ECO | $PROP_{10}$ | 0.5082960 | 4704 | 4704 | large |
| ECO | $PROP_{20}$ | 0.6300488 | 4704 | 4704 | large |
| ECO | TMFG | 0.5579538 | 4704 | 4704 | large |
| GCE | MST | 0.5682424 | 4704 | 4704 | large |
| GCE | OMST | 0.3497301 | 4704 | 4704 | moderate |
| GCE | OTMFG | 0.3209678 | 4704 | 4704 | moderate |
| GCE | PMFG | 0.0609041 | 4704 | 4704 | small |
| GCE | $PROP_{10}$ | 0.0677587 | 4704 | 4704 | small |
| GCE | $PROP_{20}$ | 0.3900709 | 4704 | 4704 | moderate |
| GCE | TMFG | 0.0968897 | 4704 | 4704 | small |
| MST | OMST | 0.7314210 | 4704 | 4704 | large |
| MST | OTMFG | 0.7144886 | 4704 | 4704 | large |
| MST | PMFG | 0.6232274 | 4704 | 4704 | large |
| MST | $PROP_{10}$ | 0.5694680 | 4704 | 4704 | large |
| MST | $PROP_{20}$ | 0.6742707 | 4704 | 4704 | large |
| MST | TMFG | 0.6506873 | 4704 | 4704 | large |
| OMST | OTMFG | 0.1072281 | 4704 | 4704 | small |
| OMST | PMFG | 0.3108939 | 4704 | 4704 | moderate |
| OMST | $PROP_{10}$ | 0.2977166 | 4704 | 4704 | small |
| OMST | $PROP_{20}$ | 0.1111673 | 4704 | 4704 | small |
| OMST | TMFG | 0.2713060 | 4704 | 4704 | small |
| OTMFG | PMFG | 0.2809687 | 4704 | 4704 | small |
| OTMFG | $PROP_{10}$ | 0.2638212 | 4704 | 4704 | small |
| OTMFG | $PROP_{20}$ | 0.0798667 | 4704 | 4704 | small |
| OTMFG | TMFG | 0.2305216 | 4704 | 4704 | small |

**Table S30.** Schemes - Effect size *(continued)*

| group1 | group2 | effsize | n1 | n2 | magnitude |
|---|---|---|---|---|---|
| PMFG | $\text{PROP}_{10}$ | 0.0210514 | 4704 | 4704 | small |
| PMFG | $\text{PROP}_{20}$ | 0.1549509 | 4704 | 4704 | small |
| PMFG | TMFG | 0.1861855 | 4704 | 4704 | small |
| $\text{PROP}_{10}$ | $\text{PROP}_{20}$ | 0.4119006 | 4704 | 4704 | moderate |
| $\text{PROP}_{10}$ | TMFG | 0.0511920 | 4704 | 4704 | small |
| $\text{PROP}_{20}$ | TMFG | 0.1263285 | 4704 | 4704 | small |

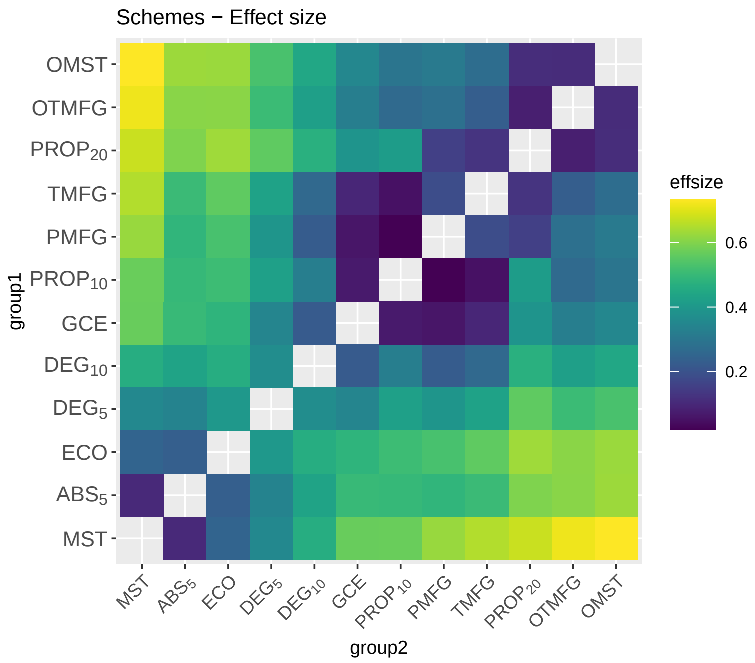

**Fig. S30.** Schemes - Effect size

### 5 Metrics - Network Analysis

#### 5.1 Metrics

Superscript $b$ (eg. $Eg^b$) is used for the binary graphs, superscript $w$ (eg. $Eg^w$) is used for the weighted graphs, and superscript $n$ (eg. $Eg^n$) is used for the normalized weighted graphs.

**Table S31.** Brief descriptions of the network metrics examined

| Level | Measure | Attribute | Character |
|---|---|---|---|
| Global | basic | degree | $k$ |
| | | number of edge | $n_e$ |
| | | density | $d$ |
| | integration | global efficiency | $Eg$ |
| | | average shortest path length | $Lp_a$ |
| | | average nodal path length | $Lp_b$ |
| | | pseudo diameter | $D$ |
| | segregation | clusteringcoef by BCT toolbox | $Cp_a$ |
| | | clusteringcoef by Graph-tool | $Cp_b$ |
| | | local efficiency algorithm 1 by BCT toolbox | $Eloc_1$ |
| | | local efficiency algorithm 2 by BCT toolbox | $Eloc_2$ |
| | | modularity | $Q$ |
| | | transitivity by BCT toolbox | $Tr_a$ |
| | | transitivity by Graph-tool | $Tr_b$ |
| | centrality | average betweenness | $Bc$ |
| | | average eigenvector | $Ec$ |
| | | average pagerank | $Pc$ |
| | | average subgraph | $Sc$ |
| | resilience | assortativity | $r$ |
| | | scalar assortativity | $r_s$ |
| | | synchronizability | $S$ |
| | | average resolvent | $Rv$ |
| Nodal | integration | nodal path length | $Lp_a$ |
| | | local path length | $Lp_b$ |
| | segregation | clustering coefficient by BCT toolbox | $Cp_a$ |
| | | clustering coefficient Graph-tool | $Cp_b$ |
| | | local efficiency 1 by BCT toolbox | $Eloc_1$ |
| | | local efficiency 2 by BCT toolbox | $Eloc_2$ |
| | | nodal efficiency | $E_{nodal}$ |
| | centrality | degree centrality | $Dc$ |
| | | betweeness centrality | $Bc$ |
| | | eigenvector centrality | $Ec$ |
| | | pagerank centrality | $Pc$ |
| | | subgraph centrality | $Sc$ |
| | | resolvent centrality | $Rc$ |

#### 5.2 ICC Density distribution

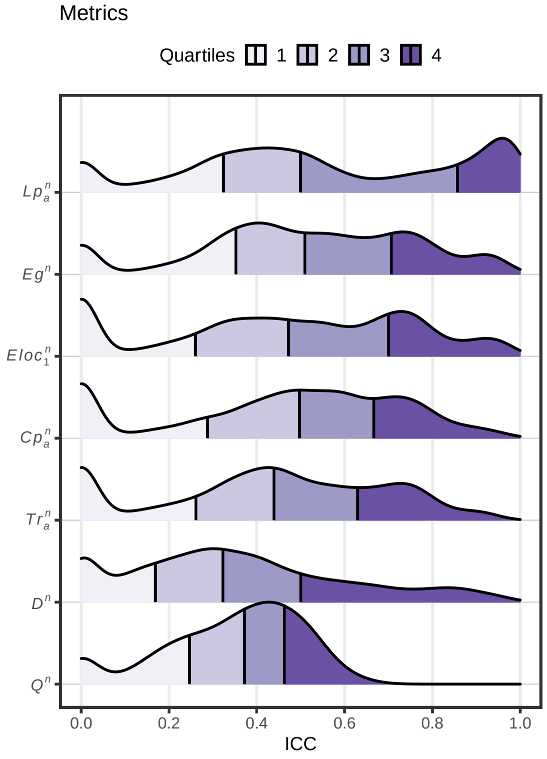

**Fig. S31.** Metrics - Density distribution

#### 5.3 Almost Perfect ( ICCs > 0.8 )

**Table S32.** Metrics - Number of ICCs > 0.8

| | character | n |
|---|---|---|
| 3 | $Lp_a^n$ | 2350 |
| 4 | $Eg^n$ | 924 |
| 2 | $Eloc_1^n$ | 835 |
| 1 | $Cp_a^n$ | 575 |
| 5 | $D^n$ | 559 |
| 6 | $Tr_a^n$ | 433 |

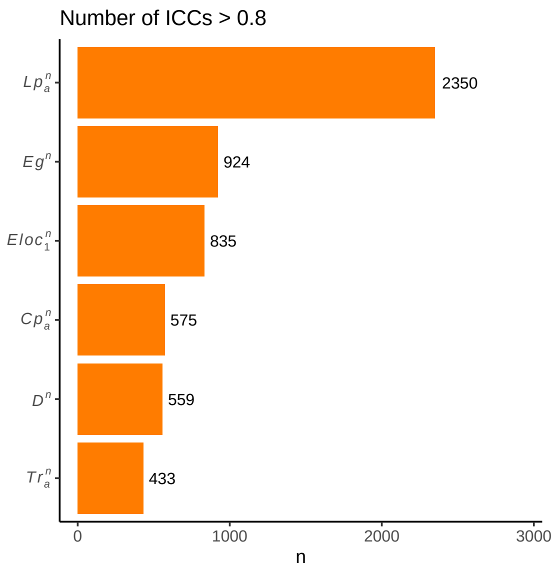

**Fig. S32.** Metrics - Number of ICC > 0.8

#### 5.4 Substantial or Above ( ICCs > 0.6 )

**Table S33.** Metrics - Number of ICCs > 0.6

| | character | n |
|---|---|---|
| 3 | $Lp_a^n$ | 3194 |
| 4 | $Eg^n$ | 3032 |
| 2 | $Eloc_1^n$ | 2886 |
| 1 | $Cp_a^n$ | 2709 |
| 7 | $Tr_a^n$ | 2273 |
| 6 | $D^n$ | 1383 |
| 5 | $Q^n$ | 140 |

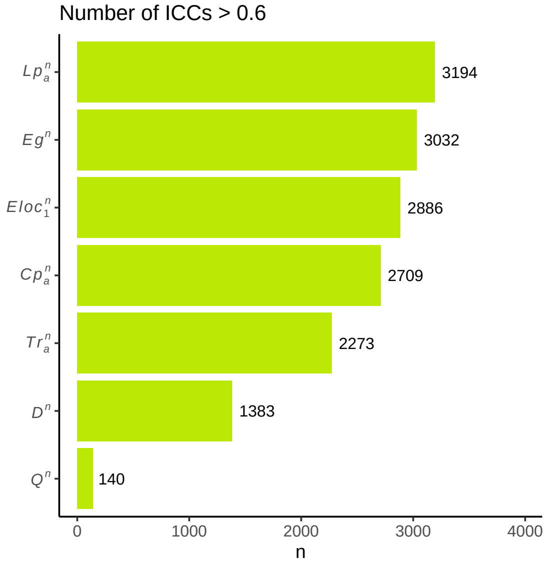

**Fig. S33.** Metrics - Number of ICC > 0.6

#### 5.5 Descriptive statistics Mean

**Table S34.** Metrics - ICC Mean

| character | n | meanz2r |
|---|---|---|
| $Lp_a^n$ | 8064 | 0.6847483 |
| $Eg^n$ | 8064 | 0.5628534 |
| $Eloc_1^n$ | 8064 | 0.5241176 |
| $Cp_a^n$ | 8064 | 0.5057482 |
| $Tr_a^n$ | 8064 | 0.4683854 |
| $D^n$ | 8064 | 0.3952442 |
| $Q^n$ | 8064 | 0.3531167 |

#### 5.6 Descriptive statistics Median

**Table S35.** Metrics - ICC Median

| character | n | median |
|---|---|---|
| $Eg^n$ | 8064 | 0.510 |
| $Lp_a^n$ | 8064 | 0.499 |
| $Cp_a^n$ | 8064 | 0.497 |
| $Eloc_1^n$ | 8064 | 0.472 |
| $Tr_a^n$ | 8064 | 0.439 |
| $Q^n$ | 8064 | 0.371 |
| $D^n$ | 8064 | 0.323 |

#### 5.7 Paired Wilcoxon signed rank test

**Table S36.** Metrics - ICC group1 vs. group2

| character1 | character2 | n1 | statistic | alternative | p | p.adj | p.adj.signif |
|---|---|---|---|---|---|---|---|
| $Cp_a^n$ | $Eloc_1^n$ | 8064 | 12227711 | two.sided | 9.20e-06 | 0.0001934 | *** |
| $Cp_a^n$ | $Lp_a^n$ | 8064 | 8726636 | two.sided | 0.00e+00 | 0.0000000 | **** |
| $Cp_a^n$ | $Eg^n$ | 8064 | 13406103 | two.sided | 0.00e+00 | 0.0000000 | **** |
| $Cp_a^n$ | $Q^n$ | 8064 | 23845905 | two.sided | 0.00e+00 | 0.0000000 | **** |
| $Cp_a^n$ | $D^n$ | 8064 | 23503271 | two.sided | 0.00e+00 | 0.0000000 | **** |
| $Cp_a^n$ | $Tr_a^n$ | 8064 | 18879595 | two.sided | 0.00e+00 | 0.0000000 | **** |
| $Eloc_1^n$ | $Lp_a^n$ | 8064 | 8214982 | two.sided | 0.00e+00 | 0.0000000 | **** |
| $Eloc_1^n$ | $Eg^n$ | 8064 | 13804539 | two.sided | 0.00e+00 | 0.0000000 | **** |
| $Eloc_1^n$ | $Q^n$ | 8064 | 23296271 | two.sided | 0.00e+00 | 0.0000000 | **** |
| $Eloc_1^n$ | $D^n$ | 8064 | 24208558 | two.sided | 0.00e+00 | 0.0000000 | **** |
| $Eloc_1^n$ | $Tr_a^n$ | 8064 | 17771904 | two.sided | 0.00e+00 | 0.0000000 | **** |
| $Lp_a^n$ | $Eg^n$ | 8064 | 21277439 | two.sided | 0.00e+00 | 0.0000000 | **** |
| $Lp_a^n$ | $Q^n$ | 8064 | 26286171 | two.sided | 0.00e+00 | 0.0000000 | **** |
| $Lp_a^n$ | $D^n$ | 8064 | 28471330 | two.sided | 0.00e+00 | 0.0000000 | **** |
| $Lp_a^n$ | $Tr_a^n$ | 8064 | 24806988 | two.sided | 0.00e+00 | 0.0000000 | **** |
| $Eg^n$ | $Q^n$ | 8064 | 26666127 | two.sided | 0.00e+00 | 0.0000000 | **** |
| $Eg^n$ | $D^n$ | 8064 | 27744888 | two.sided | 0.00e+00 | 0.0000000 | **** |
| $Eg^n$ | $Tr_a^n$ | 8064 | 21280096 | two.sided | 0.00e+00 | 0.0000000 | **** |
| $Q^n$ | $D^n$ | 8064 | 15808230 | two.sided | 3.65e-01 | 1.0000000 | ns |
| $Q^n$ | $Tr_a^n$ | 8064 | 9856527 | two.sided | 0.00e+00 | 0.0000000 | **** |
| $D^n$ | $Tr_a^n$ | 8064 | 8856878 | two.sided | 0.00e+00 | 0.0000000 | **** |

#### 5.8 Kullback-leibler divergence Map

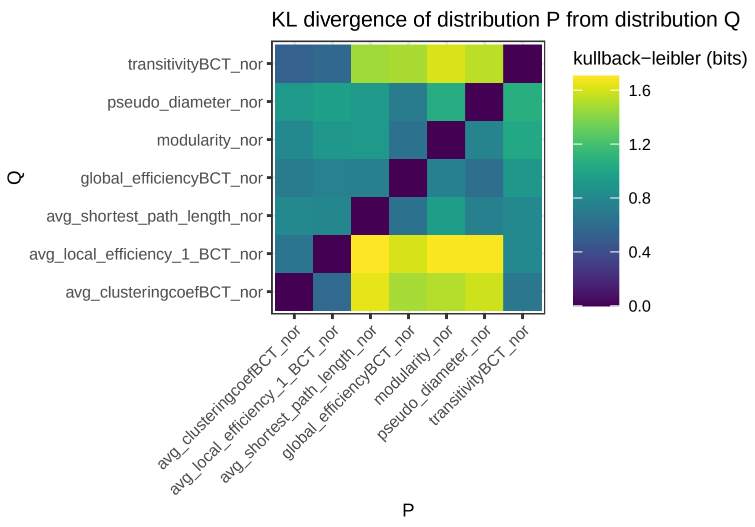

**Fig. S34.** Metrics - KL divergence Map

#### 5.9 Significance map

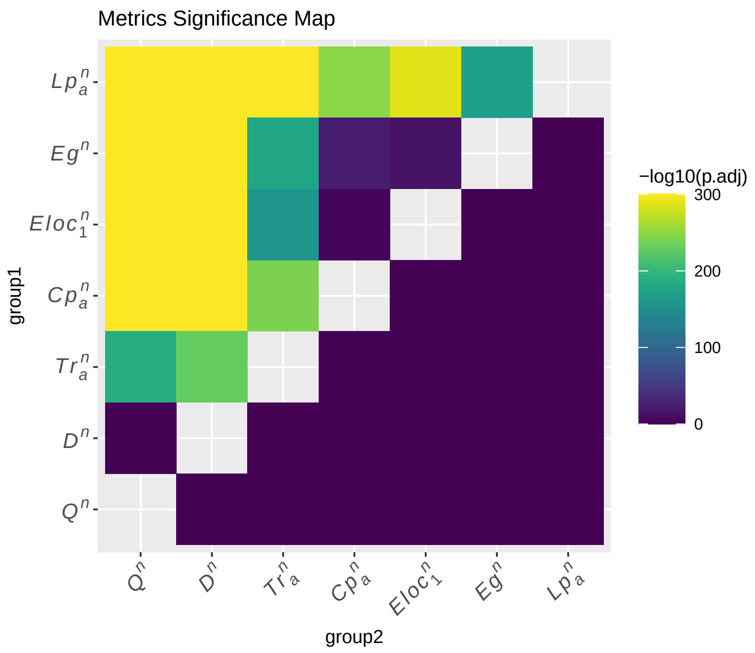

**Fig. S35.** Metrics - Significance Map

#### 5.10 Effect size

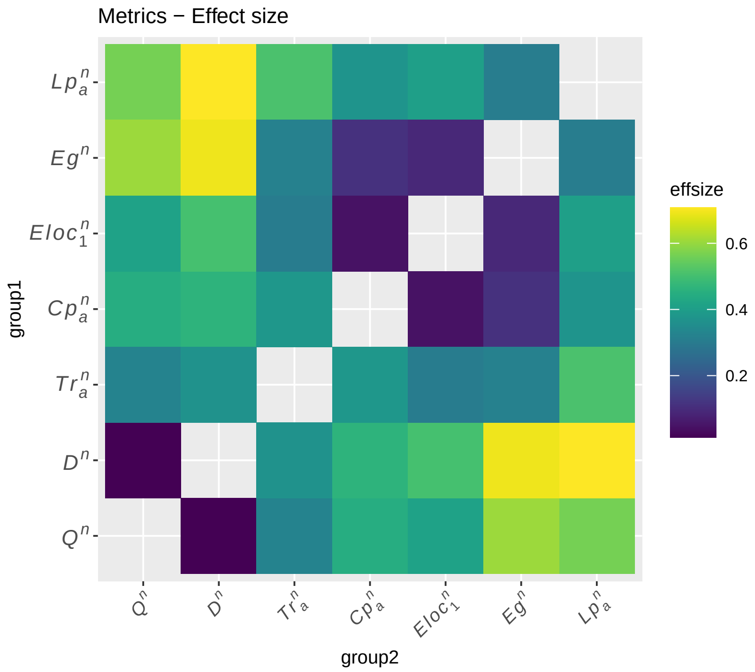

**Fig. S36.** Metrics - Effect size

**Table S37.** Metrics - Effect size

| character1 | character2 | effsize | n1 | n2 | magnitude |
|---|---|---|---|---|---|
| $Cp_a^n$ | $Eloc_1^n$ | 0.0442149 | 8064 | 8064 | small |
| $Cp_a^n$ | $Lp_a^n$ | 0.3729141 | 8064 | 8064 | moderate |
| $Cp_a^n$ | $Eg^n$ | 0.1094179 | 8064 | 8064 | small |
| $Cp_a^n$ | $Q^n$ | 0.4462449 | 8064 | 8064 | moderate |
| $Cp_a^n$ | $D^n$ | 0.4629948 | 8064 | 8064 | moderate |
| $Cp_a^n$ | $Tr_a^n$ | 0.3823045 | 8064 | 8064 | moderate |
| $Eloc_1^n$ | $Lp_a^n$ | 0.4014701 | 8064 | 8064 | moderate |
| $Eloc_1^n$ | $Eg^n$ | 0.0895200 | 8064 | 8064 | small |
| $Eloc_1^n$ | $Q^n$ | 0.4124881 | 8064 | 8064 | moderate |
| $Eloc_1^n$ | $D^n$ | 0.5027420 | 8064 | 8064 | large |
| $Eloc_1^n$ | $Tr_a^n$ | 0.3025436 | 8064 | 8064 | moderate |
| $Lp_a^n$ | $Eg^n$ | 0.3055529 | 8064 | 8064 | moderate |
| $Lp_a^n$ | $Q^n$ | 0.5641222 | 8064 | 8064 | large |
| $Lp_a^n$ | $D^n$ | 0.7079035 | 8064 | 8064 | large |
| $Lp_a^n$ | $Tr_a^n$ | 0.5101946 | 8064 | 8064 | large |
| $Eg^n$ | $Q^n$ | 0.6054757 | 8064 | 8064 | large |
| $Eg^n$ | $D^n$ | 0.6915405 | 8064 | 8064 | large |
| $Eg^n$ | $Tr_a^n$ | 0.3167091 | 8064 | 8064 | moderate |
| $Q^n$ | $D^n$ | 0.0132182 | 8064 | 8064 | small |
| $Q^n$ | $Tr_a^n$ | 0.3249259 | 8064 | 8064 | moderate |
| $D^n$ | $Tr_a^n$ | 0.3637976 | 8064 | 8064 | moderate |

### 6 More Metrics - Network Analysis

#### 6.1 ICC Density distribution

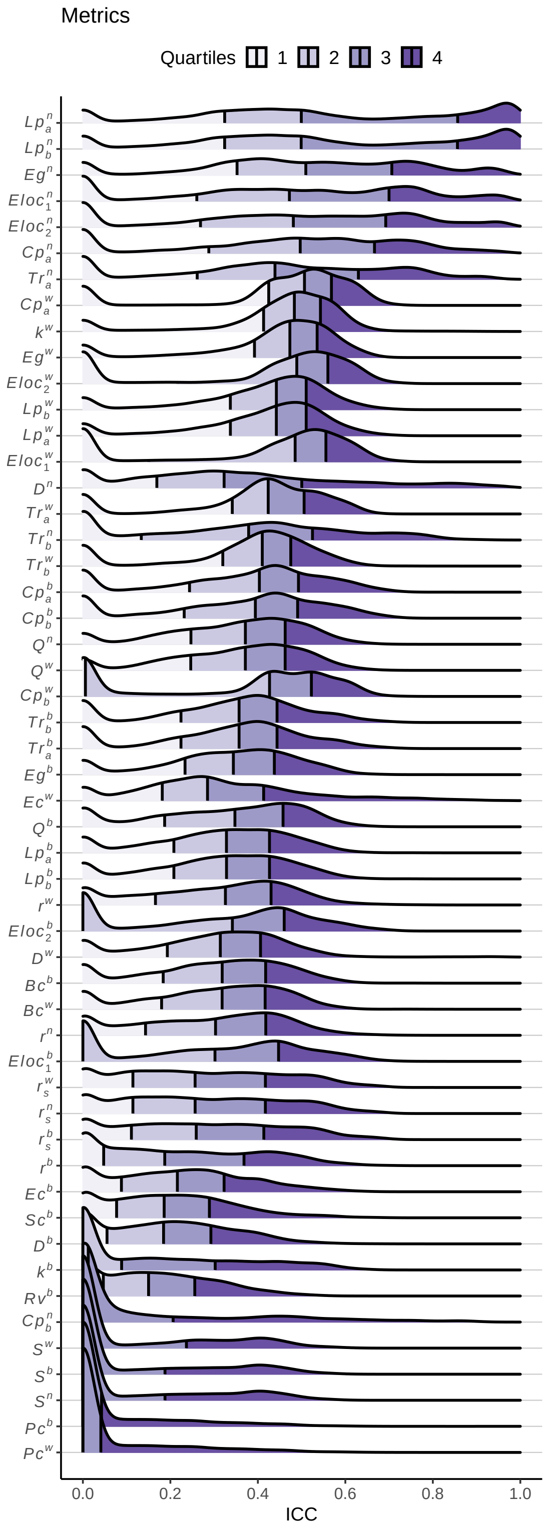

**Fig. S37.** Metrics - Density distribution

#### 6.2 Almost Perfect ( ICCs > 0.8 )

**Table S38.** Metrics - Number of ICCs > 0.8

| | character | n |
|---|---|---|
| 14 | $Lp_a^n$ | 2350 |
| 11 | $Lp_b^n$ | 2349 |
| 16 | $Eg^n$ | 924 |
| 8 | $Eloc_1^n$ | 835 |
| 9 | $Eloc_2^n$ | 792 |
| 5 | $Cp_a^n$ | 575 |
| 17 | $D^n$ | 559 |
| 21 | $Tr_a^n$ | 433 |
| 6 | $Ec^w$ | 137 |
| 19 | $Tr_b^n$ | 131 |
| 4 | $Cp_b^n$ | 114 |
| 18 | $D^w$ | 57 |
| 15 | $k^w$ | 27 |
| 12 | $Pc^b$ | 4 |
| 1 | $r^w$ | 2 |
| 2 | $Bc^b$ | 1 |
| 3 | $Bc^w$ | 1 |
| 7 | $Eloc_1^b$ | 1 |
| 10 | $Lp_b^b$ | 1 |
| 13 | $Lp_a^b$ | 1 |
| 20 | $Tr_b^w$ | 1 |

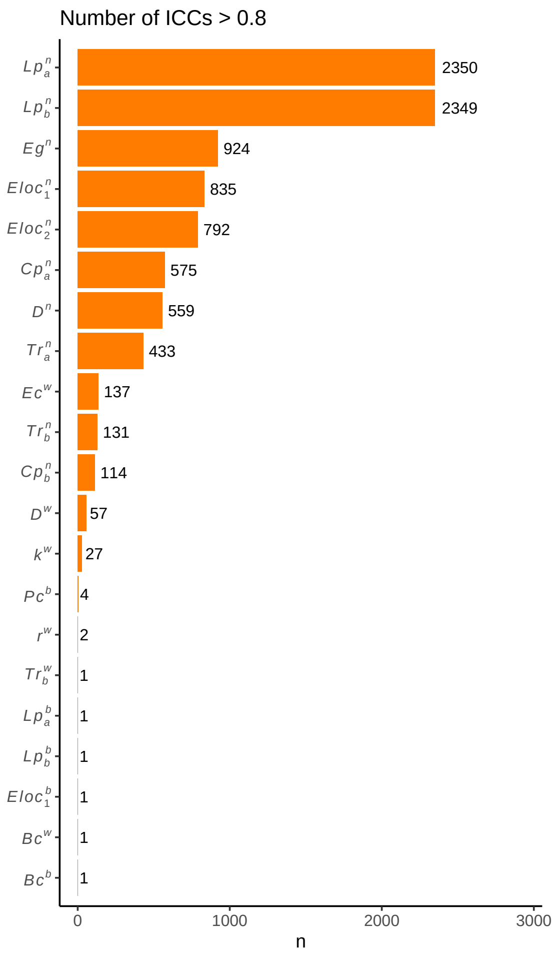

**Fig. S38.** Metrics - Number of ICC > 0.8

#### 6.3 Substantial or Above ( ICCs > 0.6 )

**Table S39.** Metrics - Number of ICCs > 0.6

| | character | n |
|---|---|---|
| 27 | $Lp_a^n$ | 3194 |
| 21 | $Lp_b^n$ | 3190 |
| 33 | $Eg^n$ | 3032 |
| 18 | $Eloc_2^n$ | 2913 |
| 15 | $Eloc_1^n$ | 2886 |
| 10 | $Cp_a^n$ | 2709 |
| 51 | $Tr_a^n$ | 2273 |
| 39 | $D^n$ | 1383 |
| 48 | $Tr_b^n$ | 1316 |
| 11 | $Cp_a^w$ | 1302 |
| 19 | $Eloc_2^w$ | 1085 |
| 16 | $Eloc_1^w$ | 998 |
| 13 | $Ec^w$ | 664 |

**Table S39.** Metrics - Number of ICCs > 0.6 *(continued)*

| | character | n |
|---|---|---|
| 9 | $Cp_a^b$ | 603 |
| 6 | $Cp_b^b$ | 595 |
| 8 | $Cp_b^w$ | 592 |
| 31 | $k^w$ | 481 |
| 34 | $Eg^w$ | 467 |
| 7 | $Cp_b^n$ | 463 |
| 52 | $Tr_a^w$ | 385 |
| 22 | $Lp_b^w$ | 375 |
| 28 | $Lp_a^w$ | 375 |
| 17 | $Eloc_2^b$ | 345 |
| 14 | $Eloc_1^b$ | 314 |
| 47 | $Tr_b^b$ | 276 |
| 50 | $Tr_a^b$ | 276 |
| 41 | $r_s^b$ | 247 |
| 42 | $r_s^n$ | 229 |
| 43 | $r_s^w$ | 229 |
| 49 | $Tr_b^w$ | 153 |
| 3 | $r^w$ | 148 |
| 36 | $Q^n$ | 140 |
| 37 | $Q^w$ | 140 |
| 35 | $Q^b$ | 109 |
| 40 | $D^w$ | 107 |
| 2 | $r^n$ | 100 |
| 32 | $Eg^b$ | 83 |
| 1 | $r^b$ | 81 |
| 20 | $Lp_b^b$ | 67 |
| 26 | $Lp_a^b$ | 67 |
| 30 | $k^b$ | 64 |
| 4 | $Bc^b$ | 50 |
| 12 | $Ec^b$ | 40 |
| 29 | $Sc^b$ | 40 |
| 5 | $Bc^w$ | 32 |
| 23 | $Pc^b$ | 18 |
| 38 | $D^b$ | 16 |
| 24 | $Pc^w$ | 14 |
| 46 | $S^w$ | 12 |
| 25 | $Rv^b$ | 2 |
| 44 | $S^b$ | 2 |
| 45 | $S^n$ | 2 |

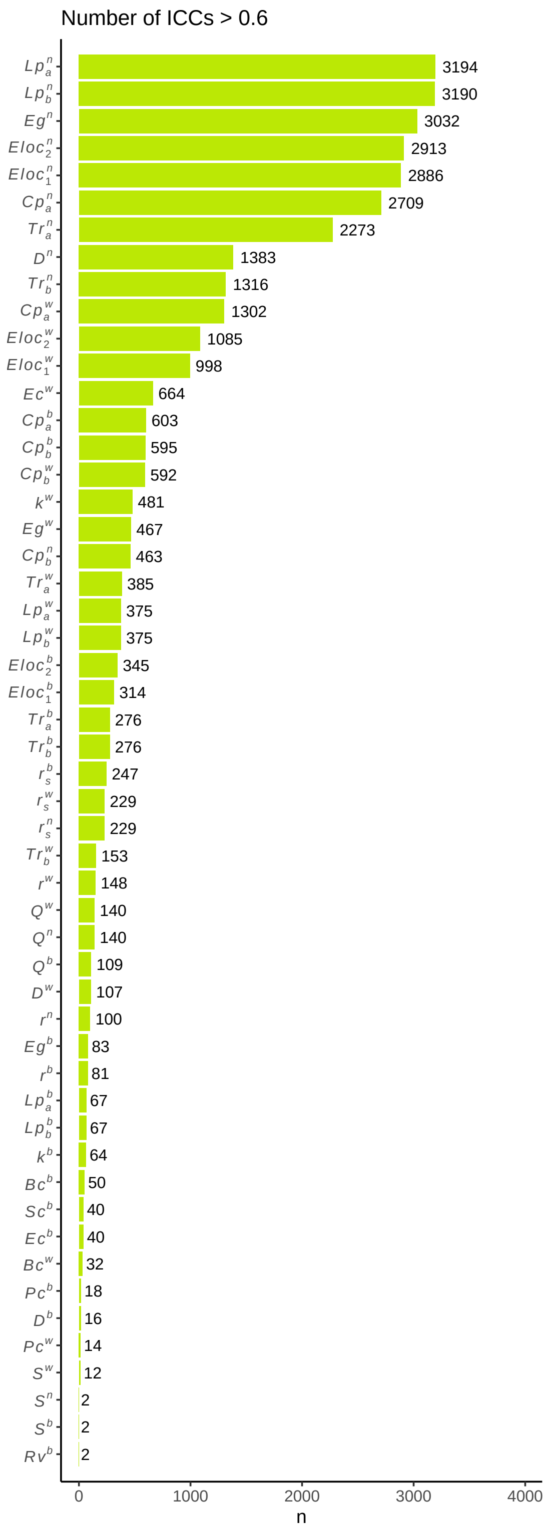

**Fig. S39.** Metrics - Number of ICC > 0.6

#### 6.4 Descriptive statistics Mean

**Table S40.** Metrics - ICC Mean

| character | n | meanz2r |
|---|---|---|
| $Lp_a^n$ | 8064 | 0.6847483 |
| $Lp_b^n$ | 8064 | 0.6842168 |
| $Eg^n$ | 8064 | 0.5628534 |
| $Eloc_1^n$ | 8064 | 0.5241176 |
| $Eloc_2^n$ | 8064 | 0.5219382 |
| $Cp_a^n$ | 8064 | 0.5057482 |
| $Tr_a^n$ | 8064 | 0.4683854 |
| $Cp_a^w$ | 8064 | 0.4621172 |
| $k^w$ | 8064 | 0.4542164 |
| $Eg^w$ | 8064 | 0.4381993 |
| $Eloc_2^w$ | 8064 | 0.4144730 |
| $Lp_b^w$ | 8064 | 0.4094914 |
| $Lp_a^w$ | 8064 | 0.4094914 |
| $Eloc_1^w$ | 8064 | 0.4053213 |
| $D^n$ | 8064 | 0.3952442 |
| $Tr_a^w$ | 8064 | 0.3935553 |
| $Tr_b^n$ | 8064 | 0.3790930 |
| $Tr_b^w$ | 8064 | 0.3704978 |
| $Cp_a^b$ | 8064 | 0.3679068 |
| $Cp_b^b$ | 8064 | 0.3627075 |
| $Q^n$ | 8064 | 0.3531167 |
| $Q^w$ | 8064 | 0.3531167 |
| $Cp_b^w$ | 8064 | 0.3487325 |
| $Tr_b^b$ | 8064 | 0.3346006 |
| $Tr_a^b$ | 8064 | 0.3346006 |
| $Eg^b$ | 8064 | 0.3301530 |
| $Ec^w$ | 8064 | 0.3265843 |
| $Q^b$ | 8064 | 0.3239017 |
| $Lp_b^b$ | 8064 | 0.3140209 |
| $Lp_a^b$ | 8064 | 0.3140209 |
| $r^w$ | 8064 | 0.3067918 |
| $Eloc_2^b$ | 8064 | 0.3049789 |
| $D^w$ | 8064 | 0.3031638 |
| $Bc^b$ | 8064 | 0.3013466 |
| $Bc^w$ | 8064 | 0.2986165 |
| $r^n$ | 8064 | 0.2885648 |
| $Eloc_1^b$ | 8064 | 0.2858122 |
| $r_s^n$ | 8064 | 0.2802930 |
| $r_s^w$ | 8064 | 0.2802930 |
| $r_s^b$ | 8064 | 0.2784491 |
| $r^b$ | 8064 | 0.2231797 |

**Table S40.** Metrics - ICC Mean *(continued)*

| character | n | meanz2r |
|---|---|---|
| $Ec^b$ | 8064 | 0.2222293 |
| $Sc^b$ | 8064 | 0.1983362 |
| $D^b$ | 8064 | 0.1916023 |
| $k^b$ | 8064 | 0.1712945 |
| $Rv^b$ | 8064 | 0.1664371 |
| $Cp_b^n$ | 8064 | 0.1566982 |
| $S^w$ | 8064 | 0.1135087 |
| $S^b$ | 8064 | 0.1036267 |
| $S^n$ | 8064 | 0.1036267 |
| $Pc^b$ | 8064 | 0.0629168 |
| $Pc^w$ | 8064 | 0.0619207 |

#### 6.5 Descriptive statistics Median

**Table S41.** Metrics - ICC Median

| character | n | median |
|---|---|---|
| $Eg^n$ | 8064 | 0.510 |
| $Cp_a^w$ | 8064 | 0.507 |
| $Lp_b^n$ | 8064 | 0.499 |
| $Lp_a^n$ | 8064 | 0.499 |
| $Cp_a^n$ | 8064 | 0.497 |
| $Eloc_2^w$ | 8064 | 0.489 |
| $Eloc_1^w$ | 8064 | 0.485 |
| $k^w$ | 8064 | 0.483 |
| $Eloc_2^n$ | 8064 | 0.481 |
| $Eg^w$ | 8064 | 0.473 |
| $Eloc_1^n$ | 8064 | 0.472 |
| $Lp_b^w$ | 8064 | 0.442 |
| $Lp_a^w$ | 8064 | 0.442 |
| $Tr_a^n$ | 8064 | 0.439 |
| $Cp_b^w$ | 8064 | 0.427 |
| $Tr_a^w$ | 8064 | 0.424 |
| $Tr_b^w$ | 8064 | 0.410 |
| $Cp_a^b$ | 8064 | 0.403 |
| $Cp_b^b$ | 8064 | 0.394 |
| $Tr_b^n$ | 8064 | 0.379 |
| $Q^n$ | 8064 | 0.371 |
| $Q^w$ | 8064 | 0.371 |
| $Tr_b^b$ | 8064 | 0.357 |
| $Tr_a^b$ | 8064 | 0.357 |

**Table S41.** Metrics - ICC Median *(continued)*

| character | n | median |
|---|---|---|
| $Q^b$ | 8064 | 0.348 |
| $Eg^b$ | 8064 | 0.344 |
| $Eloc_2^b$ | 8064 | 0.342 |
| $Lp_b^b$ | 8064 | 0.328 |
| $Lp_a^b$ | 8064 | 0.328 |
| $r^w$ | 8064 | 0.326 |
| $D^n$ | 8064 | 0.323 |
| $Bc^b$ | 8064 | 0.318 |
| $Bc^w$ | 8064 | 0.318 |
| $D^w$ | 8064 | 0.314 |
| $r^n$ | 8064 | 0.303 |
| $Eloc_1^b$ | 8064 | 0.302 |
| $Ec^w$ | 8064 | 0.285 |
| $r_s^b$ | 8064 | 0.259 |
| $r_s^n$ | 8064 | 0.257 |
| $r_s^w$ | 8064 | 0.257 |
| $Ec^b$ | 8064 | 0.216 |
| $r^b$ | 8064 | 0.187 |
| $Sc^b$ | 8064 | 0.186 |
| $D^b$ | 8064 | 0.184 |
| $Rv^b$ | 8064 | 0.150 |
| $k^b$ | 8064 | 0.089 |
| $Cp_b^n$ | 8064 | 0.012 |
| $Pc^b$ | 8064 | 0.000 |
| $Pc^w$ | 8064 | 0.000 |
| $S^b$ | 8064 | 0.000 |
| $S^n$ | 8064 | 0.000 |
| $S^w$ | 8064 | 0.000 |

#### 6.6 Kullback-leibler divergence Map

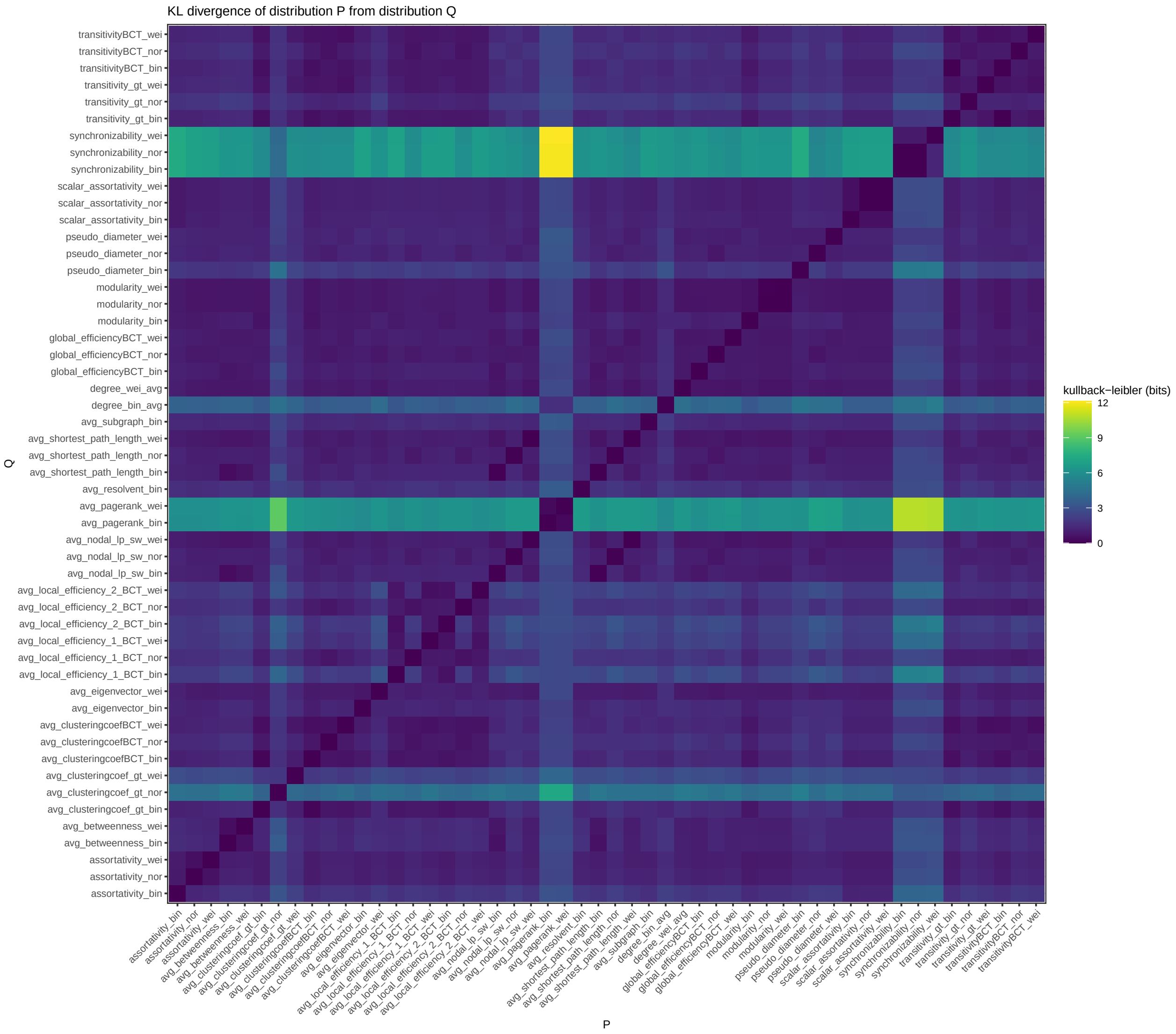

**Fig. S40.** Metrics - KL divergence Map

#### 6.7 Significance map

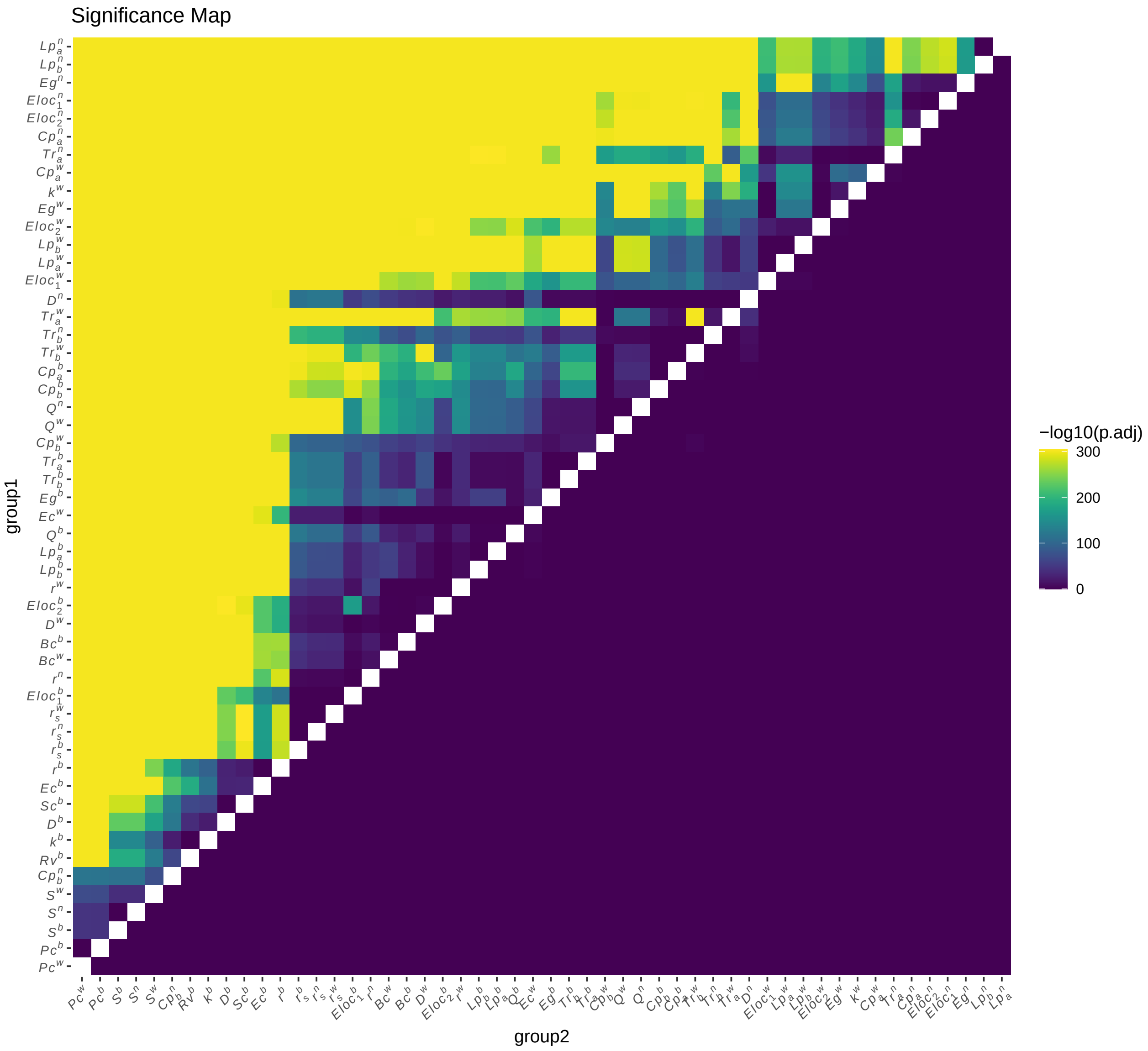

**Fig. S41.** Metrics - Significance Map

### 7 fMRIPrep pipeline

We generated the HCP data preprocessed by the fMRI-Prep pipeline and applied our analytical workflow to the preprocessed data. According to the high computational cost and resource intensity of fully computing the results using this pipeline, as well as the large number of conditions derived from the approximately 40+ subjects and the use of threshold-based methods such as GCE, we were only able to study the results generated by the fMRIPrep pipeline to a limited extent but the general patterns of the findings are reproducible.

Our analysis of the fMRIPrep results showed a similar gradient of changes across the frequency bands (Figure S42), but the whole-brain results were not significantly superior to the cortical results. The figure shows that the ICC values generally increase with higher frequency bands (slow6 to slow2, with slow1 not fully shown), consistent with the results in the original study. However, the differences between parcellations are not significant, with no clear advantage of whole brain over cortex, which may be related to the variable

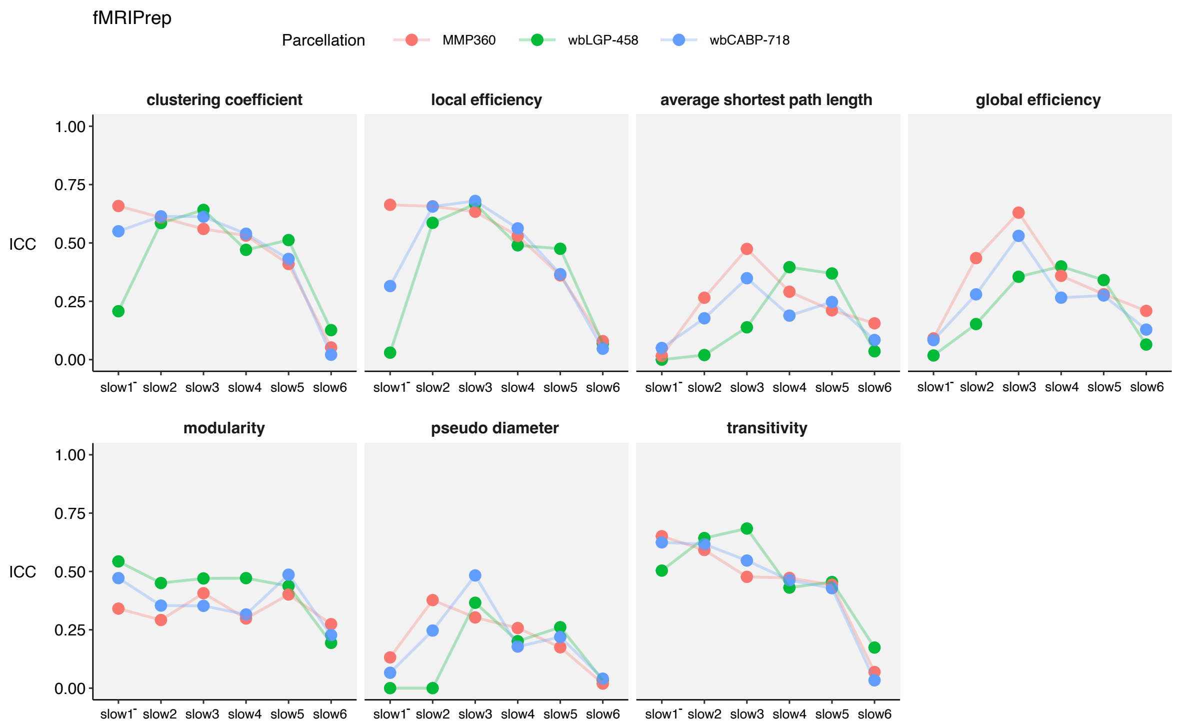

**Fig. S42. Test-retest reliability of fMRIPrep-preprocessed data with different frequency bands and parcellations.** The ICC values of global graph measures including average shortest path length ($L_p$), global efficiency ($E_g$), pseudo diameter ($D$), clustering coefficient ($C_p$), local efficiency ($E_{\text{loc}}$), transitivity ($Tr$), modularity ($Q$) under sampling conditions (pos and OMST) in different frequency bands (slow6 to slow1) and parcellations (MMP-360, wbLGP-458, wbCABP-718) are plotted. The dots are color-coded according to the parcellations: red for MMP-360, green for wbLGP-458, and blue for wbCABP-718.

pre-processing pipelines. Although the results were not as impressive as those obtained using the HCP pipeline, the overall trend of the results remained consistent. It is worth noting that this study is primarily based on the HCP pipeline, and we have added a new section to the discussion to clarify this point, in which we discuss the potential impact of different preprocessing choices on the reliability of functional connectivity measures, and the importance of carefully considering these choices in future studies. We will continue to consider the fMRIPrep pipeline in future research, as appropriate. The details of the fMRI-prep analyses are documented below and included as supplementary materials as well as discussed in terms of reproducibility in the revised manuscript.

The figure S43 compares the reliability of graph theory measures and frequency bands using fMRIPrep preprocessing, following the parameterization of the optimal pipeline described in the original manuscript. As in Figure 3a, we decomposed the reliability of different network metrics onto the reliability anatomy plane, which plots reliability as a function of between-subject variability ($V_b$) and within-subject variability ($V_w$). Comparison to the results obtained using the HCP preprocessing pipeline (Figure 5 in the main text) shows that segregation measures have similar levels of reliability for both preprocessing pipelines, while integration measures are significantly lower for the fMRIPrep pipeline. The reliability of different frequency bands onto the reliability anatomy plane shows a trend of increasing reliability from slow-6 to slow-2 bands (Figure 3b), similar to the HCP pipeline. However, slow-1 band exhibits lower reliability, likely due to the partial coverage of the slow-1 frequency range. Overall, the results suggest that fMRIPrep preprocessing produces similar trends in reliability as the HCP pipeline, but with generally lower overall reliability for both network metrics and frequency bands.

We note that the fMRIPrep preprocessing pipeline was performed using *fMRIPrep* 21.0.2 (Esteban,

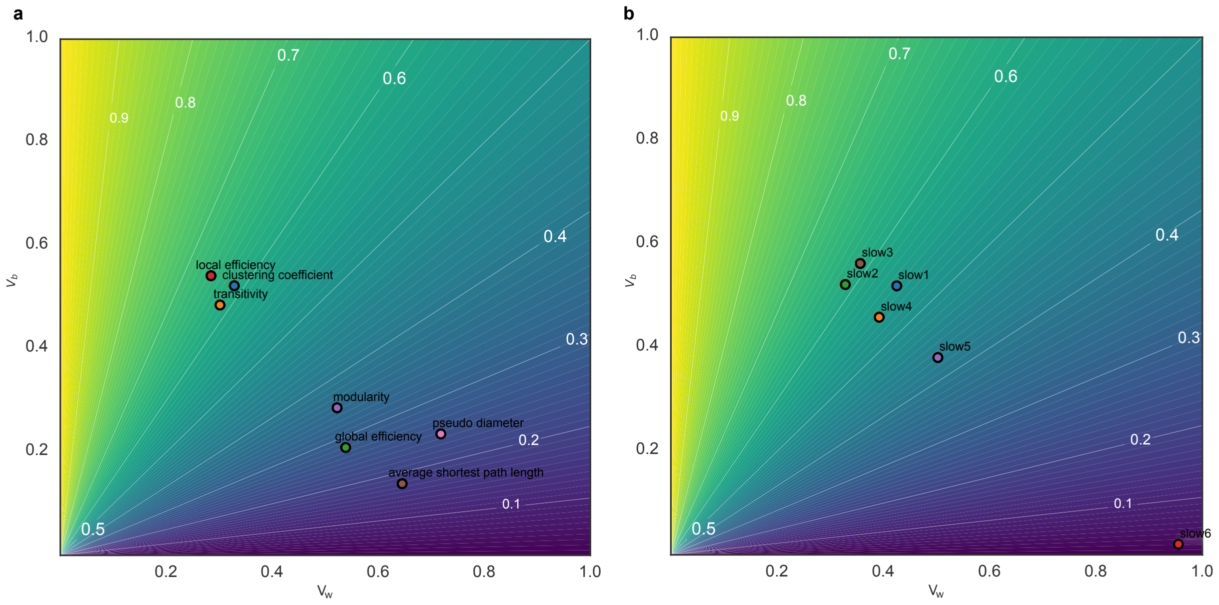

**Fig. S43. Reliability across different network measures and frequency bands using fMRIPrep.** (a) Reliability anatomy of global network measures. The reliability anatomy was plotted as a function of between-subject variability ($V_b$) and within-subject variability ($V_w$). (b) Reliability anatomy of frequency bands.

Markiewicz, et al. (2018); Esteban, Blair, et al. (2018); RRID:SCR_016216), which is based on *Nipype* 1.6.1 (K. Gorgolewski et al. (2011); K. J. Gorgolewski et al. (2018); RRID:SCR_002502). We performed additional standard preprocessing steps on the fmriprep output dtseries data. This included demeaning and detrending the time series, and performing nuisance regression on the minimally preprocessed data. Nuisance regression was based on empirical validation tests by Ciric et al. (2017) to reduce the effects of motion and physiological noise. Specifically, six primary motion parameters were removed, along with their derivatives, and the quadratics of all regressors (24 motion regressors in total). Physiologic noise was modeled using aCompCor on time series extracted from the white matter and ventricles (Behzadi, Restom, Liau, & Liu, 2007a). For aCompCor, the first five principal components from the white matter and ventricles were extracted separately and included in the nuisance regression. In addition, we included the derivatives of each of those components, and the quadratics of all physiological noise regressors (40 physiological noise regressors in total). The nuisance regression model contained a total of 64 nuisance parameters. Note that aCompCor was used in place of global signal regression, given evidence that it has similar benefits as global signal regression for removing artifacts (Power et al., 2018) but without regressing gray matter signals (mixed with other gray matter signals) from themselves, which may result in false correlations (Murphy, Birn, Handwerker, Jones, & Bandettini, 2009; Power, Laumann, Plitt, Martin, & Petersen, 2017).

Then, the cleaned data was fed into our previous ICC optimization pipeline. In order to further verify the results of our previous ICC optimization pipeline, we expanded our analyses to include additional parcellations (MMP-360, wbLGP-458, wbCABP-718) and frequency bands (slow6 to slow1). These additional analyses were performed using the same main calculation conditions as our previous optimal pipeline (wbLGP-458, slow2, OMST, pos, normalization weighted). However, due to considerations of computational complexity and resources, we were unable to completely replicate all of the conditions from our previous analyses and instead only included a subset of the data. As a result, we were unable to generate distribution plots similar to those in our previous analyses, and instead only present discrete points for these additional ICC values. Despite this, we were still able to decompose these ICC values in terms of between- and within-subject variability. Through these additional analyses, we were able to gain some insight into the impact of using the fmriprep preprocessing pipeline on ICC values.

**Preprocessing of B0 inhomogeneity mappings** A total of 3 fieldmaps were found available within the input BIDS structure for this particular subject. A *B0* nonuniformity map (or *fieldmap*) was estimated from the phase-drift map(s) measure with two consecutive GRE (gradient-recalled echo) acquisitions. The corresponding phase-map(s) were phase-unwrapped with `prelude` (FSL 6.0.5.1:57b01774). A *B0*-nonuniformity map (or *fieldmap*) was estimated based on two (or more) echo-planar imaging (EPI) references with `topup` (Andersson, Skare, and Ashburner (2003); FSL 6.0.5.1:57b01774).

**Anatomical data preprocessing** A total of 1 T1-weighted (T1w) images were found within the input BIDS dataset.The T1-weighted (T1w) image was corrected for intensity non-uniformity (INU) with `N4BiasFieldCorrection` (Tustison et al., 2010), distributed with ANTs 2.3.3 (Avants, Epstein, Grossman, & Gee, 2008, RRID:SCR_004757), and used as T1w-reference throughout the workflow. The T1w-reference was then skull-stripped with a *Nipype* implementation of the workflow (from ANTs), using OASIS30ANTs as target template. Brain tissue segmentation of cerebrospinal fluid (CSF), white-matter (WM) and gray-matter (GM) was performed on the brain-extracted T1w using `fast` (FSL 6.0.5.1:57b01774, RRID:SCR_002823, Zhang, Brady, & Smith, 2001). Brain surfaces were reconstructed using `recon-all` (FreeSurfer 6.0.1, RRID:SCR_001847, Dale, Fischl, & Sereno, 1999), and the brain mask estimated previously was refined with a custom variation of the method to reconcile ANTs-derived and FreeSurfer-derived segmentations of the cortical gray-matter of Mindboggle (RRID:SCR_002438, Klein et al., 2017). Volume-based spatial normalization to two standard spaces (MNI152NLin2009cAsym, MNI152NLin6Asym) was performed through nonlinear registration with `antsRegistration` (ANTs 2.3.3), using brain-extracted versions of both T1w reference and the T1w template. The following templates were selected for spatial normalization: *ICBM 152 Nonlinear Asymmetrical template version 2009c* [Fonov, Evans, McKinstry, Almli, and Collins (2009), RRID:SCR_008796; TemplateFlow ID: MNI152NLin2009cAsym], *FSL's MNI ICBM 152 non-linear 6th Generation Asymmetric Average Brain Stereotaxic Registration Model* [Evans, Janke, Collins, and Baillet (2012), RRID:SCR_002823; TemplateFlow ID: MNI152NLin6Asym].

**Functional data preprocessing** For each of the 4 BOLD runs found per subject (across all tasks and sessions), the following preprocessing was performed. First, a reference volume and its skull-stripped version were generated by aligning and averaging 1 single-band references (SBRefs). Head-motion parameters with respect to the BOLD reference (transformation matrices, and six corresponding rotation and translation parameters) are estimated before any spatiotemporal filtering using `mcflirt` (FSL 6.0.5.1:57b01774, Jenkinson, Bannister, Brady, & Smith, 2002). The estimated *fieldmap* was then aligned with rigid-registration to the target EPI (echo-planar imaging) reference run. The field coefficients were mapped on to the reference EPI using the transform. The BOLD reference was then co-registered to the T1w reference using `bbregister` (FreeSurfer) which implements boundary-based registration (Greve & Fischl, 2009). Co-registration was configured with six degrees of freedom. First, a reference volume and its skull-stripped version were generated using a custom methodology of *fMRIPrep*. Several confounding time-series were calculated based on the *preprocessed BOLD*: framewise displacement (FD), DVARS and three region-wise global signals. FD was computed using two formulations following Power (absolute sum of relative motions, Power et al. (2014)) and Jenkinson (relative root mean square displacement between affines, Jenkinson et al. (2002)). FD and DVARS are calculated for each functional run, both using their implementations in *Nipype* (following the definitions by Power et al., 2014). The three global signals are extracted within the CSF, the WM, and the whole-brain masks. Additionally, a set of physiological regressors were extracted to allow for component-based noise correction (*CompCor*, Behzadi, Restom, Liau, & Liu, 2007b). Principal components are estimated after high-pass filtering the *preprocessed BOLD* time-series (using a discrete cosine filter with 128s cut-off) for the two *CompCor* variants: temporal (tCompCor) and anatomical (aCompCor). tCompCor components are then calculated from the top 2% variable voxels within the brain mask. For aCompCor, three probabilistic masks (CSF, WM and combined CSF+WM) are generated in anatomical space. The implementation differs from that of Behzadi et al. in that instead of eroding the masks by 2 pixels on BOLD space, the aCompCor masks are subtracted a mask of pixels that likely contain a volume fraction of GM. This mask is obtained by dilating a GM mask extracted from the FreeSurfer's *aseg* segmentation, and it ensures components are not extracted from voxels containing a minimal fraction of GM. Finally, these masks are resampled into BOLD space and binarized by thresholding at 0.99 (as in the original implementation). Components are also calculated separately within the WM and CSF masks. For each CompCor decomposition, the *k* components with the largest singular values are retained, such that the retained components' time series are sufficient to explain 50 percent of variance across the nuisance mask (CSF, WM, combined, or temporal). The remaining components are dropped from consideration. The head-motion estimates calculated in the correction step were also placed within the corresponding confounds file. The confound time series derived from head motion estimates and global signals were expanded with the inclusion of temporal derivatives and quadratic terms for each (Satterthwaite et al., 2013). Frames that exceeded a threshold of 0.5 mm FD or 1.5 standardised DVARS were annotated as motion outliers. The BOLD time-series were resampled into standard space, generating a *preprocessed BOLD run in MNI152NLin2009cAsym space*. First, a reference volume and its skull-stripped version were generated using a custom methodology of *fMRIPrep*. The BOLD time-series were resampled onto the following surfaces (FreeSurfer reconstruction nomenclature): *fsaverage*. Automatic removal of motion artifacts using independent component analysis (ICA-AROMA, Pruim et al., 2015) was performed on the *preprocessed BOLD on MNI space* time-series after removal of non-steady state volumes and spatial smoothing with an isotropic, Gaussian kernel of 6mm FWHM (full-width half-maximum). Corresponding "non-aggresively" denoised runs were produced after such smoothing. Additionally, the "aggressive" noise-regressors were collected and placed in the corresponding confounds file. *Grayordinates* files (Glasser et al., 2013) containing 91k samples were also generated using the highest-resolution `fsaverage` as intermediate standardized surface space. All resamplings can be performed with *a single interpolation step* by composing all the pertinent transformations (i.e. head-motion transform matrices, susceptibility distortion correction when available, and co-registrations to anatomical and output spaces). Gridded (volumetric) resamplings were performed using `antsApplyTransforms` (ANTs), configured with Lanczos interpolation to minimize the smoothing effects of other kernels (Lanczos, 1964). Non-gridded (surface) resamplings were performed using `mri_vol2surf` (FreeSurfer). Many internal operations of *fMRIPrep* use *Nilearn* 0.8.1 (Abraham et al., 2014, RRID:SCR_001362), mostly within the functional processing workflow. For more details of the pipeline, see the section corresponding to workflows in *fMRIPrep*'s documentation.
